# A general framework for modeling pathogen transmission in co-roosting host communities

**DOI:** 10.1101/2023.11.21.568148

**Authors:** Molly C. Simonis, Daniel J. Becker

## Abstract

Cross-species transmission of pathogens can be facilitated by frequent contact among wildlife. Cross-species transmission is often driven by phylogenetic similarity between host species, but the role this plays when multiple host species co-roost is unknown. We developed a generalizable framework for understanding how cross-species transmission is driven by contact among co-roosting species spanning evolutionary similarities and the net impact on roost-level infection prevalence. We developed ordinary differential equation models describing population and infection dynamics between two and three co-roosting species. We parameterized models using co-roosting Neotropical bat systems, with interspecific transmission exponentially declining with phylogenetic distance. To assess the relative contribution of contact rates and phylogenetic similarity, we co-varied intraspecific transmission rates and phylogenetic distances while considering sensitivity to host and pathogen traits. While our models converged on similar equilibria under high intraspecific transmission or long durations of infection and immunity or latency, simulations with lower intraspecific transmission and shorter such periods revealed roost-level prevalence was greatest when hosts were most closely related. However, we identified regions of parameter space where roost-level prevalence also maximized when hosts were distantly related, driven by species-specific traits. Our generalizable models are adaptable to other co-roosting systems and informs our understanding of pathogen spillover.

## Introduction

Zoonotic pathogens, which spillover from wildlife to humans, are one of the largest threats to human health [1,2]. In recent years, we have seen multiple deadly epidemics (and at least two pandemics) of emerging infectious diseases that were a result of zoonotic spillover (e.g., Nipah virus, Ebola virus, Hendra virus, Marburg virus, H1N1 influenza virus, MERS-CoV, SARS-CoV, and SARS-CoV-2 [3]). Inter-species contact and susceptibility to infection are required for cross-species transmission, with the latter formed by coevolutionary immunological barriers between hosts and their pathogens [4]. As such, cross-species transmission is more common for closely related host species [5–12]. However, the role phylogenetic similarity plays when multiple host species have regular and epidemiologically relevant contacts, such as occurs within shared roosts, is less understood.

Co-roosting (i.e. multiple species sharing a roost) is common across wildlife taxa and can facilitate cross-species transmission due to close interactions between hosts [13,14]. There are many examples of co-roosting in natural and anthropogenic systems (Table 1). Many meso- and top carnivores are known to share dens [15–17], and multiple bird species can share natural and anthropogenic cavities [18–21]. Even though co-roosting and overall gregariousness are strong predictors of viral sharing among species at macroecological scales [11,22], few studies present co-roosting as facilitating pathogen transmission (but see [23,24]). However, growing concerns of cross-species transmission in light of the recent SARS-CoV-2 pandemic have made natural history reporting of co-roosting, such as has been naturally observed between pangolins and bats [25], even more important.

**Table 1.** Examples of co-roosting multi-species host systems, including pathogens assessed.

| Roosting Habitat | Co-roosting Species | Pathogen of Concern<br>Mentioned or Tested | Reference |
| --- | --- | --- | --- |
| underground burrows | giant pangolin (*Smutsia gigantea*)<br>white-bellied pangolin (*Phataginus tricuspis*)<br>Old World leaf-nosed bat *sp.* (*Hipposideridae sp.*)<br>sac-winged bat *sp.* (*Emballonuridae sp.*)<br>bent-winged bat *sp.* (*Miniopterus sp.*)<br>other unknown bat *sp.* | SARS-CoV-2 | [25] |
| chimney | chimney swift (*Chaetura pelagica*)<br>rock pigeon (*Columba livia*) |  | [21] |
| nest (brood parasitism) | brown-headed cowbirds (*Molothrus ater*)<br>common grackle (*Quiscalus quiscula*) |  | [19] |
| nest box (nest predation) | house wren (*Troglodytes aedon*)<br>tree swallow (*Tachycineta bicolor*) |  | [18] |
| cave | common vampire bat (*Desmodus rotundus*)<br>Jamaican fruit bat (*Artibeus jamaicensis*)<br>Seba's short-tailed bat (*Carollia perspicillata*)<br>hairy-legged vampire bat (*Diphylla ecaudata*)<br>little big-eared bat (*Micronycteris megalotis*)<br>cave myotis (*Myotis velifer*)<br>Mexican funnel-eared bat (*Natalus stramineus*)<br>hairy-legged myotis (*Myotis keaysi*)<br>fringed myotis (*Myotis thysanodes*) |  | [30] |
| hollow tree cavity or underground tunnel | common vampire bat (*Desmodus rotundus*)<br>Pallas's long-tongued bat (*Glossophaga soricina*)<br>greater sac-winged bat (*Saccopteryx bilineata*)<br>fringe-lipped bat (*Trachops cirrhosus*)<br>big-eared woolly bat (*Chrotopterus auritus*) | *Bartonella spp.*;<br>haemosplasmas | [36,37] |
| artificial roost in flight enclosure | common vampire bat (*Desmodus rotundus*)<br>pale spear-nosed bat (*Phyllostomus discolor*)<br>little yellow-shouldered bat (*Sturnira lilium*) |  | [48] |
| abandoned mines | little brown bat (*Myotis lucifugus*)<br>northern long-eared bat (*Myotis septentrionalis*)<br>tri-colored bat (*Perimyotis subflavus*)<br>big brown bat (*Eptesicus fuscus*) | *Pseudogymnoascus destructans* | [23] |
| trees | ornate flying fox (*Pteropus ornatus*)<br>Pacific flying fox (*Pteropus tonganus*) | *Leptospira* spp. | [24] |
| rocky outcrop den | raccoon (*Procyon lotor*)<br>Virginia opossum (*Didelphis virginiana*)<br>striped skunk (*Mephitis mephitis*) |  | [15] |
| tree cavity | helmeted woodpecker (*Celeus galeatus*)<br>white-throated woodcreeper (*Xiphocolaptes albicollis*) |  | [20] |
| underground burrow | red fox (*Vulpes vulpes*)<br>raccoon dog (*Nyctereutes procyonoides*)<br>Japanese badger (*Meles anakuma*)<br>Japanese weasel (*Mustela itatsi*)<br>Japanese hare (*Lepus brachyurus*)<br>unclassified rodent and bat species |  | [17] |
| underground den | Arctic fox (*Vulpes lagopus*)<br>wolf (*Canis lupus*) |  | [16] |
| underground burrow | European badger (*Meles meles*)<br>crested porcupine (*Hystrix cristata*)<br>red fox (*Vulpes vulpes*)<br>pine marten (*Martes martes*)<br>stone marten (*Martes foina*)<br>eastern cottontail (*Sylvilagus floridanus*)<br>wood mouse (*Apodemus* sp.)<br>brown rat (*Rattus norvegicus*)<br>coypu (*Myocastor coypus*) |  | [71] |
| sugarcane field | white-tailed kite (*Elanus leucurus*)<br>merlin (*Falco columbarius*)<br>northern harriers (*Circus cyaneus*)<br>American kestrel (*Falco sparverius*)<br>white-tailed hawk (*Buteo albicaudatus*)<br>crested caracara (*Caracara cheriway*) |  | [72] |

Theoretical models offer an important tool to reconcile evolutionary barriers to cross-species transmission with the frequent contacts afforded by co-roosting behavior. Despite prior calls for developing mathematical frameworks to study pathogen spread within multi-host communities [26–28], we lack generalizable models that explicitly and flexibly consider phylogenetic barriers. Previous theory has considered how proportional reductions in transmission efficiency between two hosts affects pathogen spillover [29], and such work can be generalized and extended to consider the widely observed decline in cross-species transmission with phylogenetic distance in multi-host communities more generally..

Here, we develop generalizable models for exploring cross-species transmission in multi-species, co-roosting systems. We adapt common epidemiological modeling frameworks for pathogen transmission between two and three host species, incorporating greater transmission efficiency among closely related host species. To explore model behavior, we guided parameterization with Neotropical bat systems, which commonly display co-roosting behavior and harbor a wide range of zoonotic pathogens [30–32]. Our results support realistic contexts for cross-species pathogen transmission in two ways. First, infection prevalence across the co-roosting community most maximizes when host species are closely related, supporting positive relationships between cross-species transmission and phylogenetic relatedness. Secondly, roost-level infection prevalence infrequently maximizes when hosts are distantly related, supporting increased pathogen spillover when hosts are regularly in close contact. Our modeling framework presented here can be modified and applied across many multi-host systems where transmission requires close contact.

## Methods

To determine how variable phylogenetic relatedness influences pathogen prevalence in a co-roosting host community, we developed multiple sets of ordinary differential equation models describing population and infection dynamics. We coarsely parameterized these models around well-studied bat systems, but our model structures are sufficiently generalizable to apply to the broad range of systems in which co-roosting of host species and pathogen sharing can occur (Table 1).

### Infection dynamics of two co-roosting hosts

We first consider the population dynamics of two host species occupying a shared roost environment, with two distinct population sizes (*N_A_* and *N_B_*). Per-capita reproduction for both species is described by *b_0_* − *b_1_N*, where *b_0_*and *b_1_* are the density-independent and density-dependent birth rates and *N* = *N_A_* + *N_B_*, such that density dependence is driven by joint species occupancy. Both species experience per-capita mortality at rate *μ_i_*, where subscripts allow for species-specific lifespans. Natural mortality rates (*μ*) are unique to each host species based on average lifespans (1 / lifespan). The coupled population dynamics of the roost are thus defined by the following differential equations:

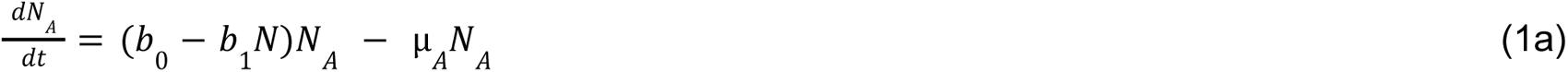

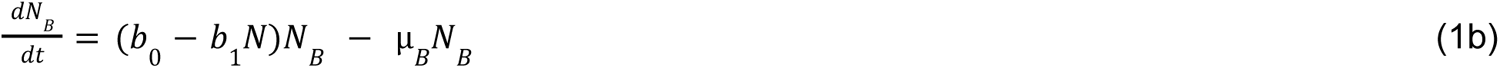

We next expand single-species epidemiological models classifying hosts as susceptible (*S*), infected (*I*), and recovered (*R*) or latent (*L*), where the population size of each species is represented by *N_i_* = *S_i_* + *I_i_* + *R_i_*| *L_i_* [33,34]. Pathogen transmission from infected to susceptible hosts within each species occurs at the rate β, and we assume equivalent intra-specific transmission rates among host species alongside density-dependent contacts. Because the probability of cross-species transmission typically declines for distantly related species across systems [5–10,12], we model inter-specific transmission (θ) as an exponential function of host phylogenetic distance. Here, ψ represents proportional phylogenetic similarity and λ is a shape parameter governing the strength of exponential decay (Fig. S1)[35]. As our model always considers distinct host species, inter-specific transmission is always some fraction of intra-specific transmission, and subscripts refer to transmission from the donor host species (*i*) to the recipient host species (*j*):

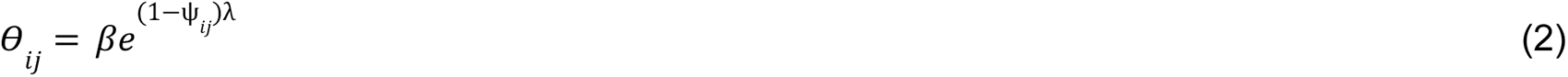

Following transmission, infectious hosts become recovered or latently infected at rate γ (i.e., 1/γ provides the infectious period). We thus consider a susceptible–infected–recovered–susceptible (SIRS) system and a susceptible–infected–latent–infected (SILI) system (Fig. 1). Here, the rate ϵ governs the loss of protective immunity (SIRS model) or reactivation of latent (or chronic) infection (SILI model). Such frameworks are applicable to many pathogens that require close-contact transmission, are temporarily immunizing, and exhibit general chronic infection or true latency [35]. We do not consider disease-induced mortality, such that our models are most applicable to low-virulence pathogens (e.g., those with strict coevolutionary history with a particular host clade). We therefore describe SIRS dynamics by the following equations:

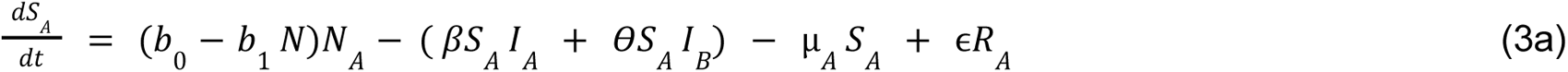

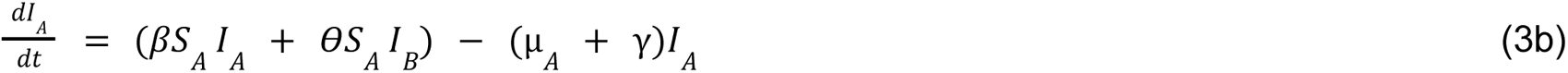

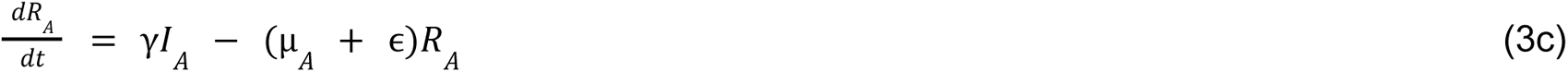

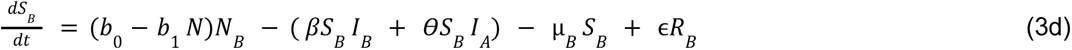

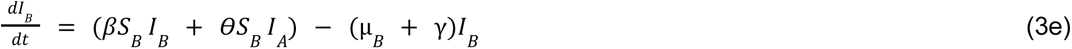

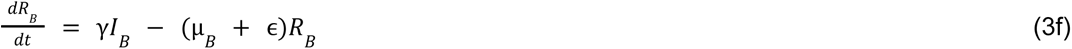

**Figure 1.**
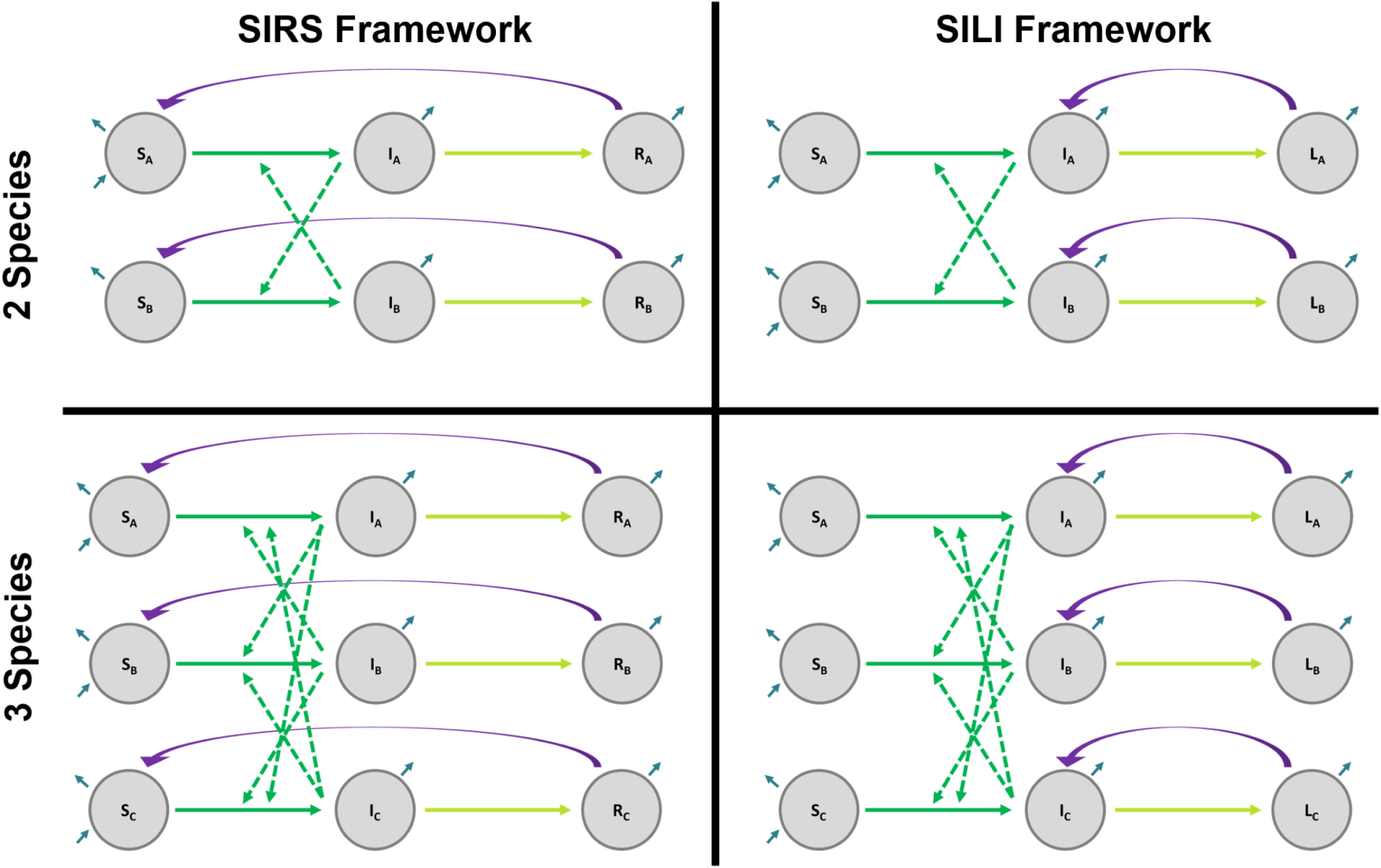
We created two- and three-species compartmental models using SIRS and SILI frameworks. Colors represent processes for pathogen transmission (green), pathogen clearance (yellow), waning immunity or reactivation of latent infection (purple), and demography (blue).

Similarly, SILI dynamics are given by the following equations:

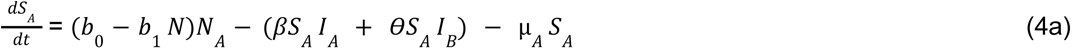

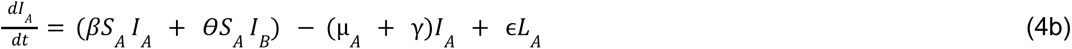

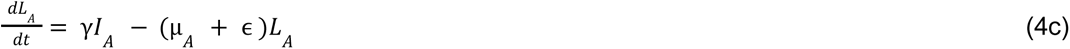

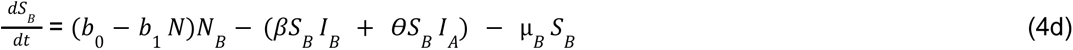

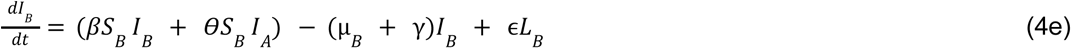

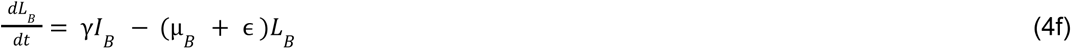

### Infection dynamics of three co-roosting hosts

We next expand equations 3–4 to consider three host species sharing a single roost. We again first consider population dynamics, now of three host species (*N_A_*, *N_B_* and *N_C_*), such that *N* = *N_A_* + *N_B_* + *N_C_*. Similar to two-species models, natural mortality rates (μ) are unique to each species. As such, the strength of density dependence in birth is now shaped by occupancy of all three species. We therefore describe SIRS dynamics in the three-species co-roosting system by the following equations:

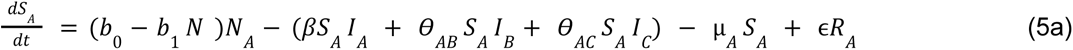

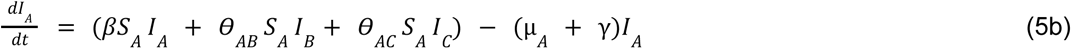

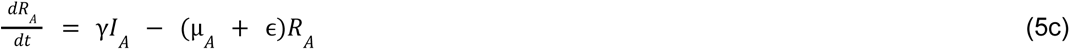

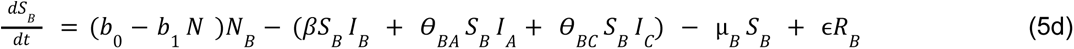

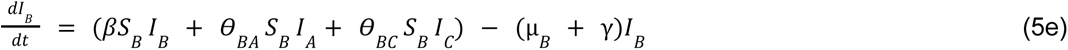

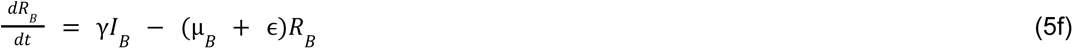

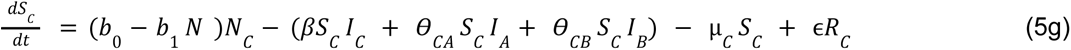

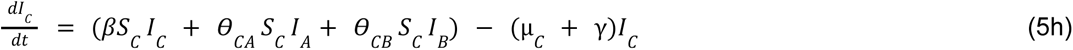

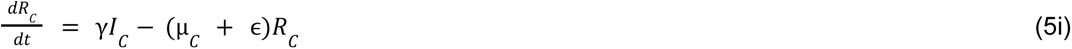

Similarly, SILI dynamics under three co-roosting species is given by the following equations:

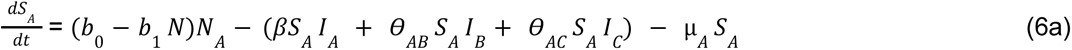

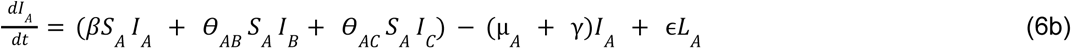

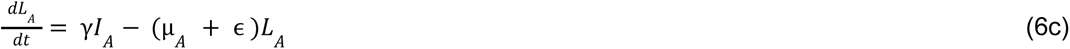

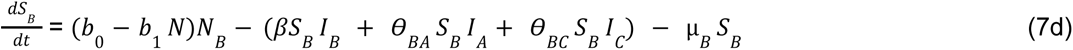

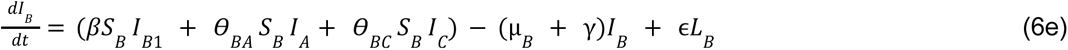

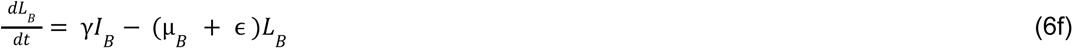

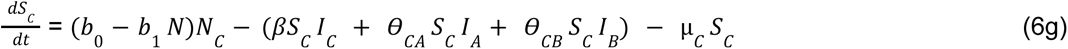

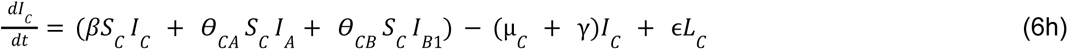

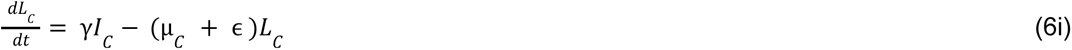

### Model parameterization

We parameterized our models around co-roosting bat species in Central and South America. We considered common vampire bats (Phyllostomidae: *Desmodus rotundus*) as the focal host species, as they commonly co-roost with different bat species that span closely and distantly related families (Table 1). For example, in Costa Rica, vampire bats roost in caves with bat species from at least three families (e.g., Vespertilionidae: hairy-legged myotis (*Myotis keaysi*), fringed myotis (*Myotis thysanodes)*, cave myotis (*Myotis velifer*); Natalidae: Mexican funnel-eared bat (*Natalus stramineus*); Phyllostomidae: Jamaican fruit bat (*Artibeus jamaicensis*), Seba’s short-tailed bat (*Carollia perspicillata*), little big-eared bat (*Micronycteris megalotis*)) and even other vampire bat species (Table 1)[30]. In Belize, common vampire bats roost in hollow trees or abandoned temples with bat species from at least two families (Phyllostomidae: Pallas’s long-tongued bat (*Glossophaga soricina*), fringe-lipped bat (*Trachops cirrhosus*), big-eared woolly bat (*Chrotopterus auritus*); Emballonuridae: greater sac-winged bat (*Saccopteryx bilineata*); Table 1)[36,37]. As a highly social species, vampire bats have fine-scale connectivity networks within their colonies [38], but also share pathogens between co-roosting bat species as well as other non-bat wildlife, livestock, and humans due to their unique diet of blood [36,37,39–42]. Within the context of this system, we parameterized infection processes around coronaviruses and herpesviruses, which circulate in diverse Neotropical bat species (including common vampire bats) and exhibit SIRS and SILI dynamics, respectively [43–46].

We similarly parameterized host demographics around these bat systems. Bats are unique, with relatively long lifespans and low reproductive rates compared to other small mammals, such as rodents [47]. We chose a maximum per-capita birth rate (*b_0_*) for all species to be two births each year, which is the maximum annual fecundity of most Neotropical bat species (Table 2)[47]. As our default parameterization, we assumed an equal starting population size among all host species (*N_A_ = N_B_, N_A_ = N_B_ = N_C_*), although we assessed the sensitivity of this assumption to numerical dominance of vampire bats [48]. Under our default assumption, we initialized our models with each species starting with 150 bats, representing a moderate-to-large colony of vampire bats [7,49,50]. The total population in three-species models when all population sizes were equal (450 bats) also denoted our carrying capacity (*k*) (table 2). When vampire bat populations (*N_A_*) were larger than other species (*N_B_* or *N_C_*), we halved the starting populations of other species within our two- and three-species models (Table 2).

**Table 2.** Descriptions of parameters and parameter space used in two- and three-species SIRS and SILI models. All units are in days unless otherwise noted. Colors represent processes for pathogen transmission (green), pathogen clearance (yellow), waning immunity or latency (purple), and demography (blue), as also visualized in Figure 1.

| Process | Parameter | Description | Parameter Space | Citation |
| --- | --- | --- | --- | --- |
| transmission | $\beta$ | intraspecific transmission rate | 0.0005, 0.0025, 0.005, 0.0075 | [51] |
| | $\theta$ | interspecific transmission rate | $\beta e^{(1-\Psi)\lambda}$ | Fig. S1; [12,73] |
| | $\lambda$ | exponential decay curve shape parameter | - 6 | Fig. S1 |
| | $\Psi$ | phylogenetic relatedness | 0.0001-0.9999 | Fig. 2; [74] |
| pathogen clearance | $\gamma$ | infection clearance rate | short: 1/3, 1/5, 1/7<br>long: 1/365, 1/548, 1/730 | [44,51] |
| waning immunity OR latency | $\varepsilon$ | waning immunity OR latency rate | short: 1/30, 1/60, 1/90<br>long: 1/1095, 1/1643, 1/2190 | [44,51,75] |
| demography | $\mu_A$ | natural death rate of vampire bats | 1/(6 * 365) | [76] |
| | $\mu_B$ | natural death rate of second bat species | 1/(9*365) | |
| | $\mu_C$ | natural death rate of third bat species | 1/(15*365) | |
| | $b_0$ | Maximum per-capita birth rate | 2 bats/365 | [47] |
| | $b_1$ | density dependent per-capita birth rate | $(b_0/k)$ | |
| | $k$ | carrying capacity | 450 bats | |
| | $N_A$ | population size of vampire bats<br>($S_A + I_A + R_A$) | starting size: 150 | [7,49,50] |
| | $N_B$ | population size of second bat species | starting size: 75, 150 | |
| | | $(S_B + I_B + R_B)$ | | |
| | $N_C$ | population size of third bat species<br>$(S_C + I_C + R_C)$ | starting size: 38, 75, 150 | |
| | $N$ | total population size | $N_A + N_B$ OR<br>$N_A + N_B + N_C$ | |

We set natural mortality rates (*μ*) in our two- and three-species models based on average Neotropical bat lifespans. For example, common vampire bats live around six years in the wild, while other known co-roosting bat species can live a maximum of six to 20 years in the wild (e.g. sac-winged bat, *Saccopteryx bilineata* = 6 years maximum wild lifespan; hairy-legged vampire bat, *Diphylla ecaudata* = 8 years maximum wild lifespan; pale spear-nosed bat, *Phyllostomus discolor* = 9 years maximum wild lifespan; cave myotis, *Myotis velifer* = 11.3 years maximum wild lifespan; fringed myotis, *Myotis thysanodes* = 18.3 years maximum wild lifespan). Therefore, we parameterized mortality rates based on lifespans of common vampire bats in the wild (six years) as species A, and chose a nine (species B) or 15 year lifespan (species C) for the other host species within our two- and three-species models.

Lastly, we parameterized pathogen traits based on prior studies of bat–virus interactions. For intraspecific transmission rates (β), we used estimates for alphacoronavirus infection in Australian southern myotis (*Myotis macropus*) (Table 2)[51]. We also systematically explored lower intraspecific transmission rates to consider cases of lesser social connectivity. Pathogen clearance rates (γ) and rates of waning immunity or infection reactivation (ϵ) were parameterized under short or long conditions, since these processes remain poorly understood in bats. Short parameterizations represent coronavirus dynamics and were chosen to fall within previously published rates for transient alphacoronavirus infections in Australian southern myotis (Table 2)[51]. Long parameterizations represent herpesvirus dynamics and were selected from maximum likelihood estimates of clearance and reactivation rates in vampire bats (Table 2)[44].

### Modeling Approach

To assess the contribution of host evolutionary history to infection dynamics in the context of co-roosting, we systematically covaried phylogenetic relatedness between paired hosts (ѱ; Fig. 2) and intraspecific transmission (β) for both SIRS and SILI frameworks, first for two-species models and then for three-species models. We also considered the sensitivity of our results to host and pathogen traits. Specifically, we simulated models across all factorial combinations of host population sizes (two-species models: *N_A_* = *N_B_*, *N_A_*> *N_B_*; three-species models: *N_A_* = *N_B_*= *N_C_*, *N_A_* > *N_B_* > *N_C_*, *N_A_* > *N_B_* < *N_C_*), durations of infection (1/γ), and durations of temporary immunity or latency (1/*ϵ*). For pathogen traits, we paired short infectious periods with short durations of immunity or latency (i.e., coronaviruses) and long infectious periods with long durations of immunity or latency (i.e., herpesviruses). As such, each two- and three-species model was simulated across 4,536 and 2,000,376 unique parameter combinations, respectively.

**Figure 2.**
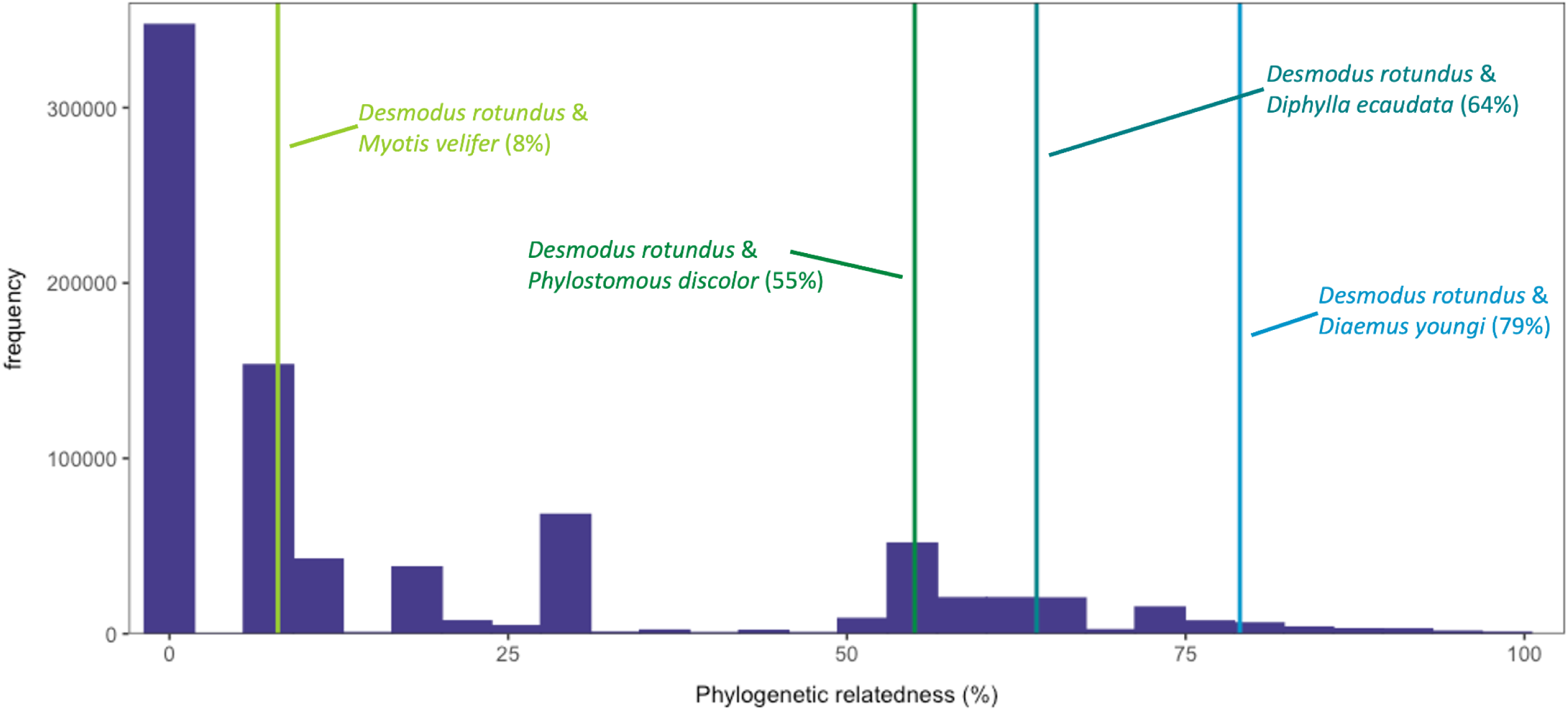
Distribution of phylogenetic relatedness among bats using the most recent mammalian phylogeny [74]. Vertical lines represent specific examples of phylogenetic relatedness between *Desmodus rotundus* and other Neotropical bat species, as we tested model sensitivity by focusing on parameterization for *D. rotundus* and potential co-roosting bat species.

We simulated each model until equilibrium, using a simulation length of 20 years to ensure each host species had a minimum two generations. All models reached equilibrium under seven years (Fig. S2-S5). Per each parameterization, we derived equilibrium roost-level pathogen prevalence (two species: (*I_A_* + *I_B_*) / *N*; three species: (*I_A_* + *I_B_* + *I_C_*) / *N*). All simulations were performed in the statistical environment R version 4.2.1 [52] using the *deSolve* package [53]. Simulations were performed remotely through the University of Oklahoma’s Supercomputing Center for Education & Research. Results were visualized using the *ggplot2* package in R version 4.3.1 [52,54].

## Results

In general, two- and three-species SIRS and SILI models yielded qualitatively similar patterns. Roost-level pathogen prevalence at equilibrium (hereafter “roost prevalence”) increasingly maximized with longer infectious periods and shorter durations of immunity or latency (Figs. 3, S6-61). For all SILI models and SIRS models with long infectious and immune or latent periods, roost prevalence reached the same equilibrium regardless of intraspecific transmission and phylogenetic relatedness in either two- or three-species models (Figs. S3, S9-S13, S15, S17, S19, S21, S23, S25, S27, S29, S31, S33, S35, S37-S61). Similar patterns were observed for SIRS models subject to short infectious and immune periods when intraspecific transmission was relatively large (β > 0.0025; Figs. S18, S20, S26, S28, S34, S36).

**Figure 3.**
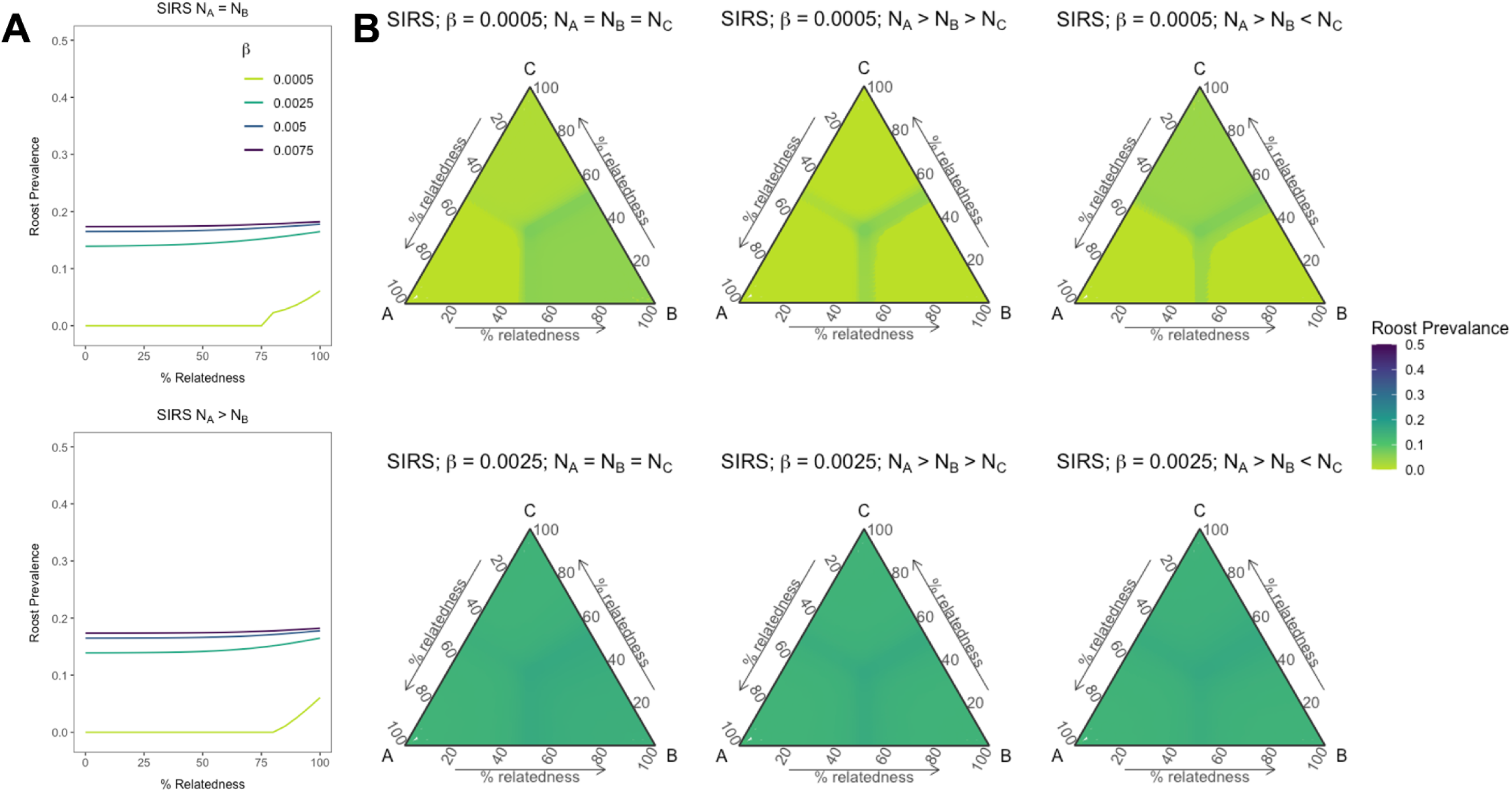
Roost prevalence at equilibrium for (**A**) two-species and (**B**) three-species SIRS models under short infectious periods (seven days) and immune durations (30 days). **A**) Roost prevalence increases with intraspecific transmission and varies with phylogenetic relatedness between two species when starting populations of co-roosting species are equal (top) or when one co-roosting species dominates the starting population size (bottom). Colors represent intraspecific transmission (β). **B**) Roost prevalence increases with β (rows, top to bottom) and varies with pairwise phylogenetic relatedness between all three species under different starting population sizes of co-roosting species (columns). Species become more closely and evenly related moving toward the center of the ternary plots and thus become more distantly and unevenly related moving toward the plot edges. Colors represent roost prevalence at equilibrium. All simulations are based on short infectious and immune periods (γ = 1/7, ε = 1/30).

When SIRS models were parameterized by short infectious and immune periods as well as low-to-moderate intraspecific transmission (β = 0.0005 and 0.0025), we found effects of phylogenetic relatedness on pathogen invasion within co-roosting communities and subsequent pathogen persistence. Specifically, we identified sharp inflections of roost prevalence as a function of phylogenetic relatedness (Figs. 3, S6, S8, S14, S16, S22, S24, S30, S32). The strongest inflections between pathogen extinction and invasion with phylogenetic relatedness were observed at the lowest transmission rates (Figs. 3, S6, S14, S22, S30). In two-species models with lowest intraspecific transmission, pathogen invasion could not occur unless co-roosting species were more than 75% related (Figs. 3A, S6, S8. In three-species models, phylogenetic boundaries in roost prevalence were further dependent on the starting populations of co-roosting species and their species-specific lifespans and thus, mortality rates. While roost prevalence was maximized when all three species were closely related (i.e., the center of ternary plots; Fig. 3B), that maximal prevalence was also expanded across phylogenetic space driven by species B and C, given their longer lifespans than species A (Figs. 3B, S14, S22, S30). Therefore, regardless of starting host population size within a roost, co-roosting population sizes of species B and C were greater than species A by equilibrium, producing more infected individuals and contributing distinct increases to roost prevalence within phylogenetic space (Figs. 3B, S14, S22, S30).

## Discussion

Co-roosting between multiple host species can facilitate cross-species transmission due to frequent interactions between hosts [13]. Here, we created a generalizable model framework for understanding how phylogenetic relatedness influences host pathogen prevalence in the context of this co-roosting environment, considering pathogens requiring close contact for transmission. When parameterizing two- and three-species models for co-roosting Neotropical bat hosts, we found roost prevalence at equilibrium generally was greatest when hosts were more closely related, given relatively lower baseline levels of intraspecific transmission. However, our models also created sharp phylogenetic boundaries between pathogen extinction and invasion, signifying species-specific drivers of equilibrium infection prevalence in a multi-host community that does not necessarily align with expectations of increasing prevalence with phylogenetic relatedness [5–10,12].

Phylogenetic relatedness between host species has long been observed to be a strong driver of cross-species transmission [5–10,12]. In SIRS models with low-to-moderate intraspecific transmission and short infectious and immune periods, our models provide further theoretical support for these findings. However, we also found specific contexts in which roost prevalence departed from this assumption. In SIRS models with low-to-moderate intraspecific transmission and short infectious and immune periods, we found roost prevalence to be maximized when species were distantly and unevenly related. While these results are seemingly contradictory with our understanding of phylogenetic barriers to cross-species transmission, such risks also increase with overlapping geography and decrease with host species dispersal [5,7,12,22]. Further, frequent interactions between susceptible and infected hosts can facilitate interspecific transmission [13]. For example, geographic overlap among host species more frequently resulted in coronavirus sharing at global scales than phylogenetic relationships [55]. In our models, we considered a single community of co-roosting host species and did not account for immigration or emigration, representing co-roosting as a strong overlap in space with shared effects on population growth. Therefore, contexts in which roost-level prevalence was highest between species more distantly and unevenly related suggests that extreme spatial overlap in host communities (i.e., co-roosting) may sometimes disrupt normal coevolutionary patterns between host species and their pathogens, even at low levels of intraspecific transmission and short infectious and immune periods.

Gregariousness has previously been shown to be a strong predictor of cross-species transmission at macroecological scales across taxa and in bats specifically, which is relevant given our explicit model parameterization [22,56,57]. Past theory in single-host systems suggests infection prevalence quickly saturates following an epidemic for large communal roosts [58]. Here, starting populations of co-roosting species and our selected carrying capacity represent a moderate-to-large community roost, and we also found our models to reach equilibrium rapidly, with similar equilibria reached across a large region of parameter space when intraspecific transmission was high and durations of infection and immunity or latency were long. Yet distinct from prior single-host theory, both two- and three-species SIRS models with lower intraspecific transmission and shorter infectious and immune periods identified sharp phylogenetic boundaries where the pathogen can invade. Within these simulations, these phylogenetic boundaries differed between each of our starting host population size scenarios. Therefore, relative abundance of co-roosting hosts may drive the strong inflection points between pathogen extinction and invasion, especially when intraspecific transmission rates are low. While our population size parameterizations are likely applicable to a wide range of co-roosting systems (Table 1), future applications of our model to larger co-roosting populations (e.g., cave bats in Mexico with greater than 1,000 individuals [59]) could find weaker effects of phylogenetic similarity depending on the numerical dominance of each host species.

Species-specific traits influenced transmission dynamics and roost-level pathogen prevalence in addition to multi-species population densities. In our three-species SIRS models with phylogenetic boundaries between pathogen extinction and invasion, infection plateaus occurred when all three species were closely and evenly related. However, species-specific traits influenced phylogenetic boundaries of pathogen extinction and invasion as well as long-term persistence. Each species in our models had distinct life spans that were used to estimate mortality rates. Species A had the shortest lifespan (six years) followed by species B (nine years) and C (15 years). Theory predicts longevity of a species to increase the prevalence of infection in a host population [60], translated here as more opportunity to gain new infections or experience relapse of chronic infection. Because species C and B lived longer than species A, species B and C obtained greater species-specific infection prevalence and dominated transmission, driving pathogen invasion across phylogenetic space. Indeed, within all possible regions of phylogenetic relatedness space where the pathogen invaded, species B and C had larger population sizes of infected individuals at equilibrium than species A. Additionally, because species B and C have lower mortality rates than species A, they obtained higher densities by equilibrium regardless of starting population sizes.

SIRS and SILI frameworks have been well-characterized to generate distinct infection dynamics [34,61]. While we likewise observed differing dynamics at the start of epidemics (Figs. S2-S5), we observed similar equilibria within each model family under long infectious and immune or latency periods. This is likely due to infections moving more rapidly through the co-roosting host system and maintaining a steady influx of infected individuals following an epidemic. For SILI models, chronic infections keep infected individuals within the co-roosting system for longer, allowing for overall high roost-level infection prevalence at equilibrium. Finally, for SIRS models under short infectious and immune periods, even moderate increases to intraspecific transmission homogenized equilibria throughout phylogenetic space.

We here contribute to growing efforts to better understand cross-species transmission, particularly in relation to evolutionary barriers. A large body of previous theory has considered pathogen dynamics in multi-host systems [28,29,62–66], but the evolutionary underpinnings of cross-species transmission remain poorly integrated into models. Recent work has explicitly modeled interspecific transmission as a proportional reduction of intraspecific transmission [29], and we expanded this approach to consider the well-studied functional relationship (i.e., exponential decay) between phylogenetic distance and probability or frequency of cross-species transmission [5–12]. These models can serve as a starting point for future theory that may further expand upon the evolutionary barriers to transmission. Integrating within- and between-host social dynamics [56,67] into our phylogenetic framework here could improve understanding species-specific drivers of roost prevalence in co-roosting environments. Further, simulating these models across empirical multi-species contact networks would provide targeted model validation within co-roosting systems of interest. Finally, explicitly tracking the force of infection generated from co-roosting donor species in our models (e.g., bats) to recipient hosts within a foraging environment (e.g., livestock and humans) could directly predict roost prevalence and spillover risks [27,28,68].

The epidemiological models we present here represent general frameworks for generating broad insights into cross-species transmission that can be altered for system-specific analyses. When parameterized for any one system, such models can generate realistic scenarios for how co-roosting species contribute to community-level infection prevalence and spillover risks. In SIRS contexts with low transmission and short infectious and immune periods, we found roost prevalence maximized when species were most phylogenetically related, as supported by many empirical findings [5–10,12]. Further, we also found some contexts where maximum roost prevalence expanded across distantly and unevenly related hosts, supporting increasing spillover risks due to frequent interactions between hosts stemming from extreme spatial overlapping behaviors such as co-roosting and gregariousness [13,22]. These frameworks can function as a template for future modeling of cross-species transmission when species are in close and frequent contact due to a limiting and shared resource. For example, our models can be adapted to better understand host community prevalence in systems like ephemeral bovine virus, where multiple ungulate hosts are infected at ephemeral water sources by arthropod vectors [69], or Hendra virus, where multiple *Pteropus* hosts can infect one another at isolated food sources (e.g., ephemeral *Eucalyptus* blooms, urban fruit and/or nectar sources [70]). Future research modeling cross-species transmission should also expand these frameworks by incorporating other barriers to pathogen transmission, such as social and foraging behaviors, in combination with the evolutionary barriers presented here.

## Data availability

R code is provided in a Github repository on MCS’ Github profile (username: simonimc; https://github.com/simonimc).

## Acknowledgements

We are grateful for the critical feedback on this manuscript provided by Amanda Vicente-Santos, Meggan Craft, Michael Patterson, Trupti Brahmbhatt, Christopher Renner, Meagan Allira, Caroline Cummings, Kristen Dyer, Lauren Lock, Isabelle Sydow, Alicia Roistacher, Taylor Verrett, Desiree Walton, and Chris Wojan.

