## Supporting Information for "A general framework for modeling pathogen transmission in co-roosting host communities"

**Running Title:** Modeling infection in co-roosting hosts

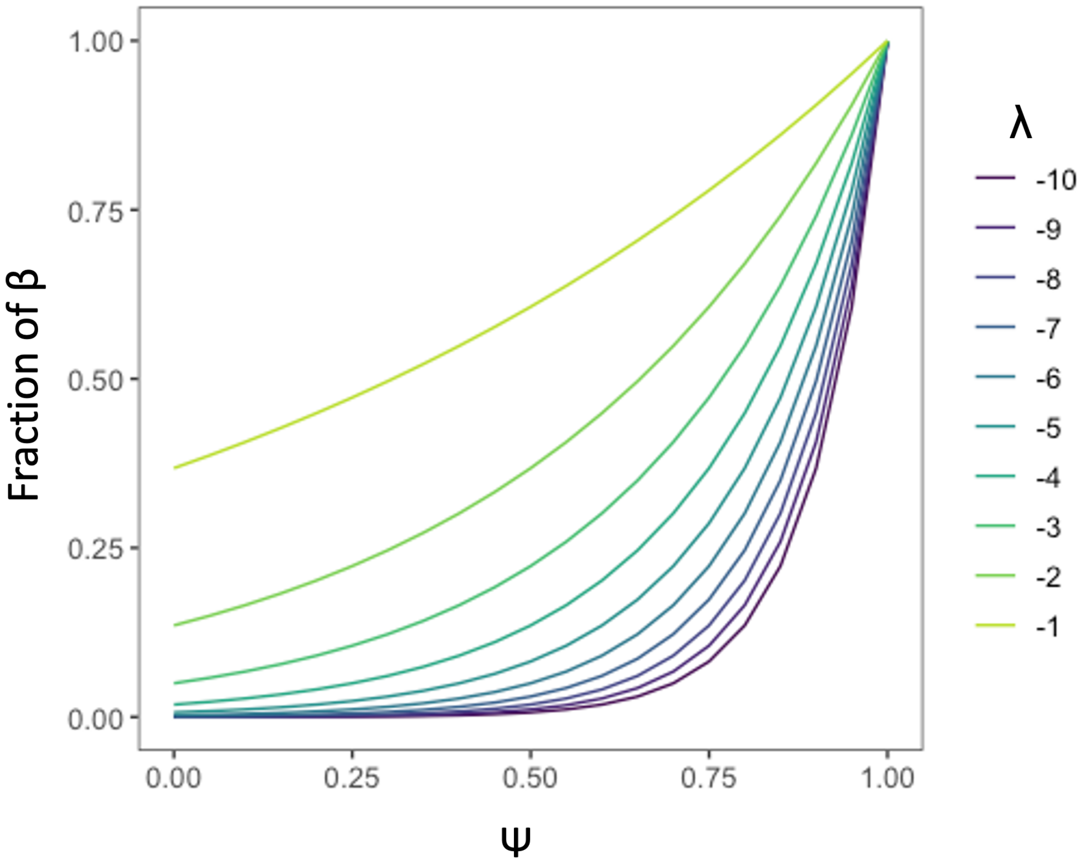


**Figure S1.** Examples of functional relationships between host phylogenetic similarity and pathogen transmission between host species. The fraction of intraspecific transmission (β) is determined by the function $e^{(1-\Psi)\lambda}$, where Ψ is the correlation coefficient of phylogenetic relatedness between species and λ is the shape of the curve (λ = - 6 for all models in this manuscript). Colors represent different λ values.

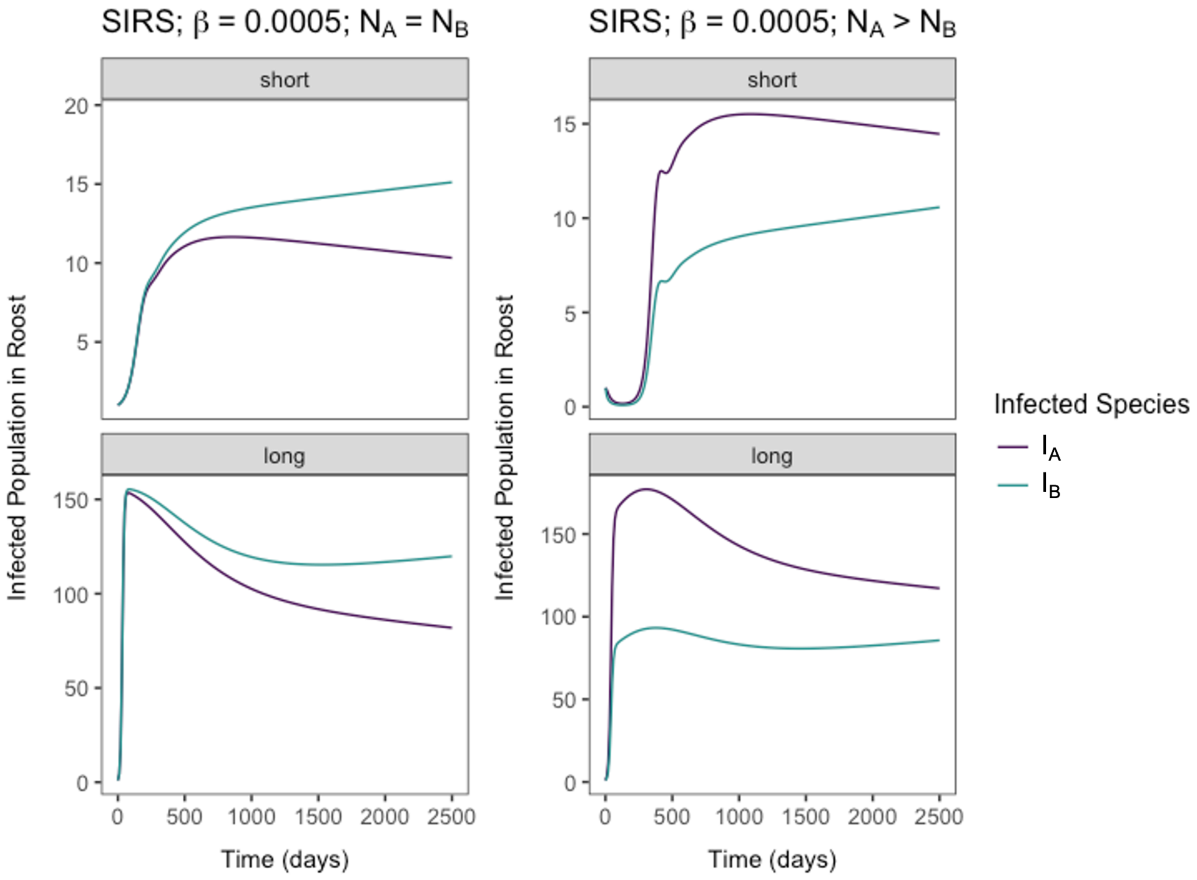


**Figure S2.** Time series examples of two-species SIRS models under the lowest interspecific transmission (β) and varying starting population sizes. For short parameter spaces, pathogen clearance rates (γ) are 1/7, and waning immunity rates (ε) are 1/30. For long parameter spaces, pathogen clearance rates (γ) are 1/730, and waning immunity rates (ε) are 1/1095. Species in each example are 99.99% related to one another (Ψ). Note that y-axes are different across plots.

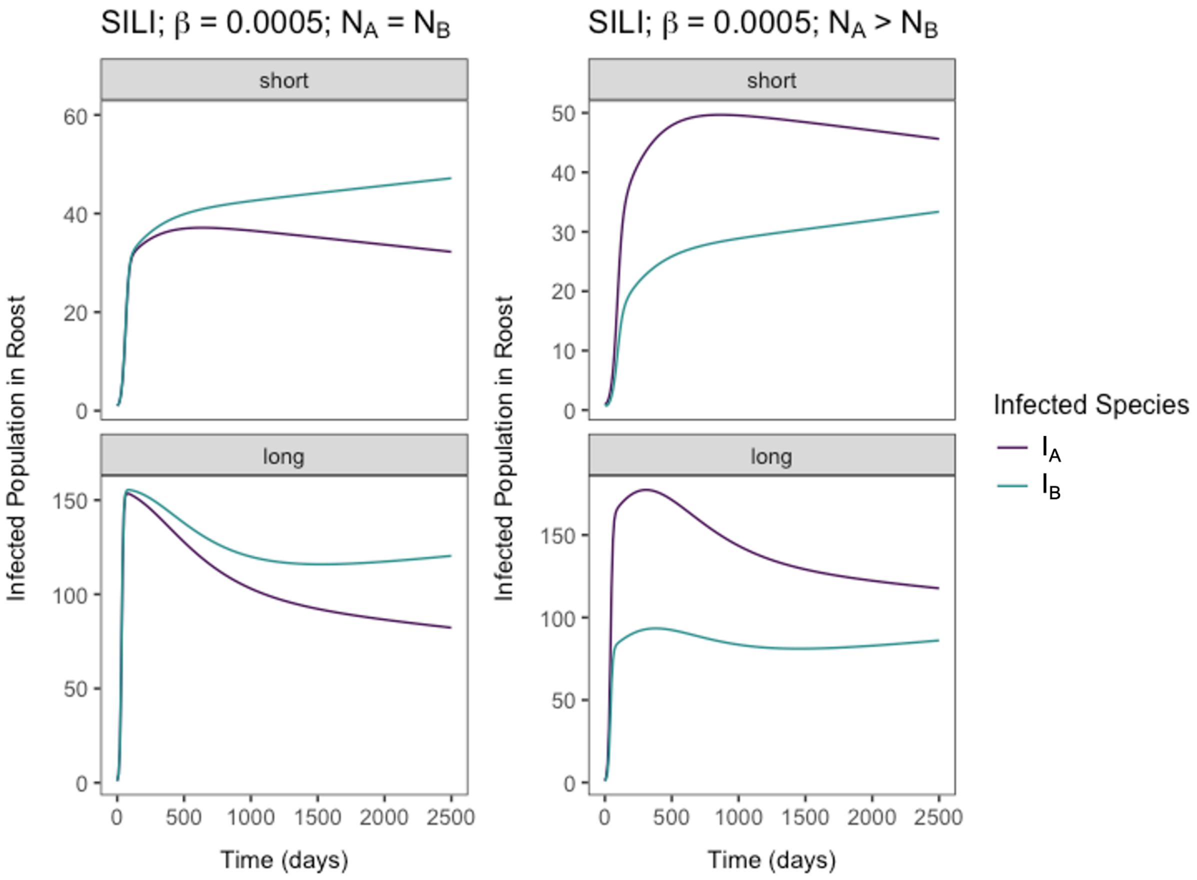


**Figure S3.** Time series examples of two-species SILI models under the lowest interspecific transmission (β) and varying starting population sizes. For short parameter spaces, pathogen clearance rates (γ) are 1/7, and waning immunity rates (ε) are 1/30. For long parameter spaces, pathogen clearance rates (γ) are 1/730, and waning immunity rates (ε) are 1/1095. Species in each example are 99.99% related to one another (Ψ). Note that y-axes are different across plots.

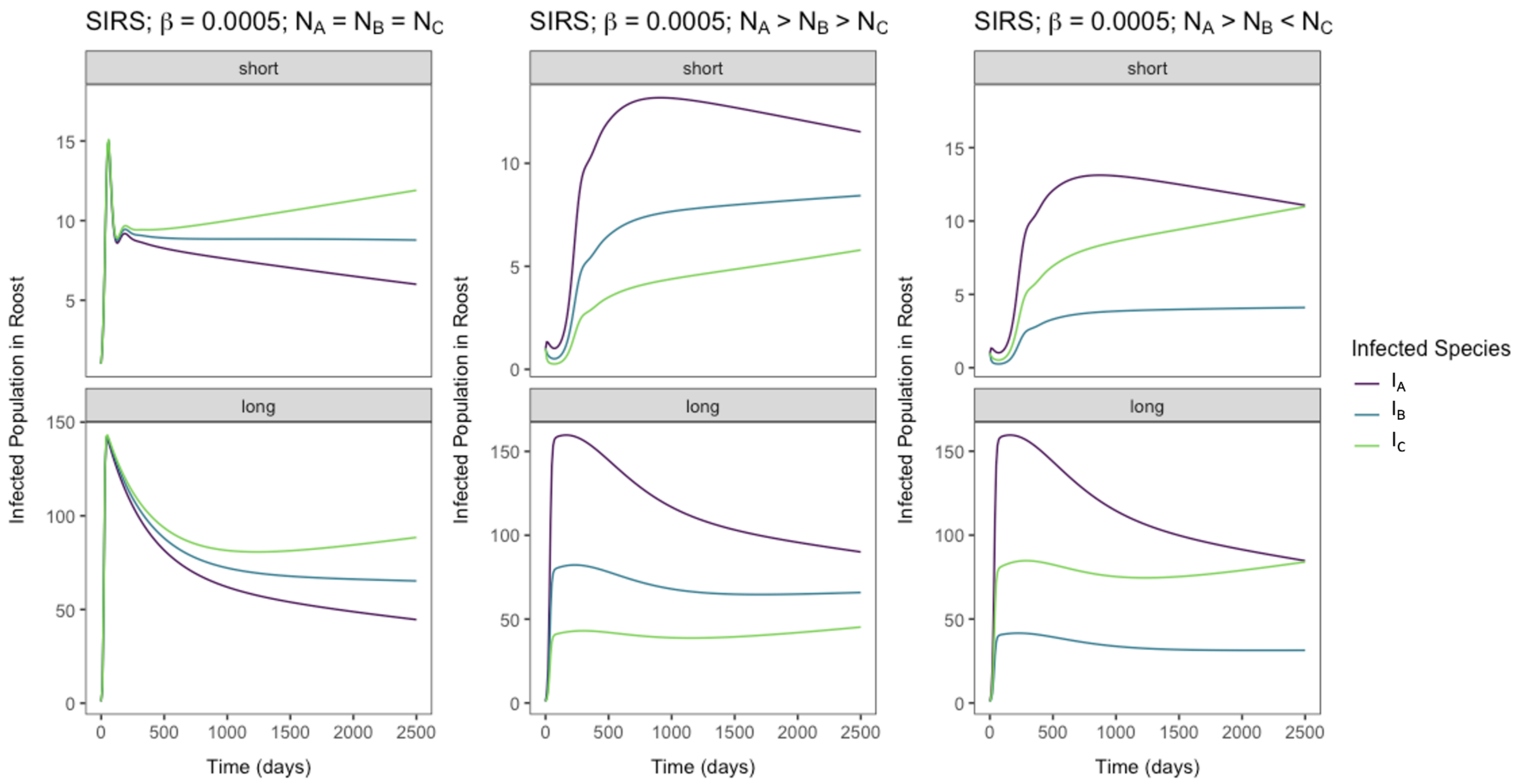


**Figure S4.** Time series examples of three-species SIRS models under the lowest interspecific transmission (β) and varying starting population sizes. For short parameter spaces, pathogen clearance rates (γ) are 1/7, and waning immunity rates (ε) are 1/30. For long parameter spaces, pathogen clearance rates (γ) are 1/730, and waning immunity rates (ε) are 1/1095. All three species in each example are 99.99% related to one another ($\Psi_{ij}$). Note that y-axes are different across plots.

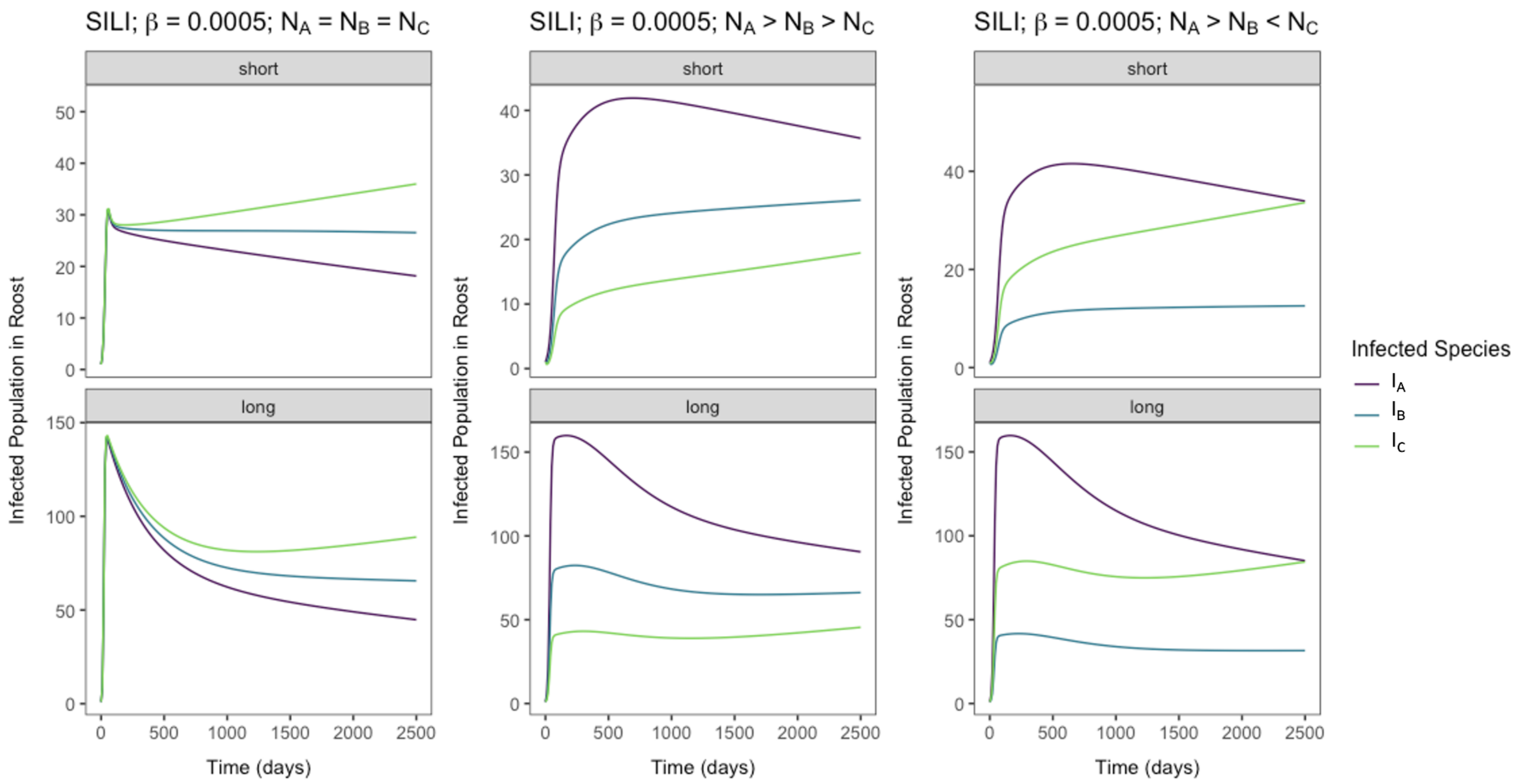


**Figure S5.** Time series examples of three-species SILI models under the lowest interspecific transmission (β) and varying starting population sizes. For short parameter spaces, pathogen clearance rates (γ) are 1/7, and infection reactivation rates (ε) are 1/30. For long parameter spaces, pathogen clearance rates (γ) are 1/730, and infection reactivation rates (ε) are 1/1095. All three species in each example are 99.99% related to one another ($\Psi_{ij}$). Note that y-axes are different across plots.

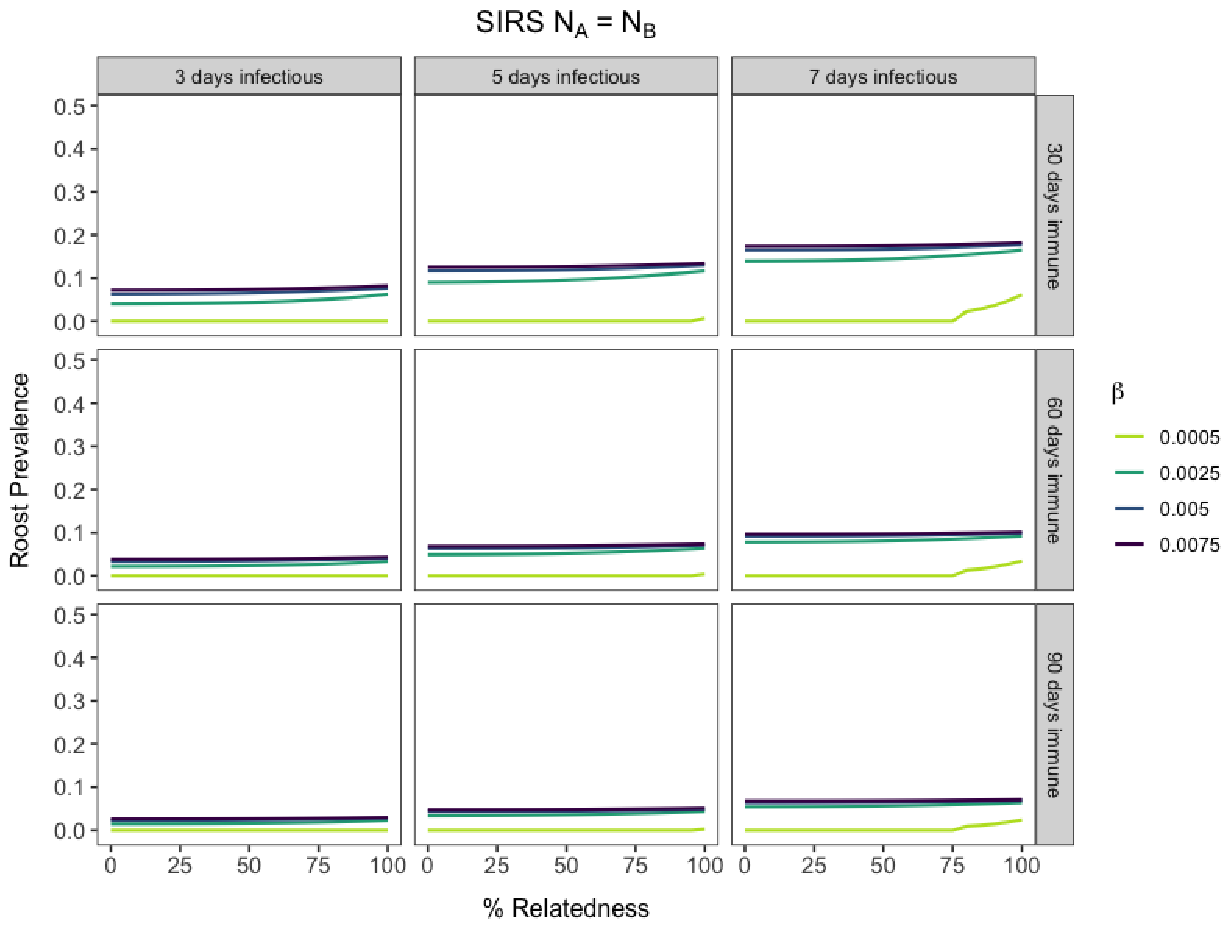


**Figure S6.** Community roost prevalence dynamics vary with phylogenetic relatedness between two host species at equilibrium. Results from two-species SIRS models across parameter space for varying short infectious and waning immunity periods and equal starting populations for both host species. Colors indicate varying intraspecific transmission values.

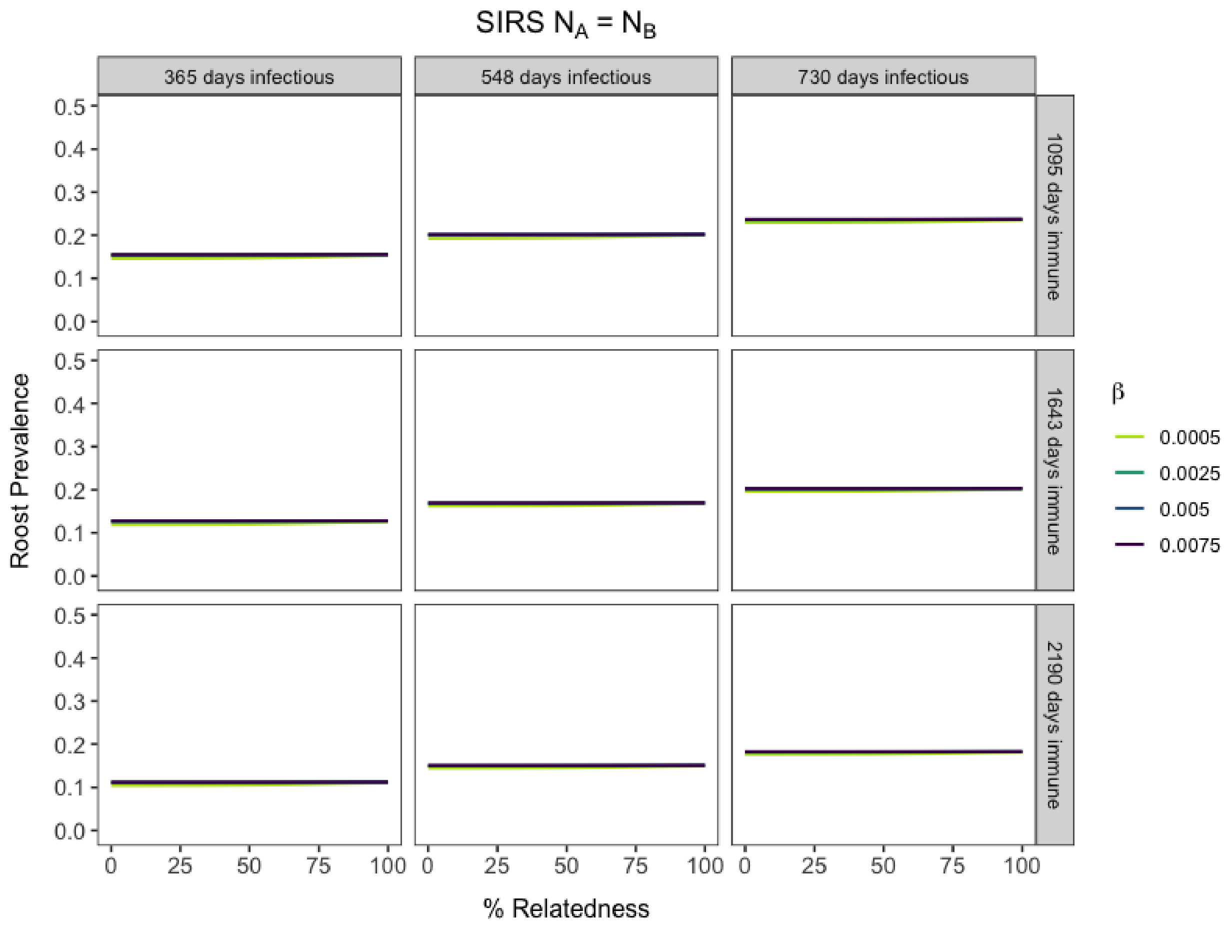


**Figure S7.** Community roost prevalence dynamics vary with phylogenetic relatedness between two host species at equilibrium. Results from two-species SIRS models across parameter space for varying long infectious and waning immunity periods and equal starting populations for both host species. Colors indicate varying intraspecific transmission values.

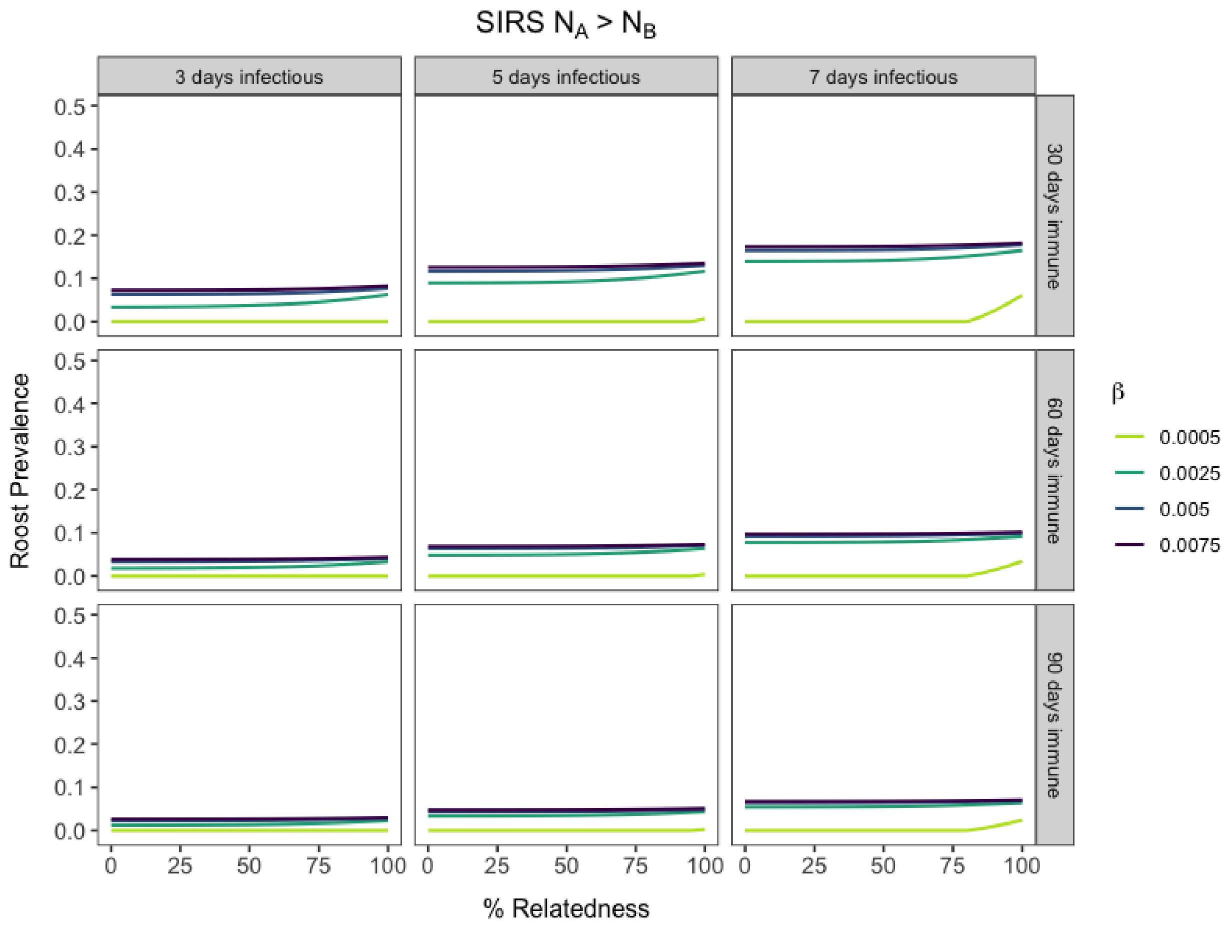


**Figure S8.** Community roost prevalence dynamics vary with phylogenetic relatedness between two host species at equilibrium. Results from two-species SIRS models across parameter space for varying short infectious and waning immunity periods and when species A's starting populations are dominant over species B. Colors indicate varying intraspecific transmission values.

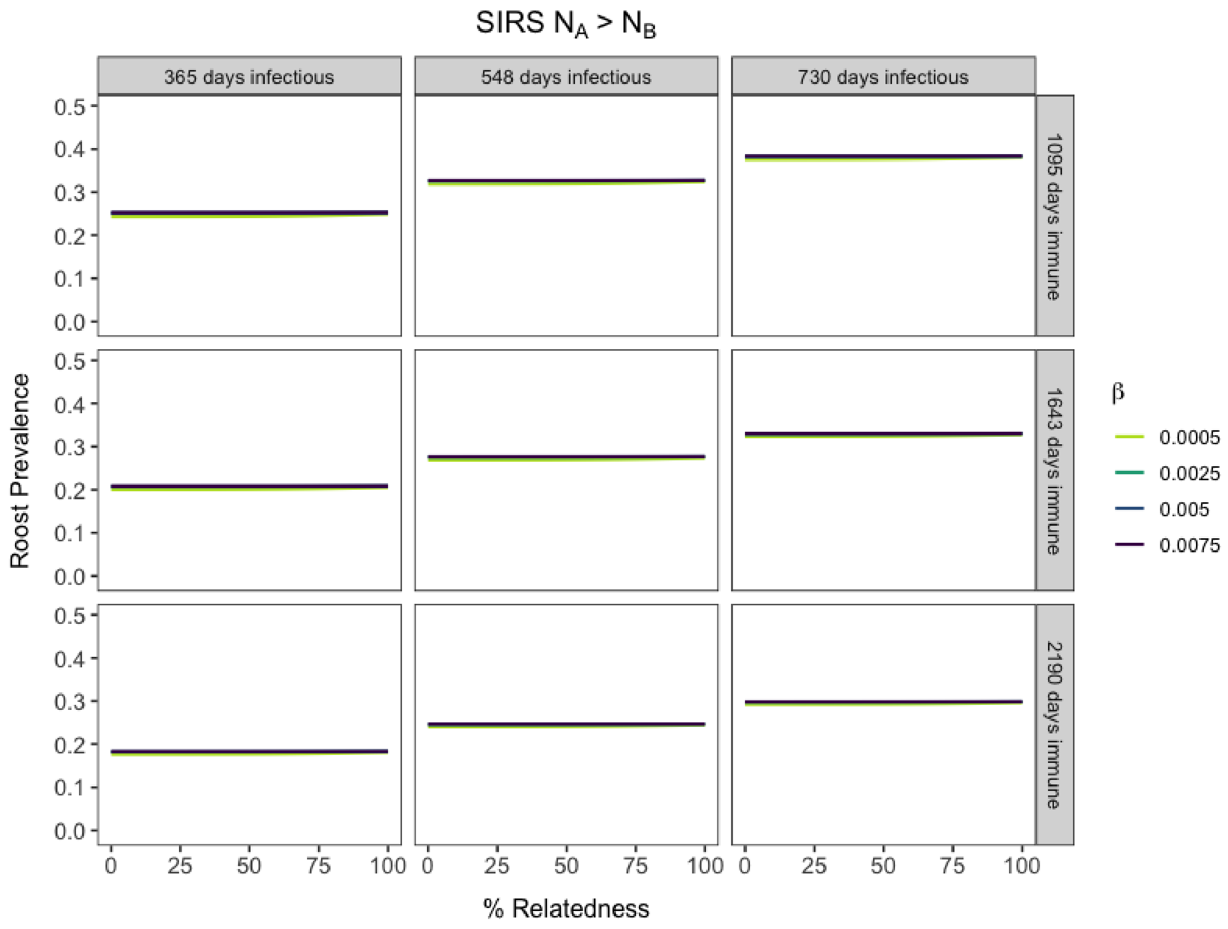


**Figure S9.** Community roost prevalence dynamics vary with phylogenetic relatedness between two host species at equilibrium. Results from two-species SIRS models across parameter space for varying long infectious and waning immunity periods and when species A's starting populations are dominant over species B. Colors indicate varying intraspecific transmission values.

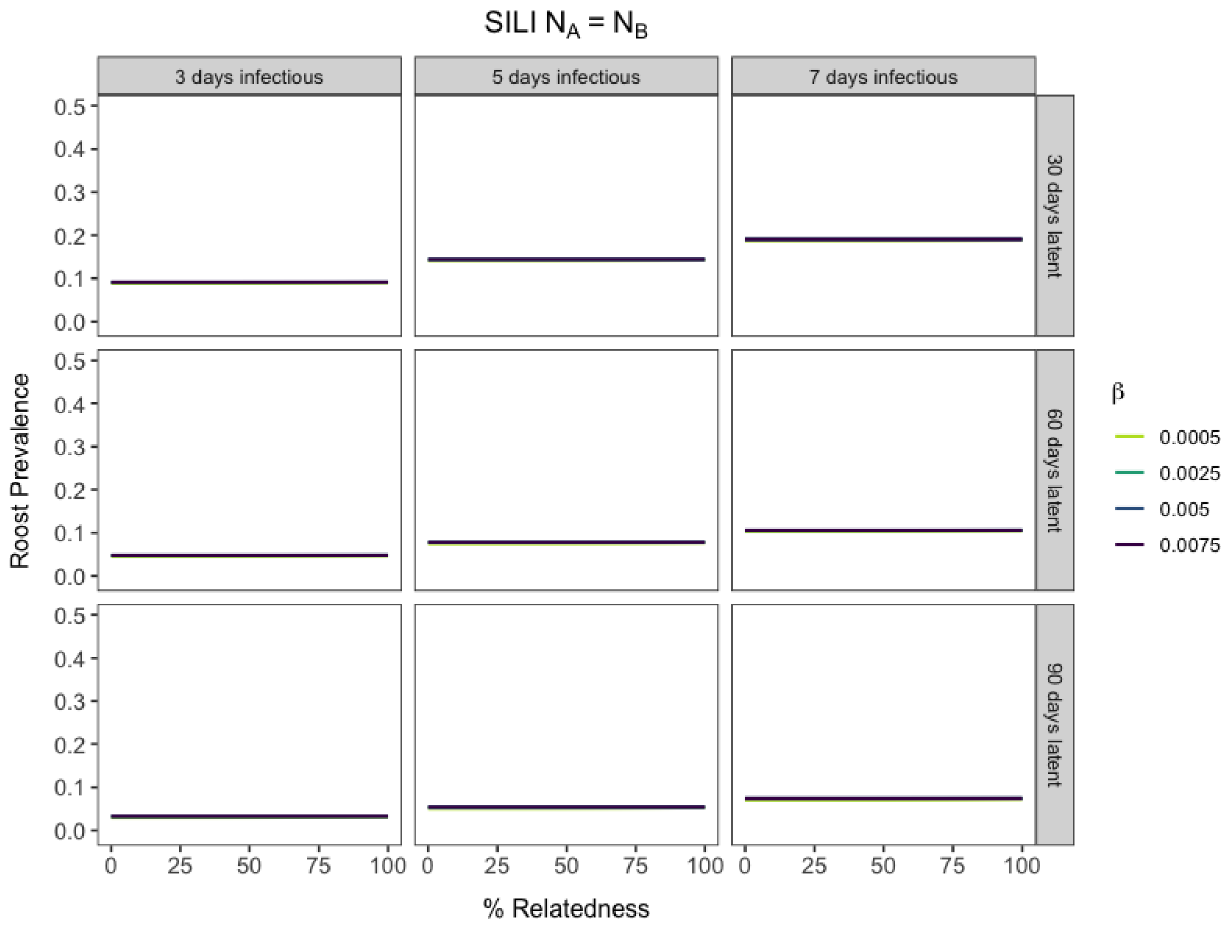


**Figure S10.** Community roost prevalence dynamics vary with phylogenetic relatedness between two host species at equilibrium. Results from two-species SILI models across parameter space for varying short infectious and waning immunity periods and equal starting populations for both host species. Colors indicate varying intraspecific transmission values.

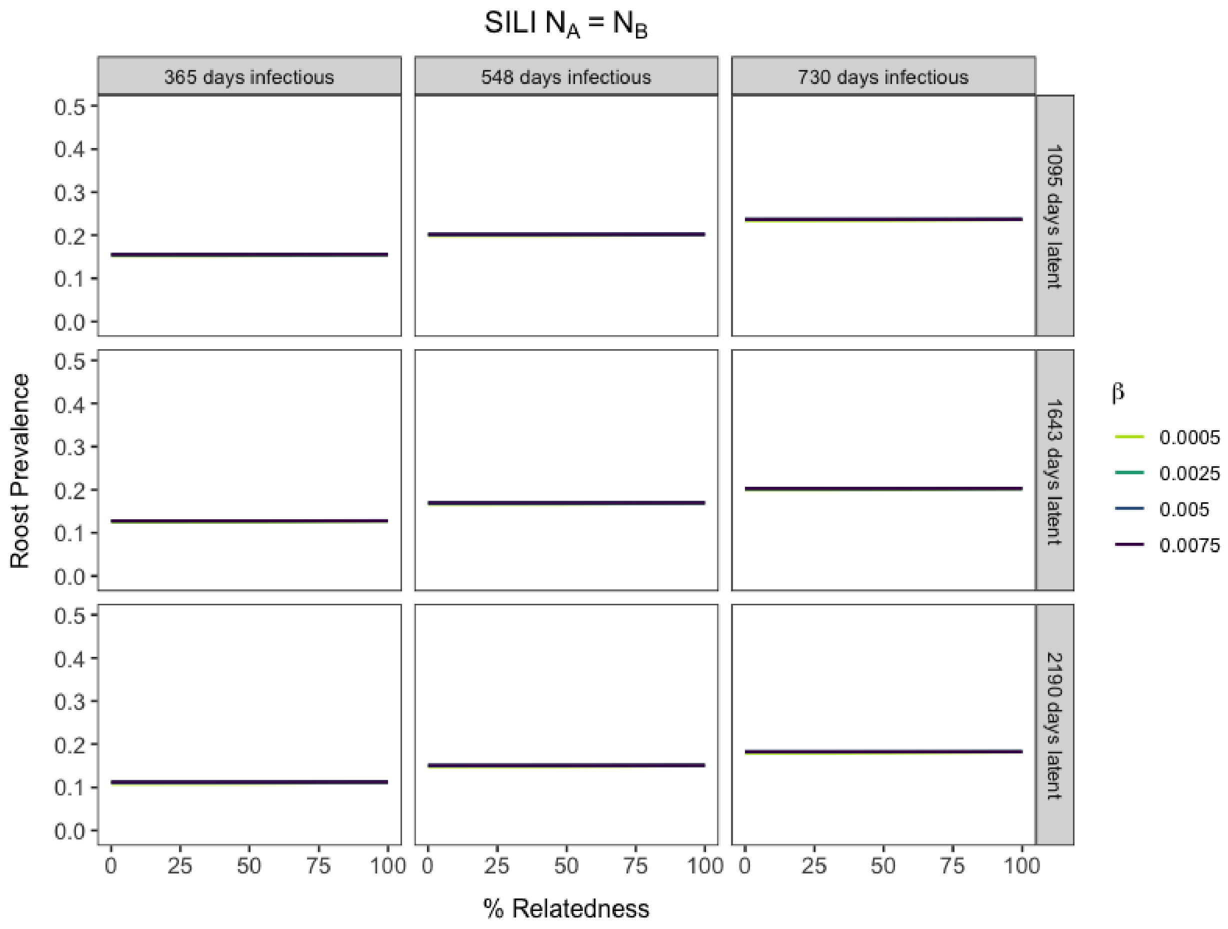


**Figure S11.** Community roost prevalence dynamics vary with phylogenetic relatedness between two host species at equilibrium. Results from two-species SILI models across parameter space for varying long infectious and waning immunity periods and equal starting populations for both host species. Colors indicate varying intraspecific transmission values.

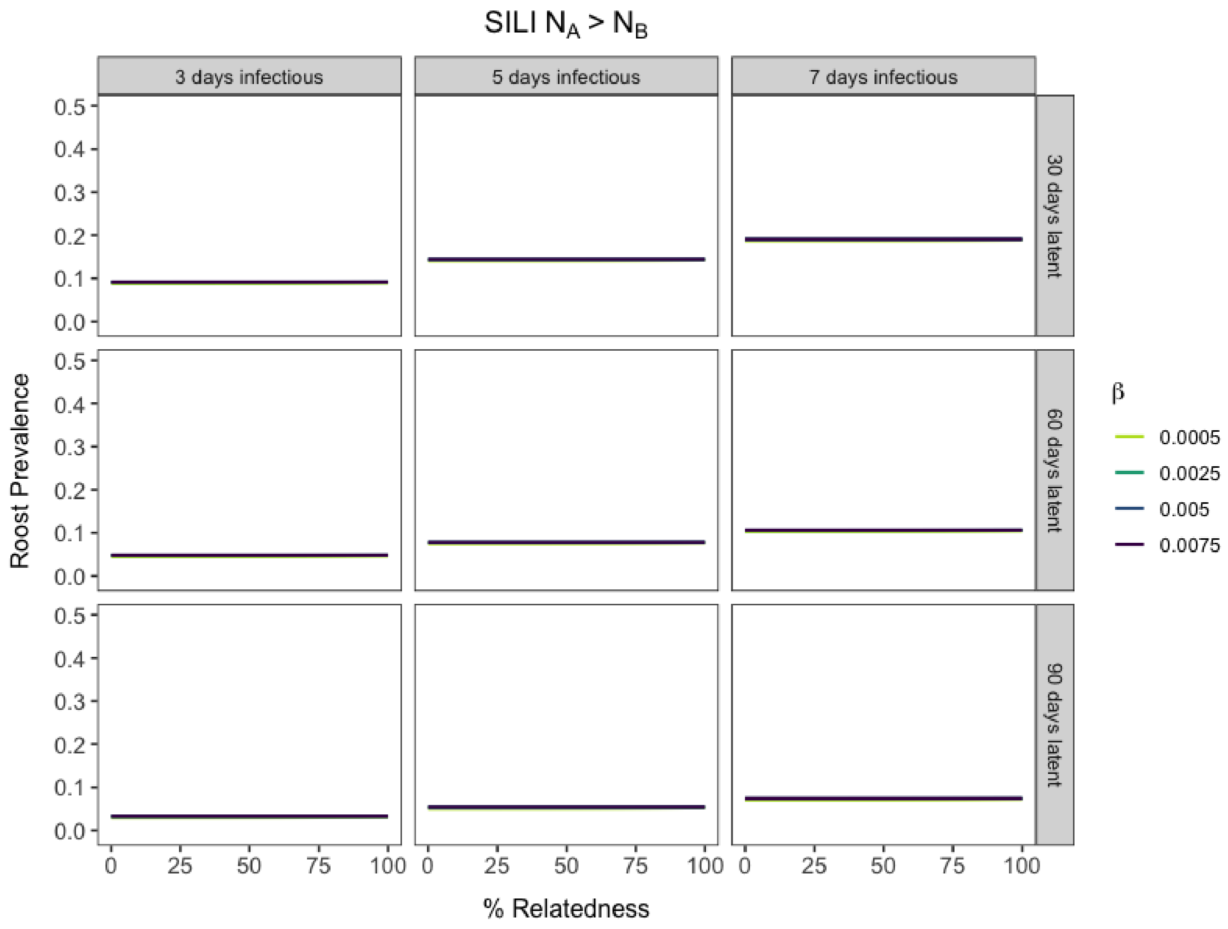


**Figure S12.** Community roost prevalence dynamics vary with phylogenetic relatedness between two host species at equilibrium. Results from two-species SILI models across parameter space for varying short infectious and waning immunity periods and when species A's starting populations are dominant over species B. Colors indicate varying intraspecific transmission values.

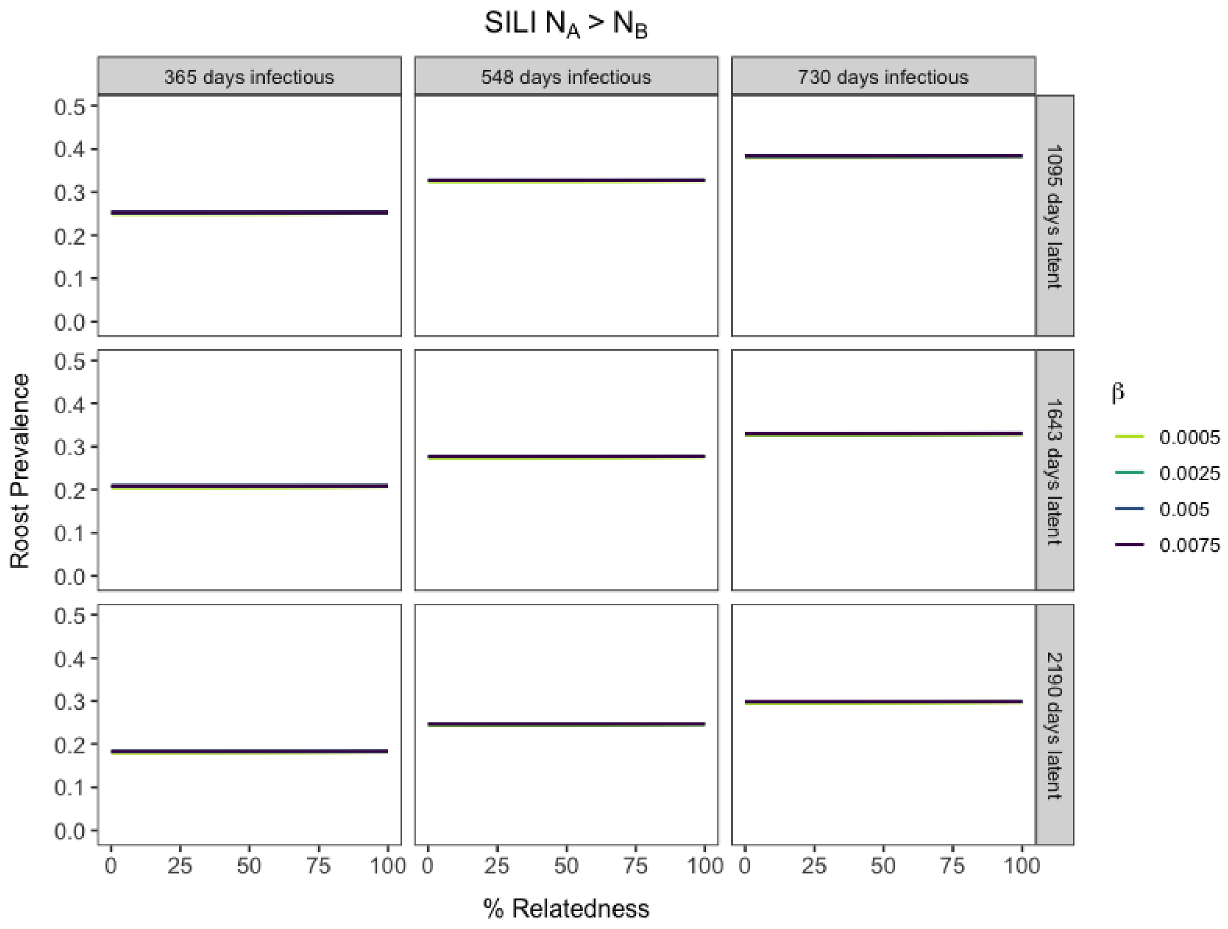


**Figure S13.** Community roost prevalence dynamics vary with phylogenetic relatedness between two host species at equilibrium. Results from two-species SILI models across parameter space for varying long infectious and waning immunity periods and when species A's starting populations are dominant over species B. Colors indicate varying intraspecific transmission values.

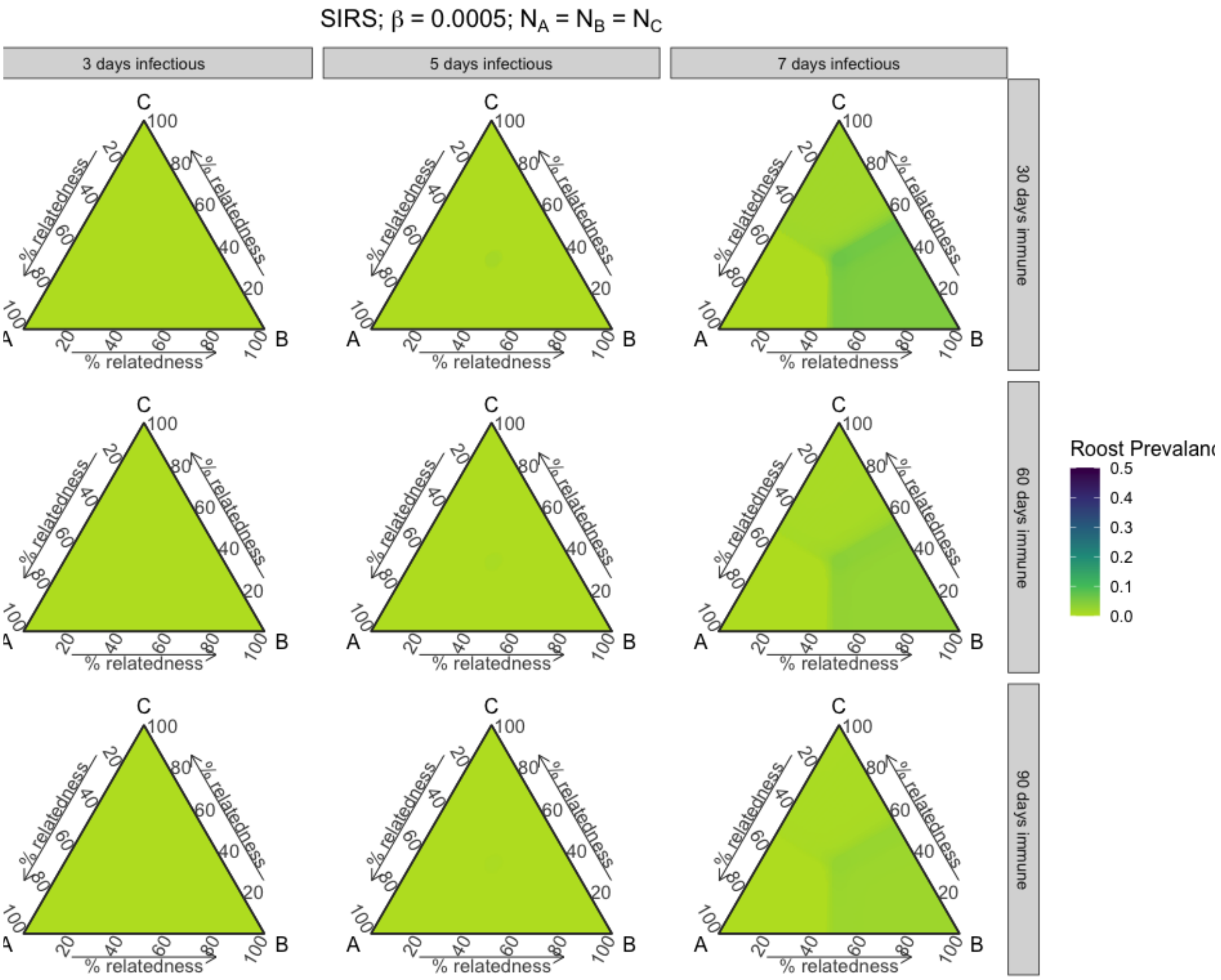


**Figure S14.** Roost prevalence dynamics vary with phylogenetic relatedness between three host species at equilibrium. Results from three-species SIRS models across parameter space for varying short infectious and waning immunity periods, when species starting populations are equal, and when intraspecific transmission (β) is 0.0005.

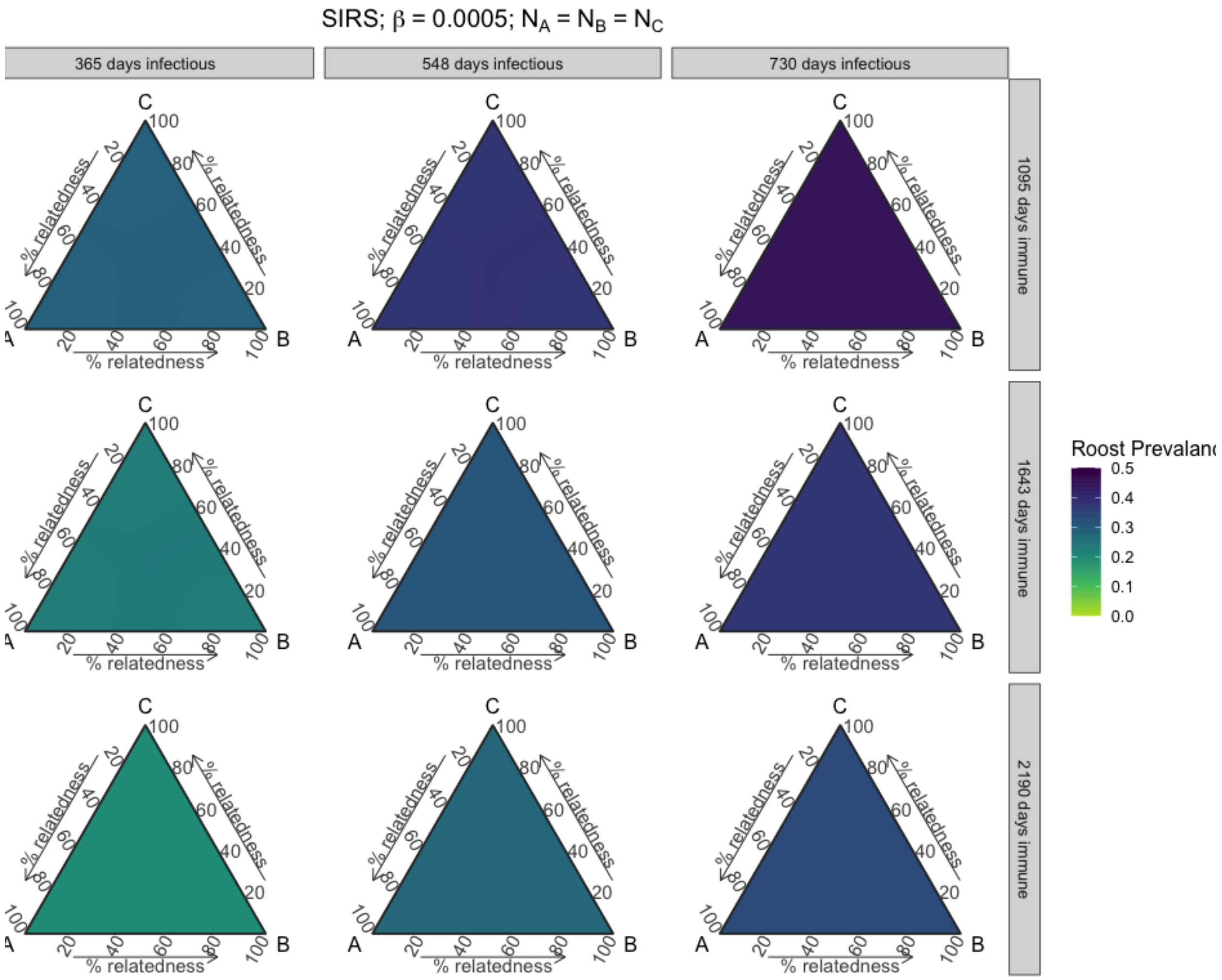


**Figure S15.** Roost prevalence dynamics vary with phylogenetic relatedness between three host species at equilibrium. Results from three-species SIRS models across parameter space for varying long infectious and waning immunity periods, when species starting populations are equal, and when intraspecific transmission (β) is 0.0005.

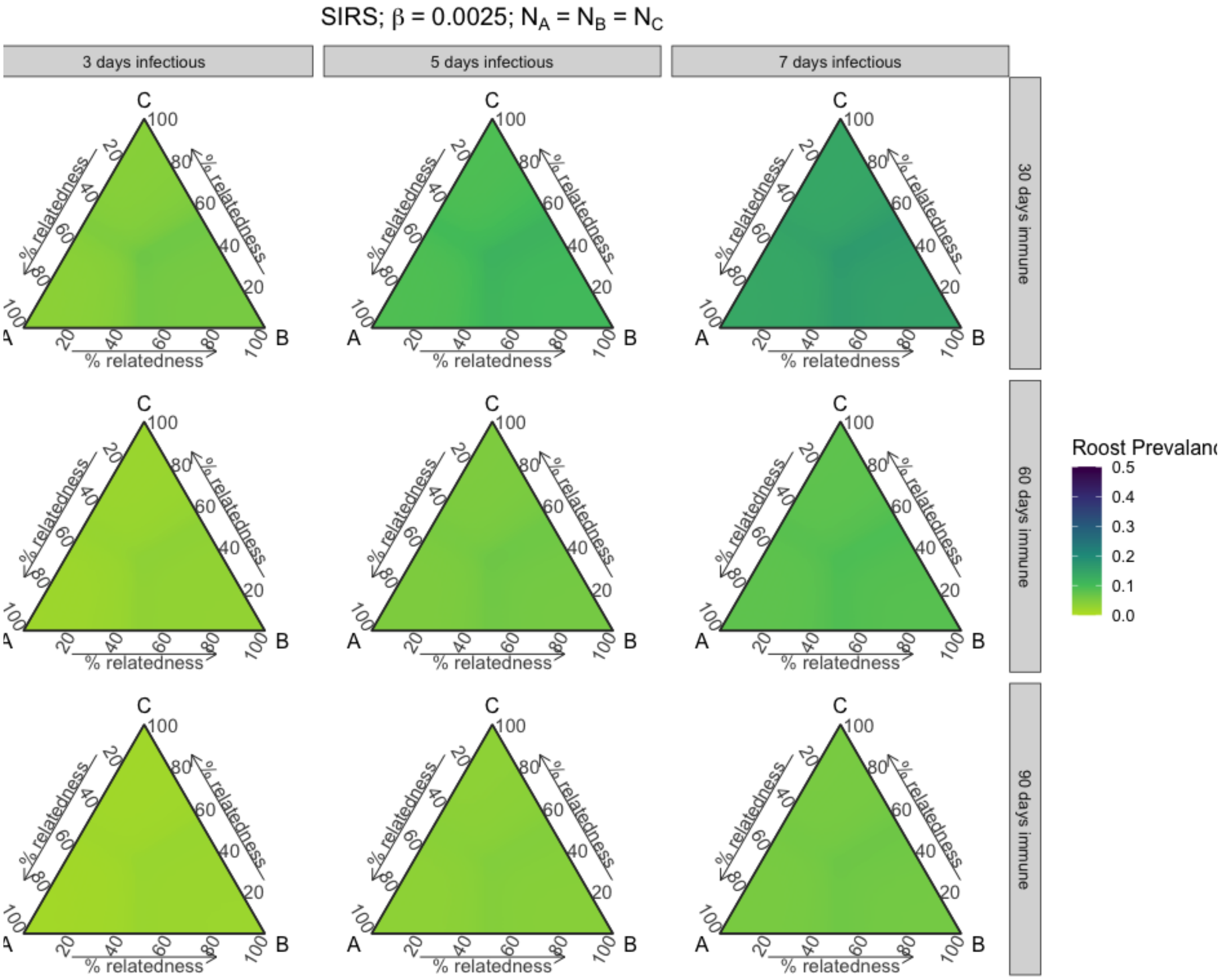


**Figure S16.** Roost prevalence dynamics vary with phylogenetic relatedness between three host species at equilibrium. Results from three-species SIRS models across parameter space for varying short infectious and waning immunity periods, when species starting populations are equal, and when intraspecific transmission (β) is 0.0025.

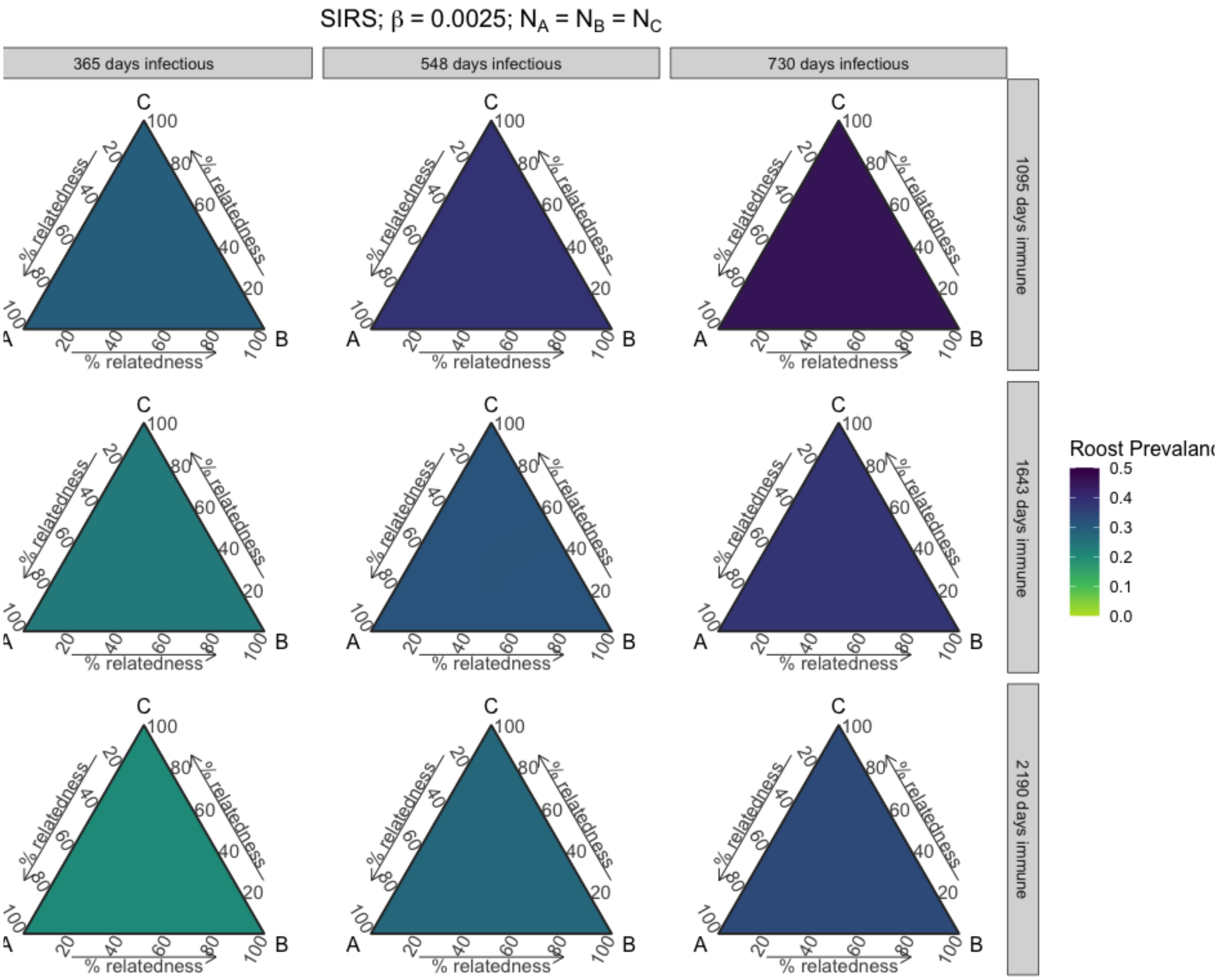


**Figure S17.** Roost prevalence dynamics vary with phylogenetic relatedness between three host species at equilibrium. Results from three-species SIRS models across parameter space for varying long infectious and waning immunity periods, when species starting populations are equal, and when intraspecific transmission (β) is 0.0025.

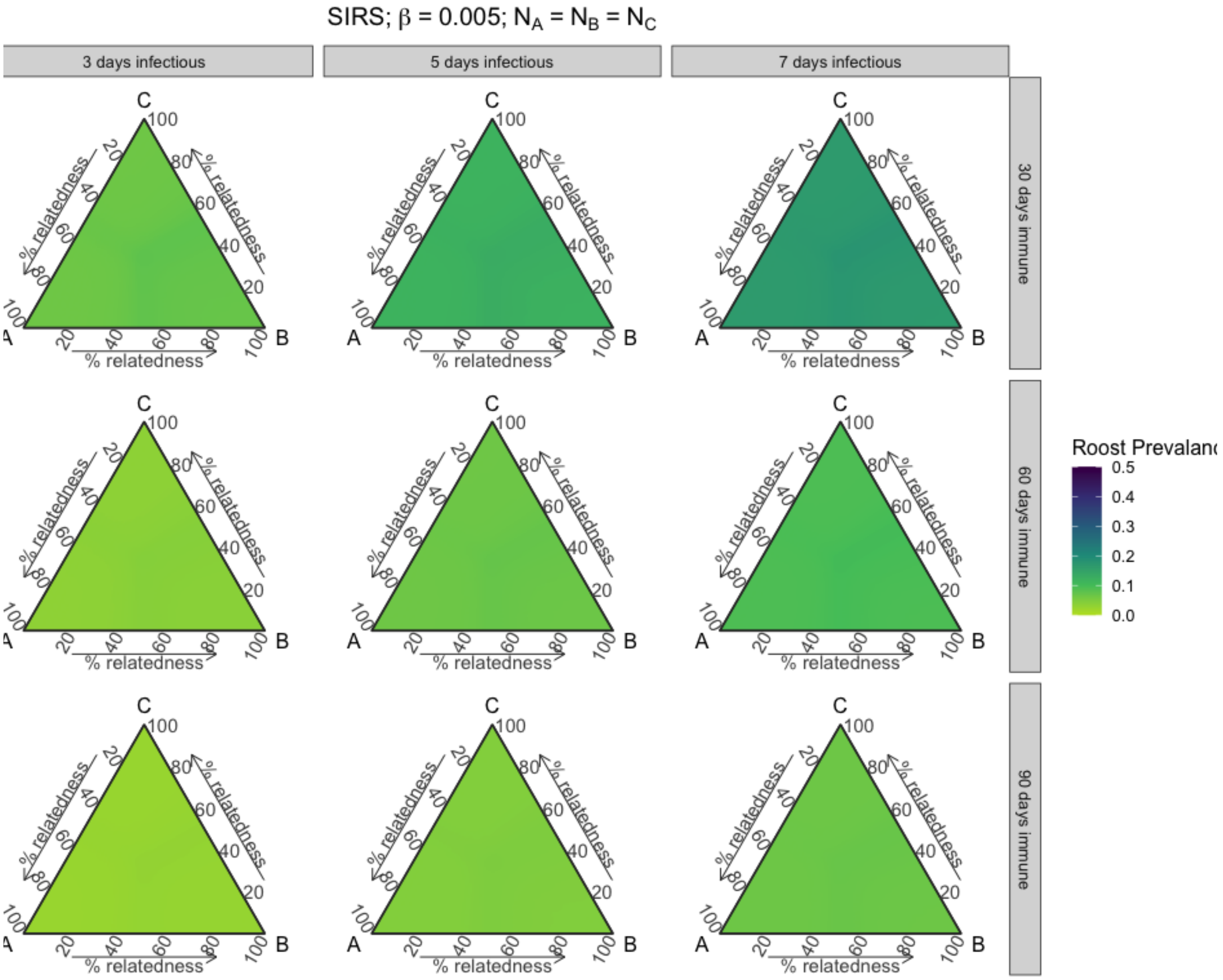


**Figure S18.** Roost prevalence dynamics vary with phylogenetic relatedness between three host species at equilibrium. Results from three-species SIRS models across parameter space for varying short infectious and waning immunity periods, when species starting populations are equal, and when intraspecific transmission (β) is 0.005.

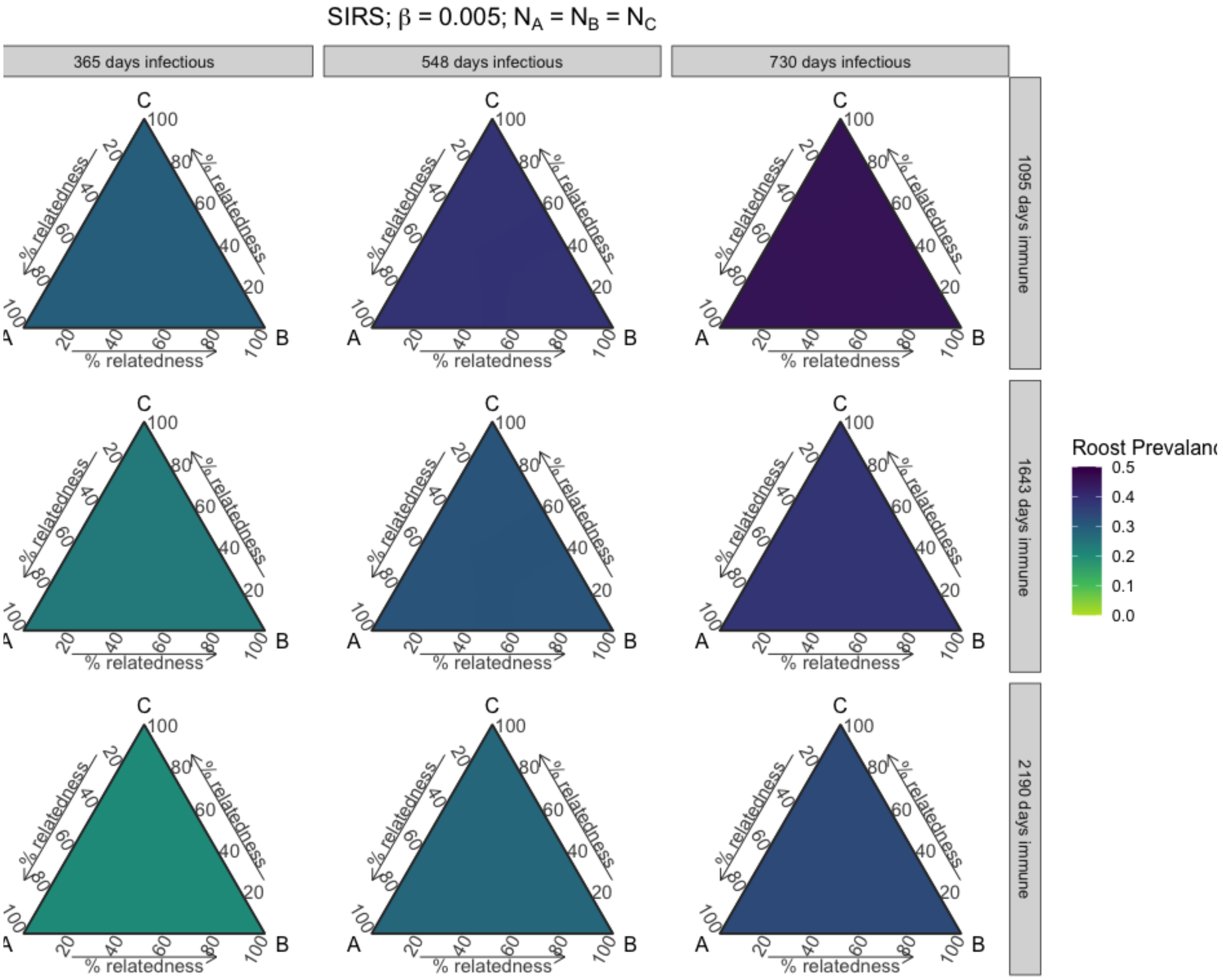


**Figure S19.** Roost prevalence dynamics vary with phylogenetic relatedness between three host species at equilibrium. Results from three-species SIRS models across parameter space for varying long infectious and waning immunity periods, when species starting populations are equal, and when intraspecific transmission (β) is 0.005.

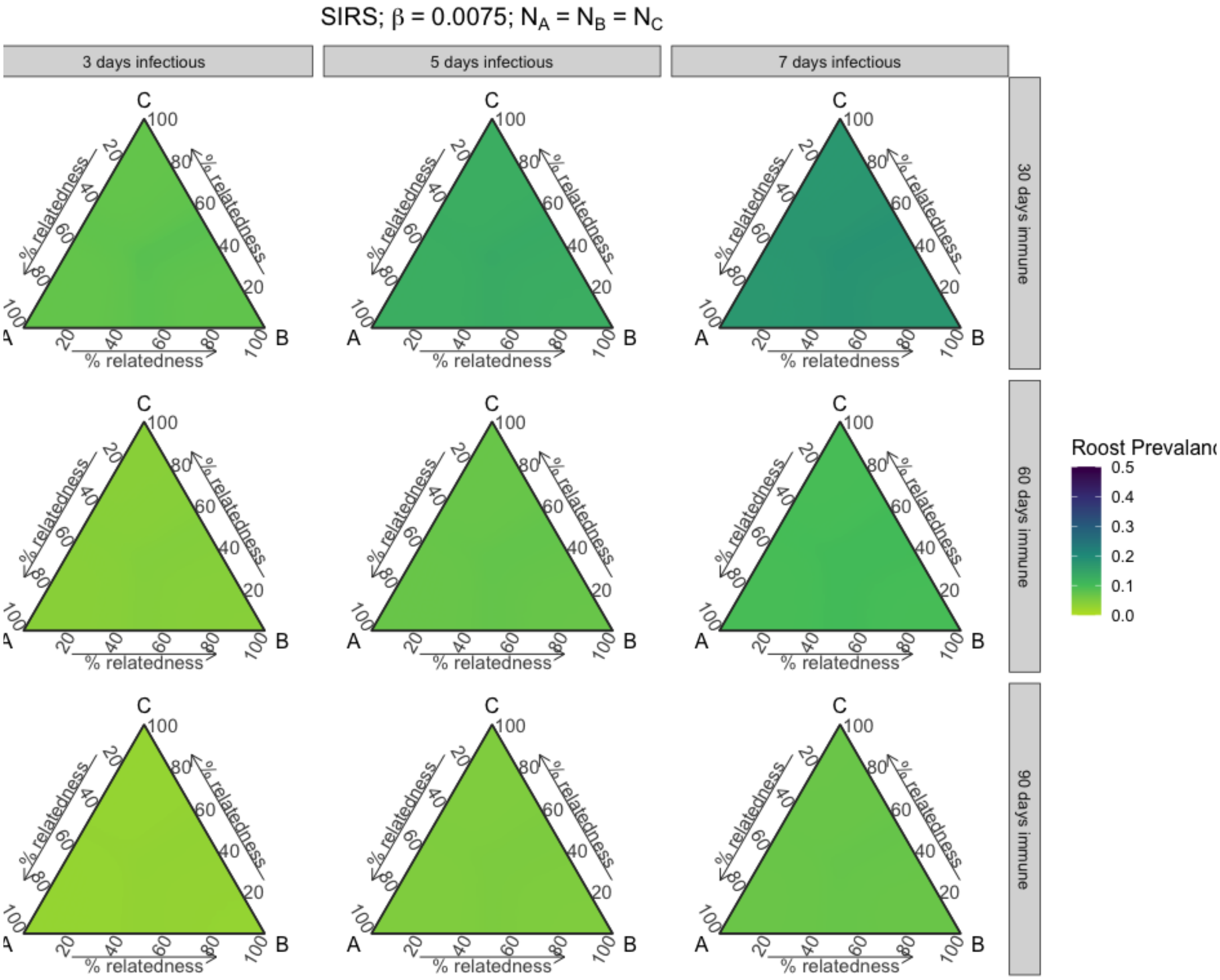


**Figure S20.** Roost prevalence dynamics vary with phylogenetic relatedness between three host species at equilibrium. Results from three-species SIRS models across parameter space for varying short infectious and waning immunity periods, when species starting populations are equal, and when intraspecific transmission (β) is 0.0075.

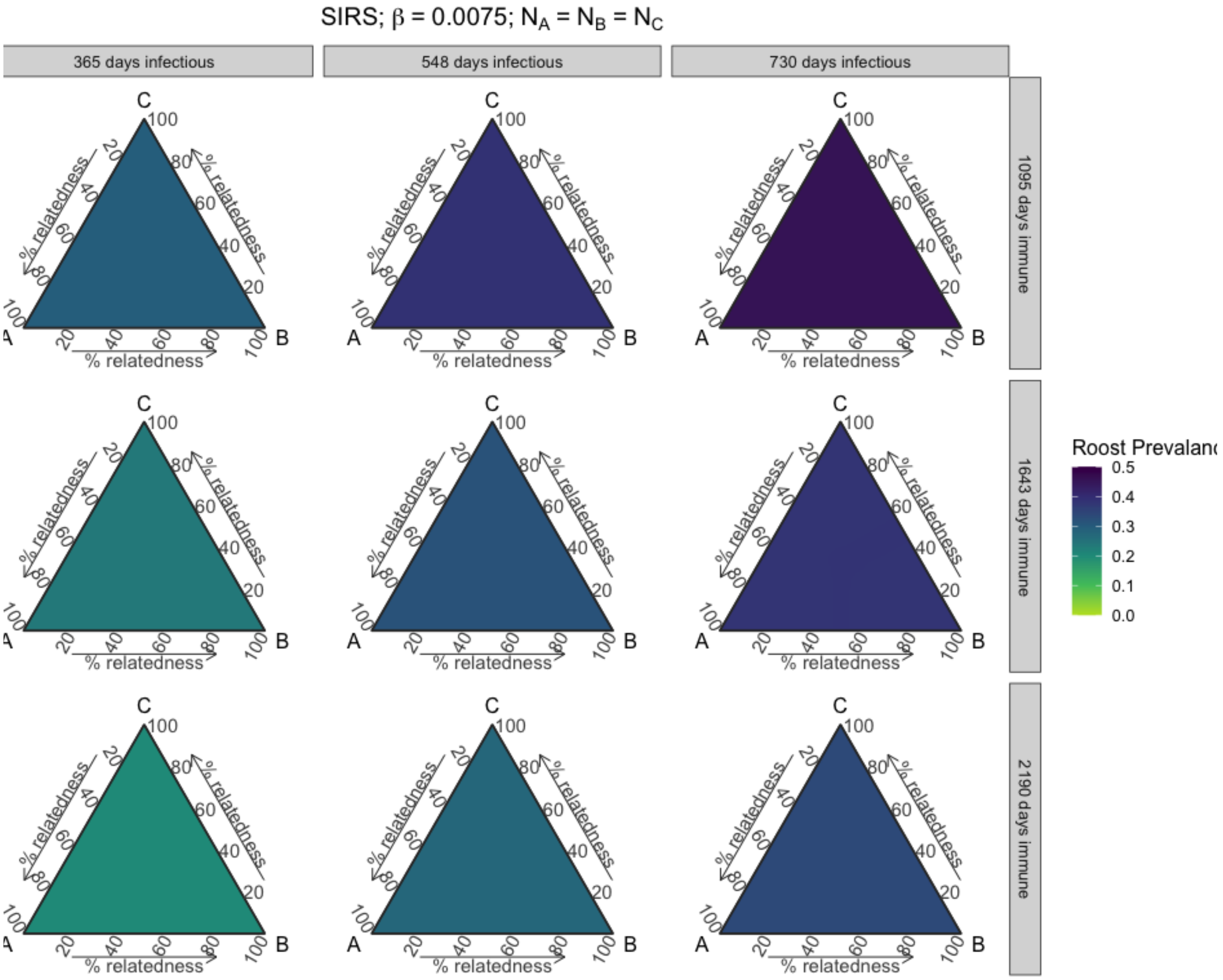


**Figure S21.** Roost prevalence dynamics vary with phylogenetic relatedness between three host species at equilibrium. Results from three-species SIRS models across parameter space for varying long infectious and waning immunity periods, when species starting populations are equal, and when intraspecific transmission (β) is 0.0075.

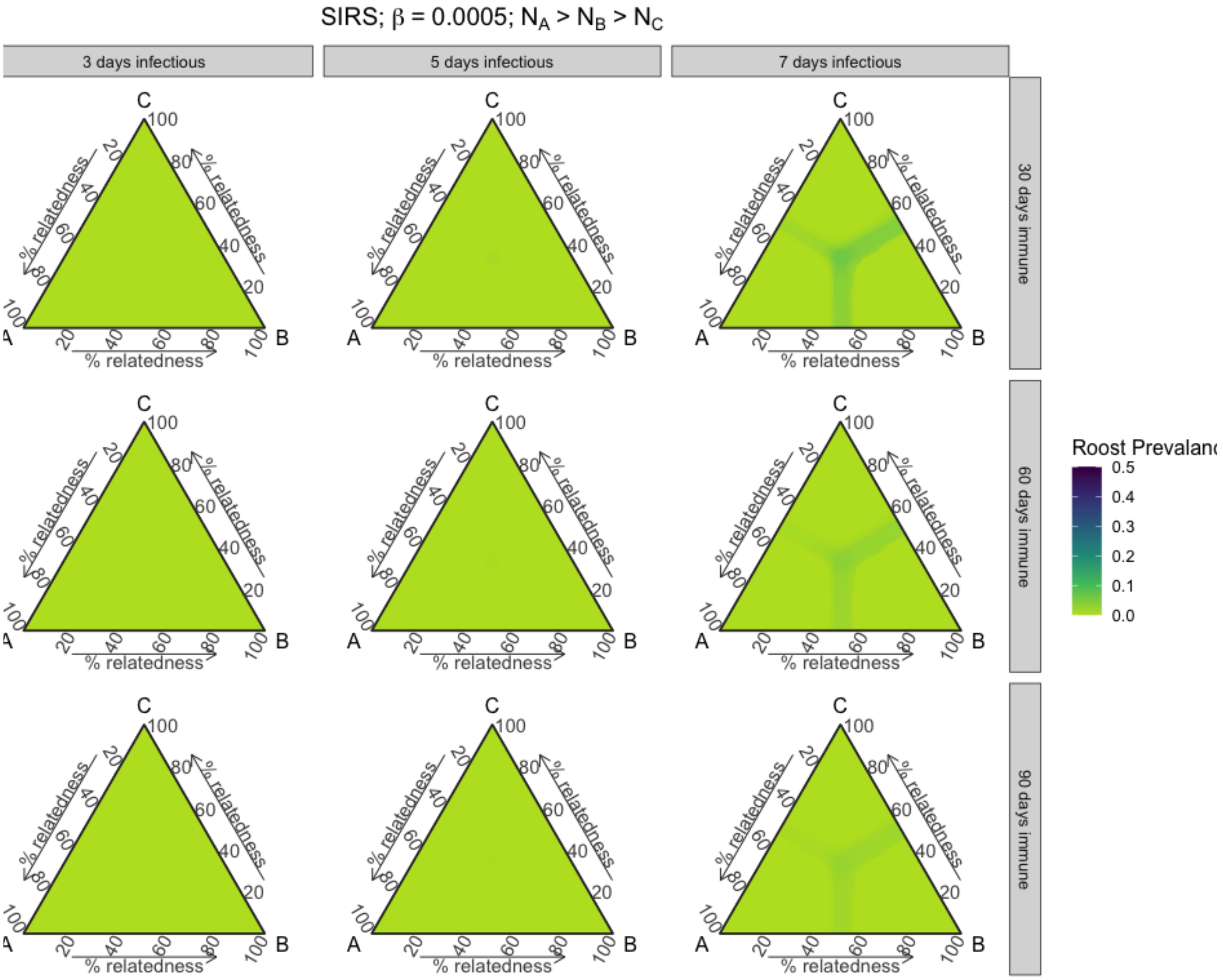


**Figure S22.** Roost prevalence dynamics vary with phylogenetic relatedness between three host species at equilibrium. Results from three-species SIRS models across parameter space for varying short infectious and waning immunity periods, when species starting populations sequentially decrease, and when intraspecific transmission (β) is 0.0005.

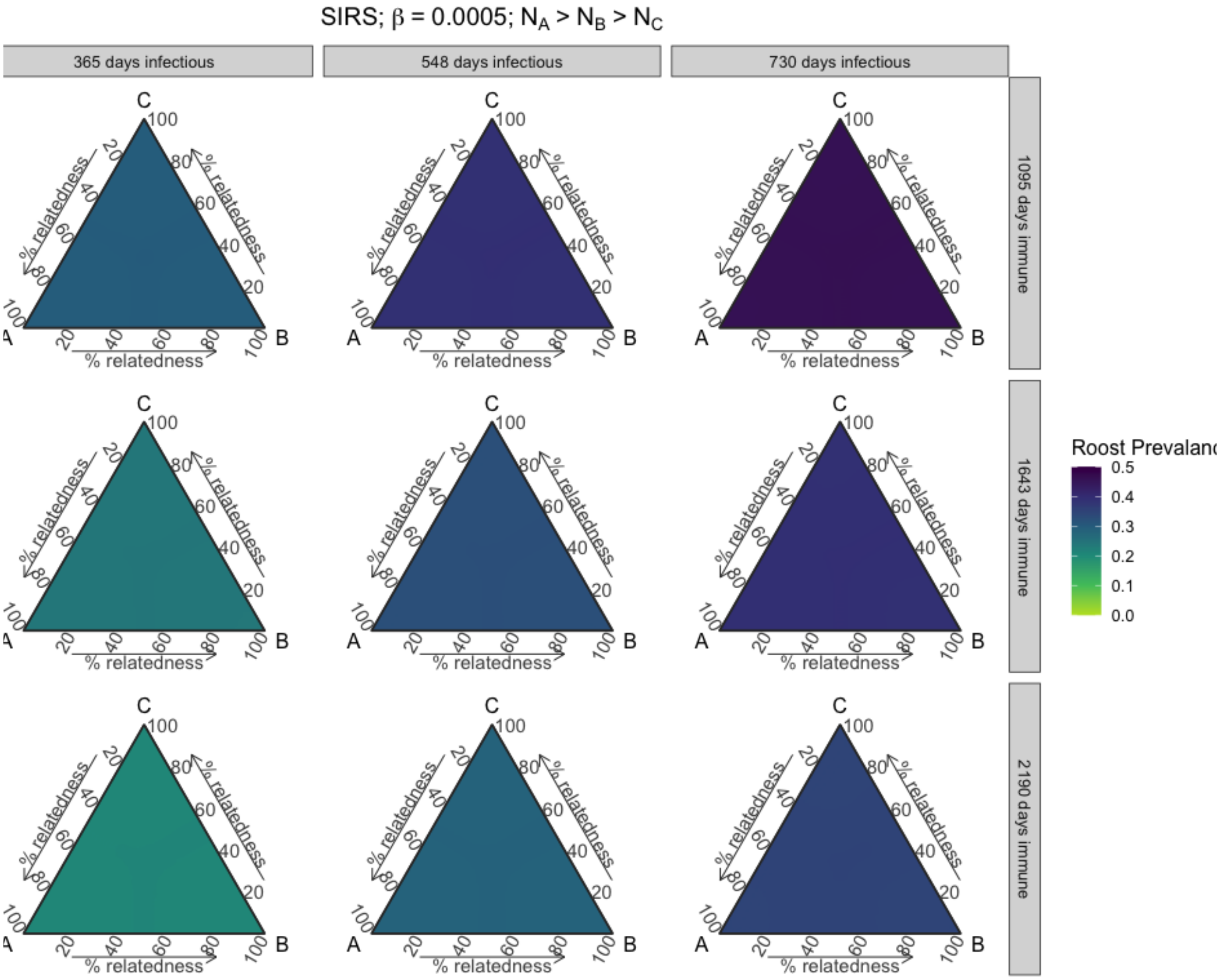


**Figure S23.** Roost prevalence dynamics vary with phylogenetic relatedness between three host species at equilibrium. Results from three-species SIRS models across parameter space for varying long infectious and waning immunity periods, when species starting populations sequentially decrease, and when intraspecific transmission (β) is 0.0005.

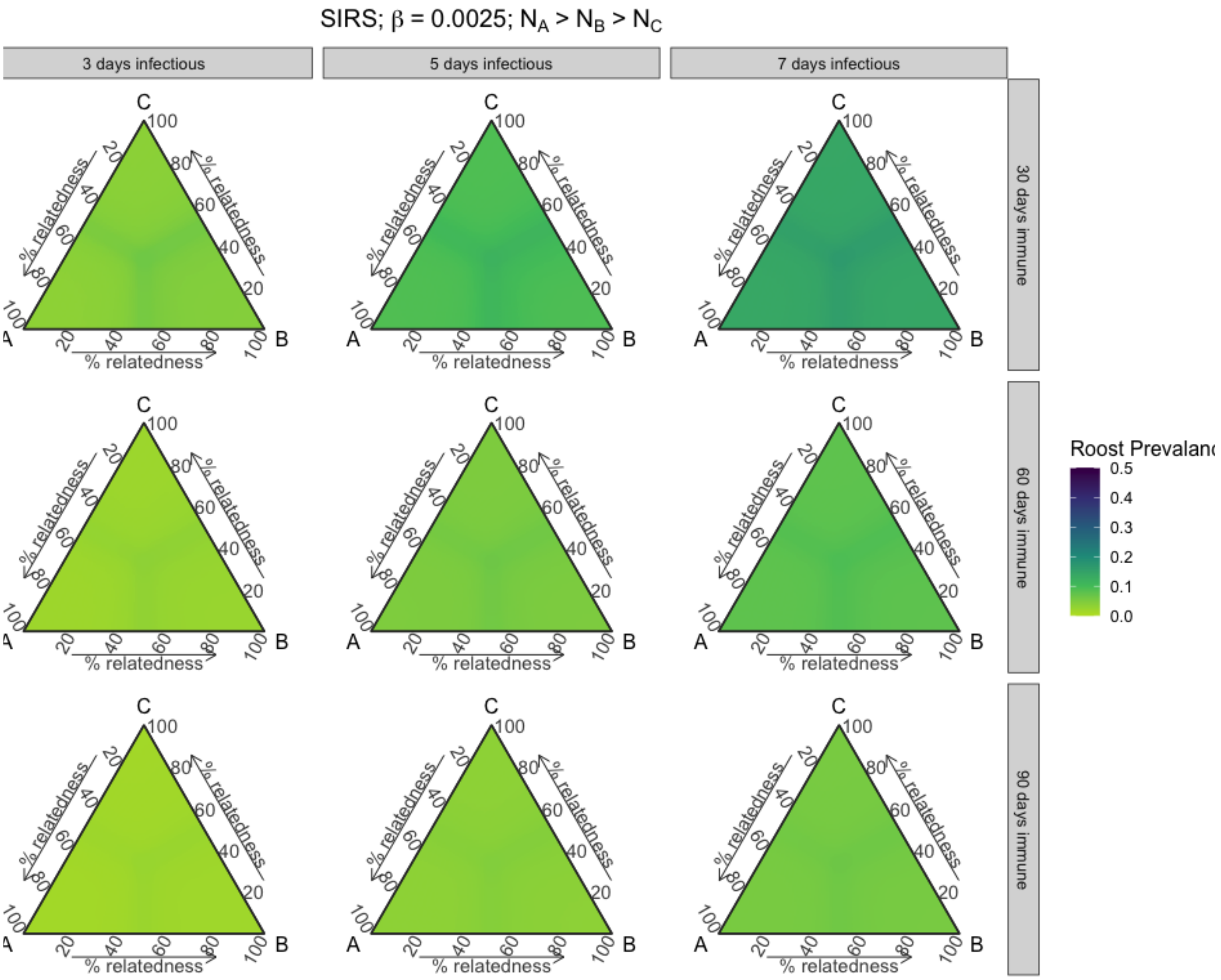


**Figure S24.** Roost prevalence dynamics vary with phylogenetic relatedness between three host species at equilibrium. Results from three-species SIRS models across parameter space for varying short infectious and waning immunity periods, when species starting populations sequentially decrease, and when intraspecific transmission (β) is 0.0025.

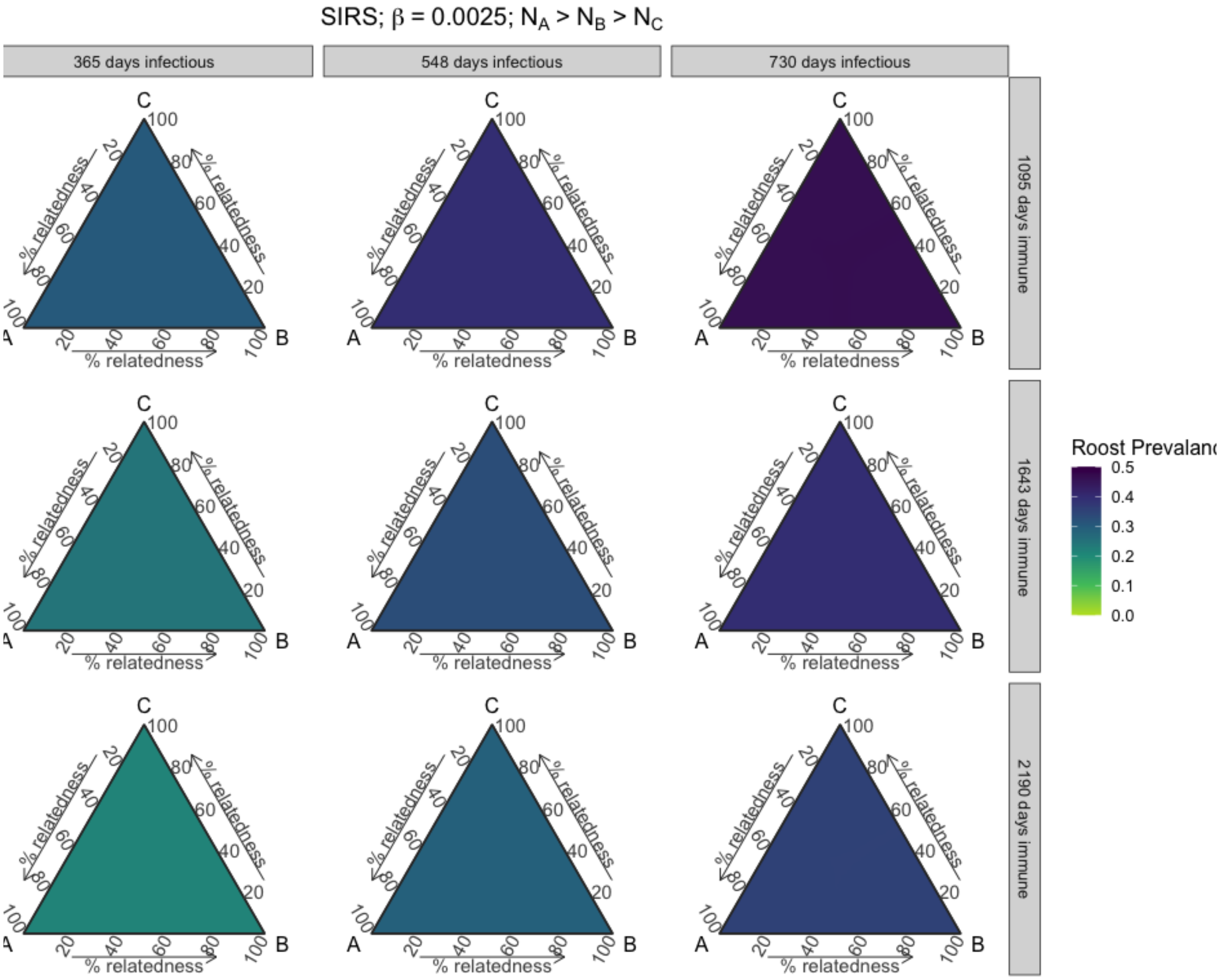


**Figure S25.** Roost prevalence dynamics vary with phylogenetic relatedness between three host species at equilibrium. Results from three-species SIRS models across parameter space for varying long infectious and waning immunity periods, when species starting populations sequentially decrease, and when intraspecific transmission (β) is 0.0025.

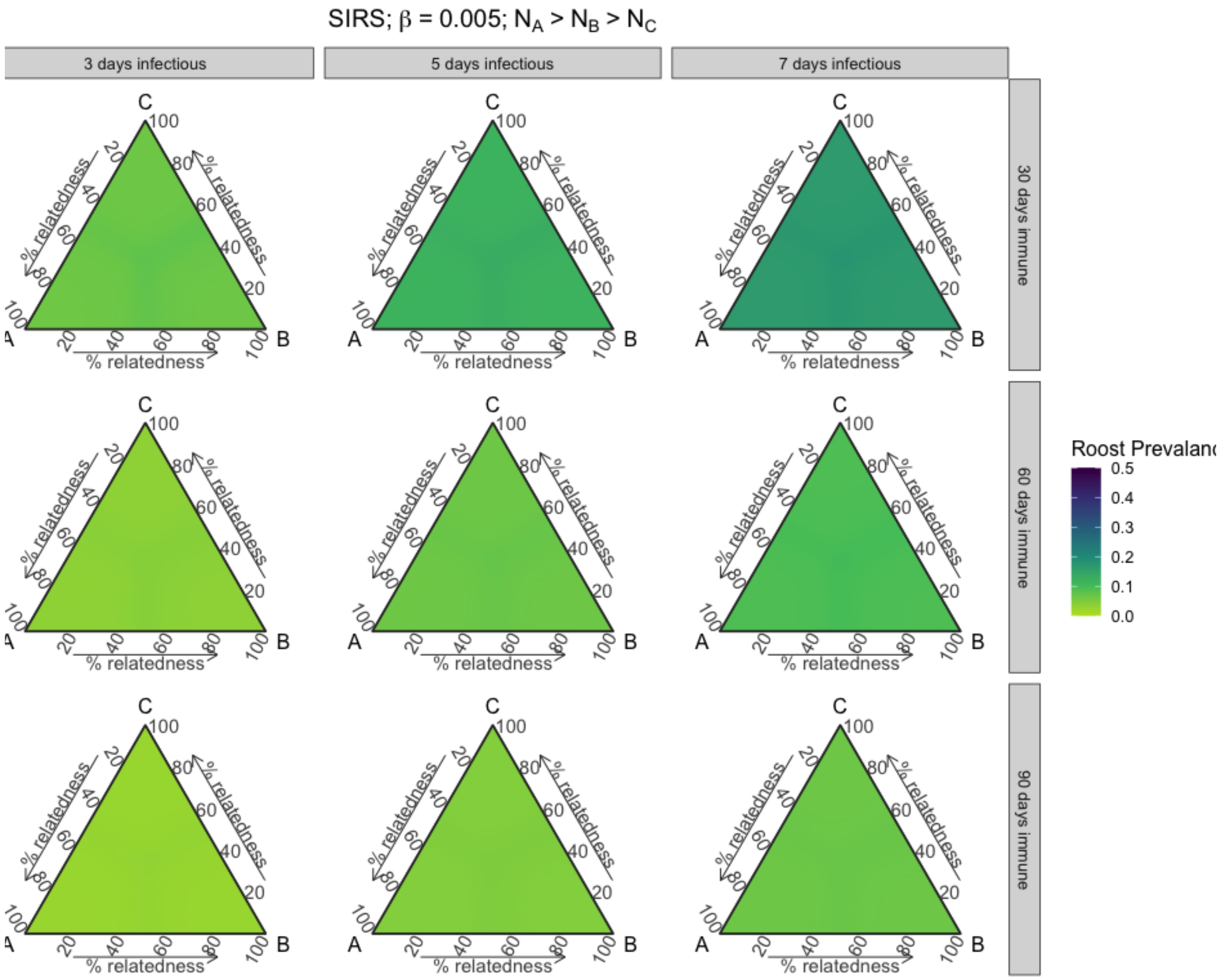


**Figure S26.** Roost prevalence dynamics vary with phylogenetic relatedness between three host species at equilibrium. Results from three-species SIRS models across parameter space for varying short infectious and waning immunity periods, when species starting populations sequentially decrease, and when intraspecific transmission (β) is 0.005.

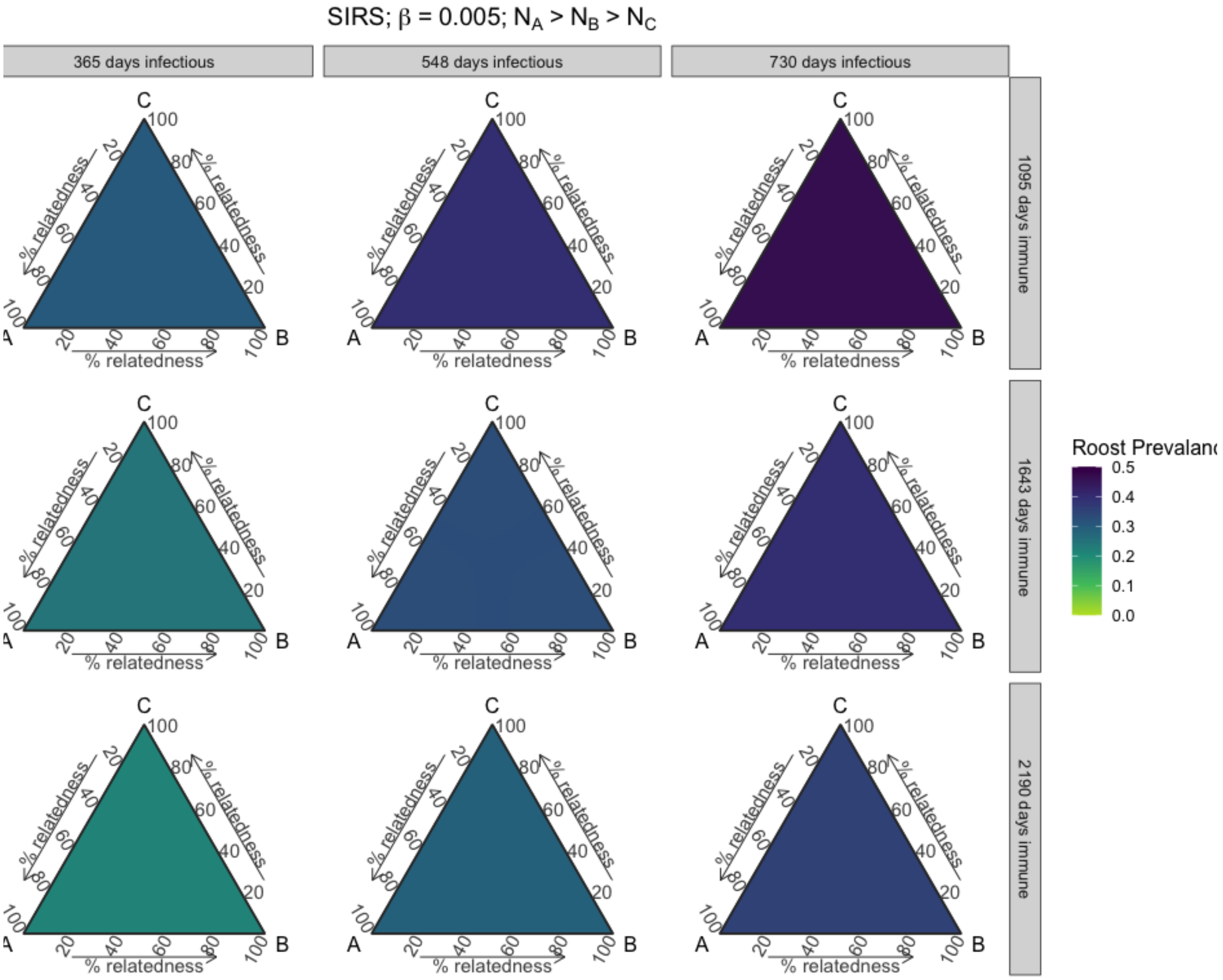


**Figure S27.** Roost prevalence dynamics vary with phylogenetic relatedness between three host species at equilibrium. Results from three-species SIRS models across parameter space for varying long infectious and waning immunity periods, when species starting populations sequentially decrease, and when intraspecific transmission (β) is 0.005.

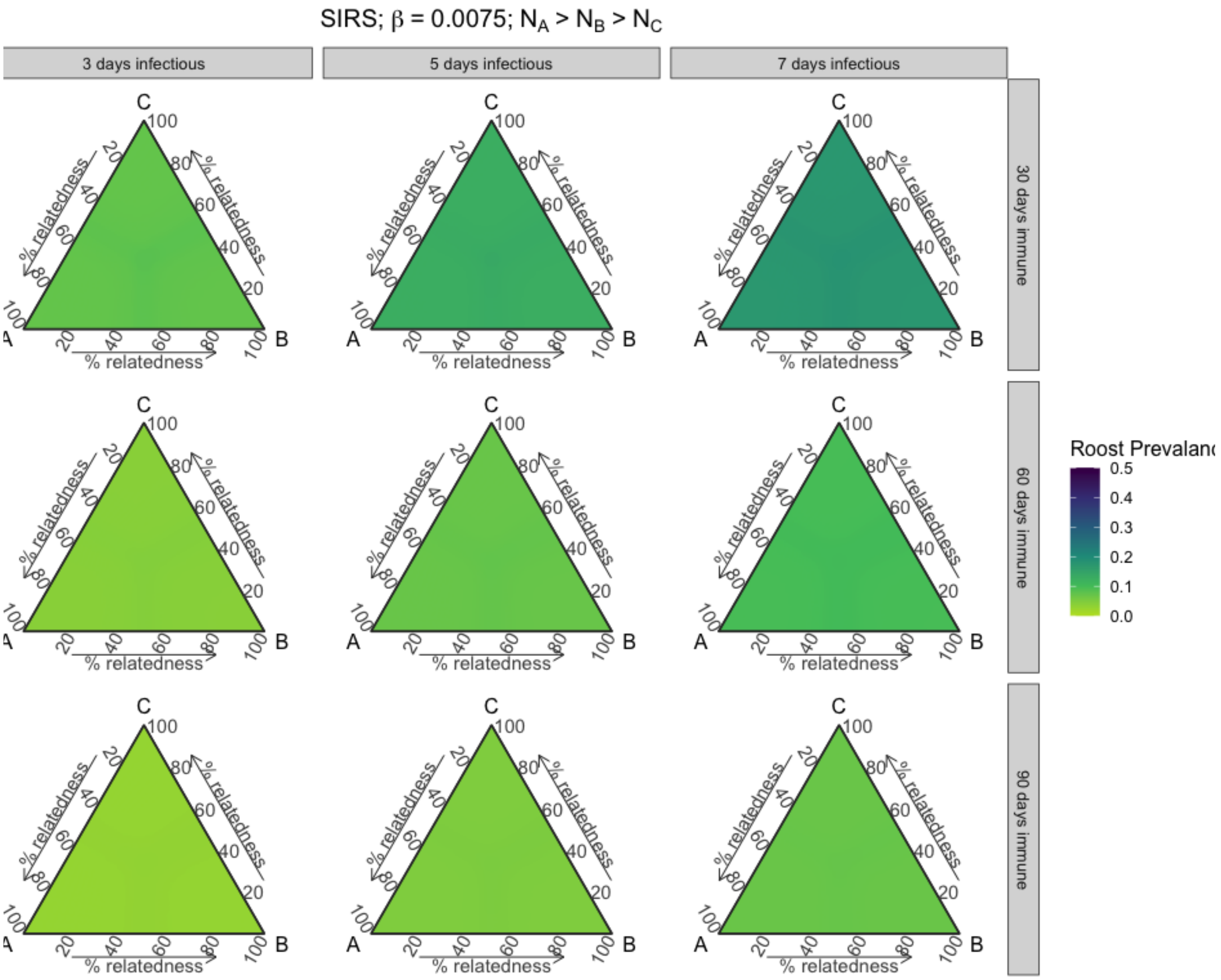


**Figure S28.** Roost prevalence dynamics vary with phylogenetic relatedness between three host species at equilibrium. Results from three-species SIRS models across parameter space for varying short infectious and waning immunity periods, when species starting populations sequentially decrease, and when intraspecific transmission (β) is 0.0075.

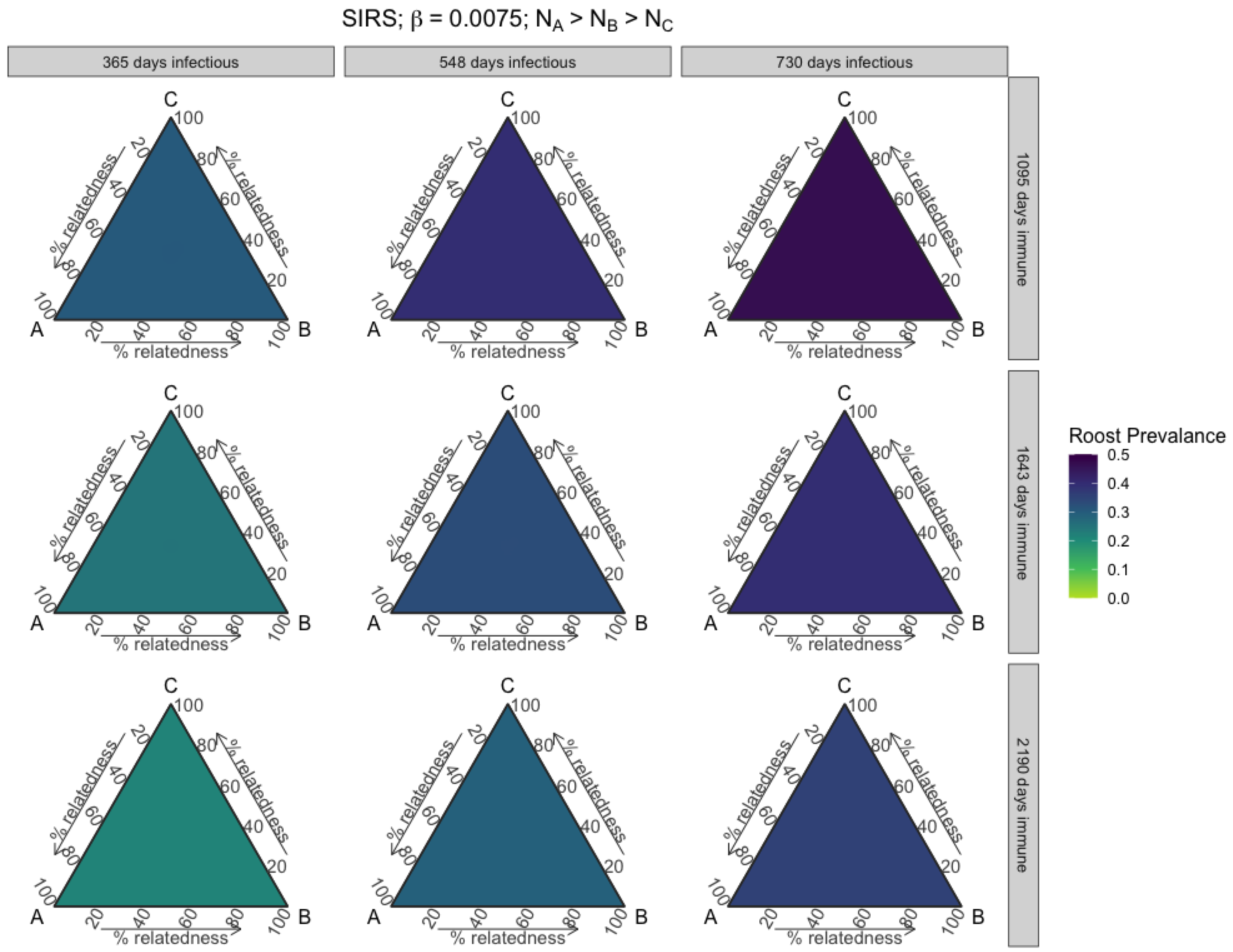


**Figure S29.** Roost prevalence dynamics vary with phylogenetic relatedness between three host species at equilibrium. Results from three-species SIRS models across parameter space for varying long infectious and waning immunity periods, when species starting populations sequentially decrease, and when intraspecific transmission (β) is 0.0075.

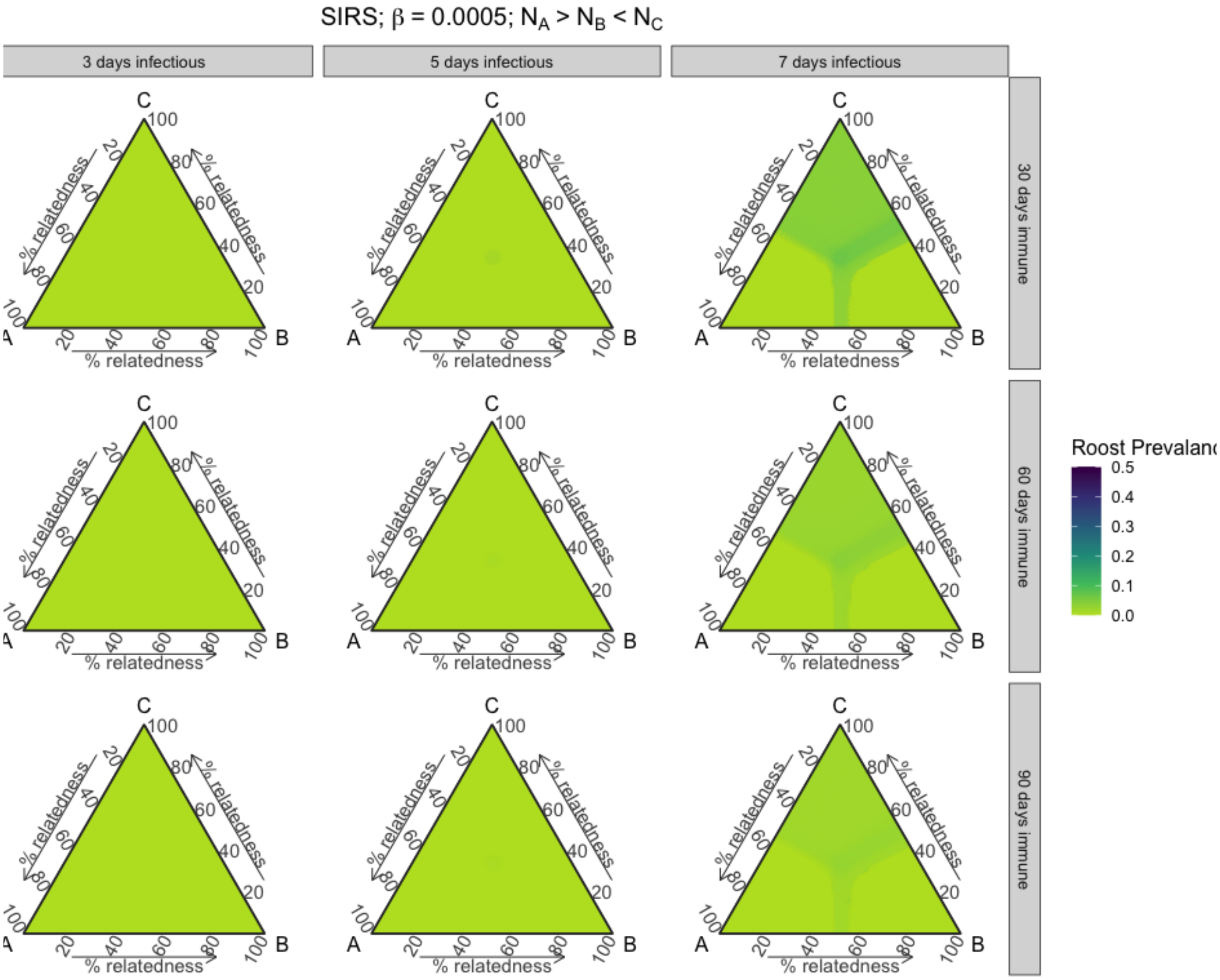


**Figure S30.** Roost prevalence dynamics vary with phylogenetic relatedness between three host species at equilibrium. Results from three-species SIRS models across parameter space for varying short infectious and waning immunity periods, when starting populations of species A is greater than both species B and C and species C is greater than B, and when intraspecific transmission (β) is 0.0005.

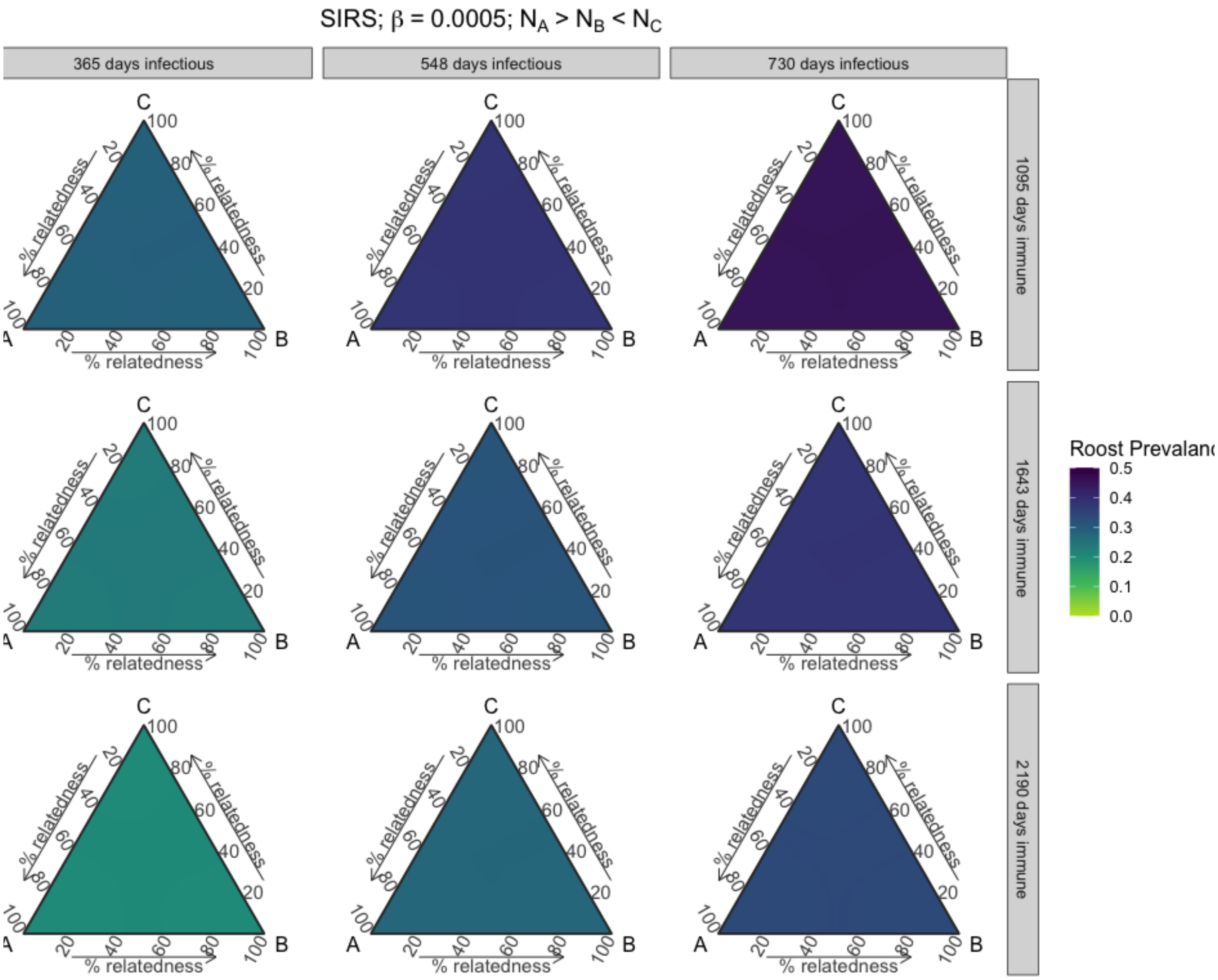


**Figure S31.** Roost prevalence dynamics vary with phylogenetic relatedness between three host species at equilibrium. Results from three-species SIRS models across parameter space for varying long infectious and waning immunity periods, when starting populations of species A is greater than both species B and C and species C is greater than B, and when intraspecific transmission (β) is 0.0005.

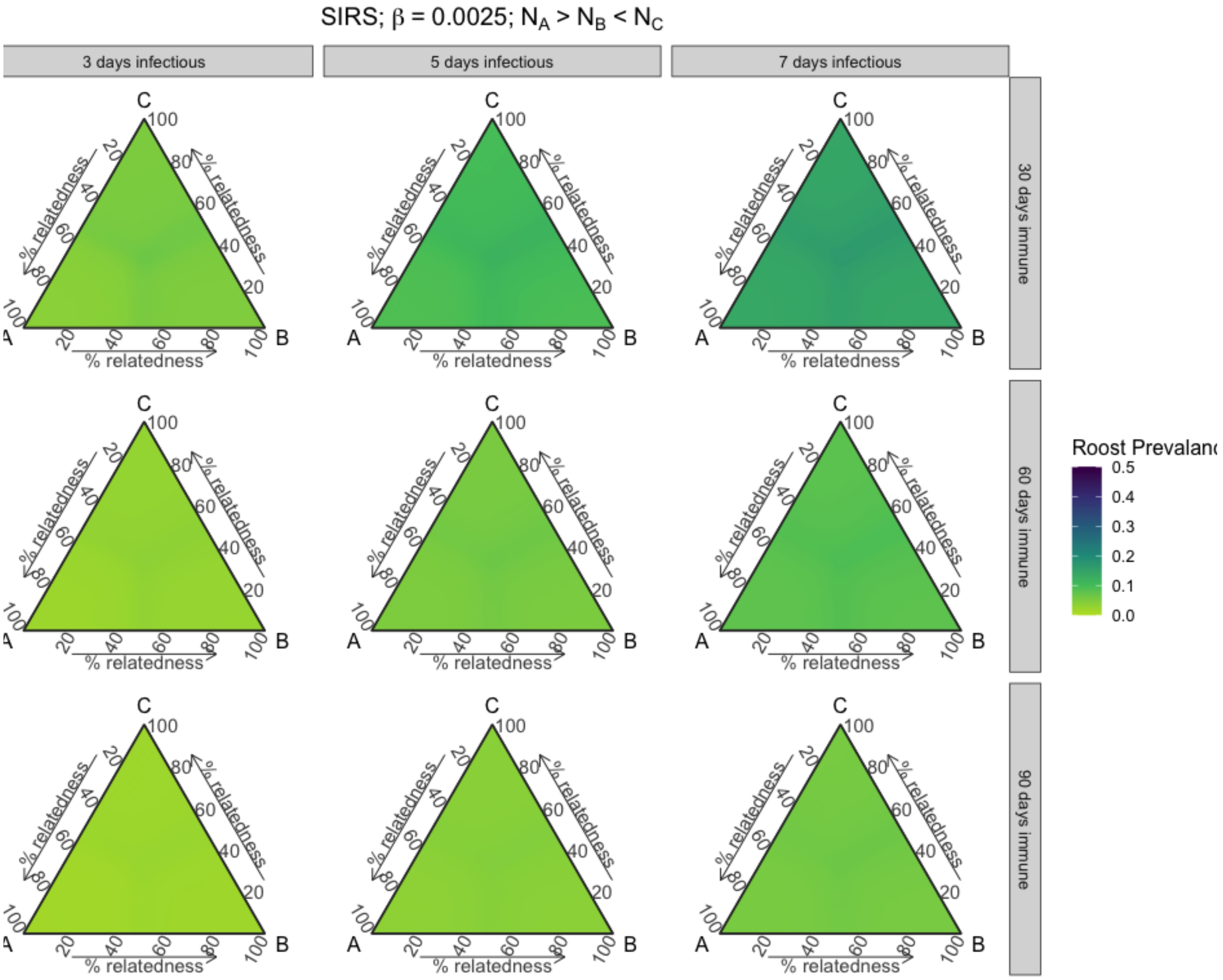


**Figure S32.** Roost prevalence dynamics vary with phylogenetic relatedness between three host species at equilibrium. Results from three-species SIRS models across parameter space for varying short infectious and waning immunity periods, when starting populations of species A is greater than both species B and C and species C is greater than B, and when intraspecific transmission (β) is 0.0025.

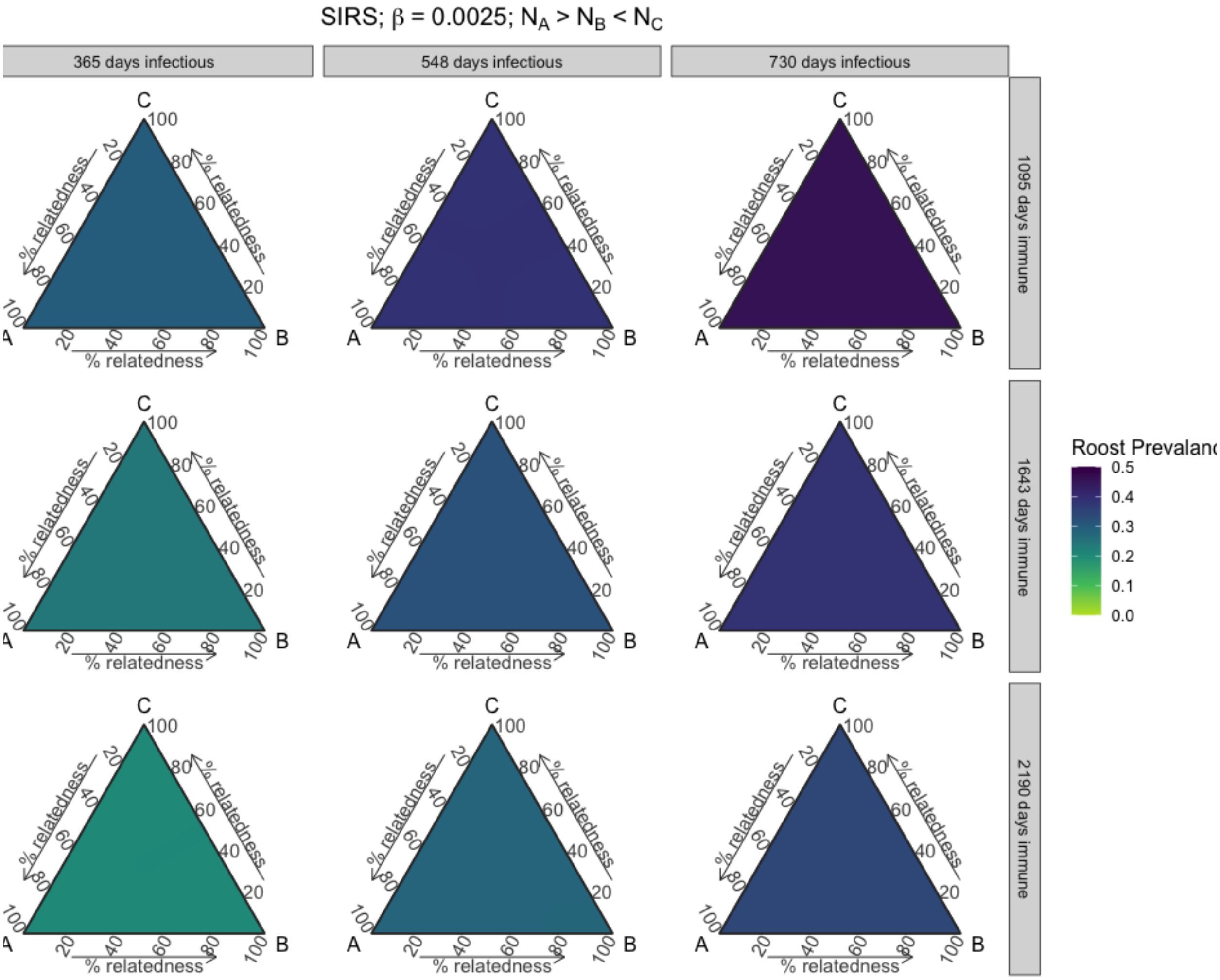


**Figure S33.** Roost prevalence dynamics vary with phylogenetic relatedness between three host species at equilibrium. Results from three-species SIRS models across parameter space for varying long infectious and waning immunity periods, when starting populations of species A is greater than both species B and C and species C is greater than B, and when intraspecific transmission (β) is 0.0025.

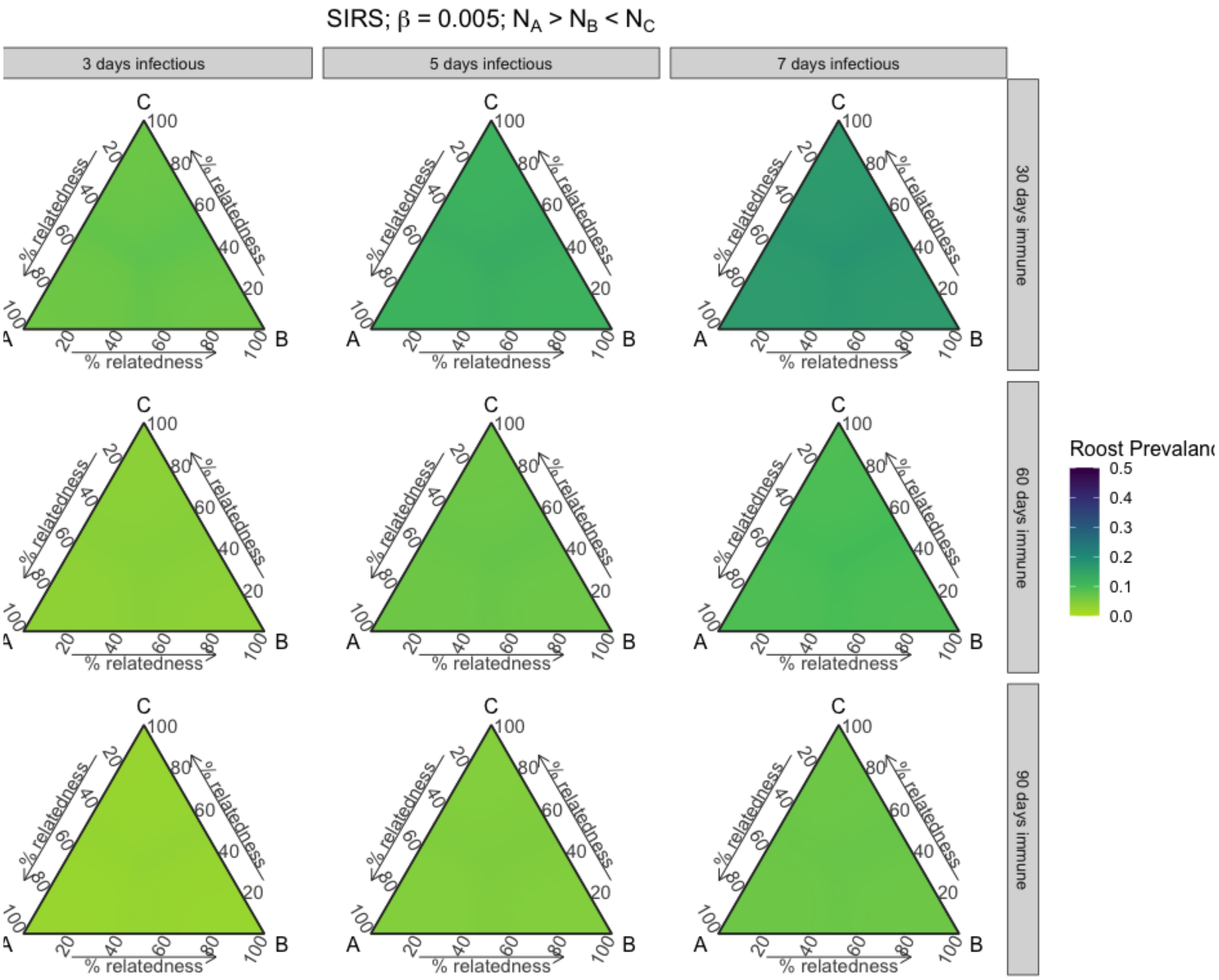


**Figure S34.** Roost prevalence dynamics vary with phylogenetic relatedness between three host species at equilibrium. Results from three-species SIRS models across parameter space for varying short infectious and waning immunity periods, when starting populations of species A is greater than both species B and C and species C is greater than B, and when intraspecific transmission (β) is 0.005.

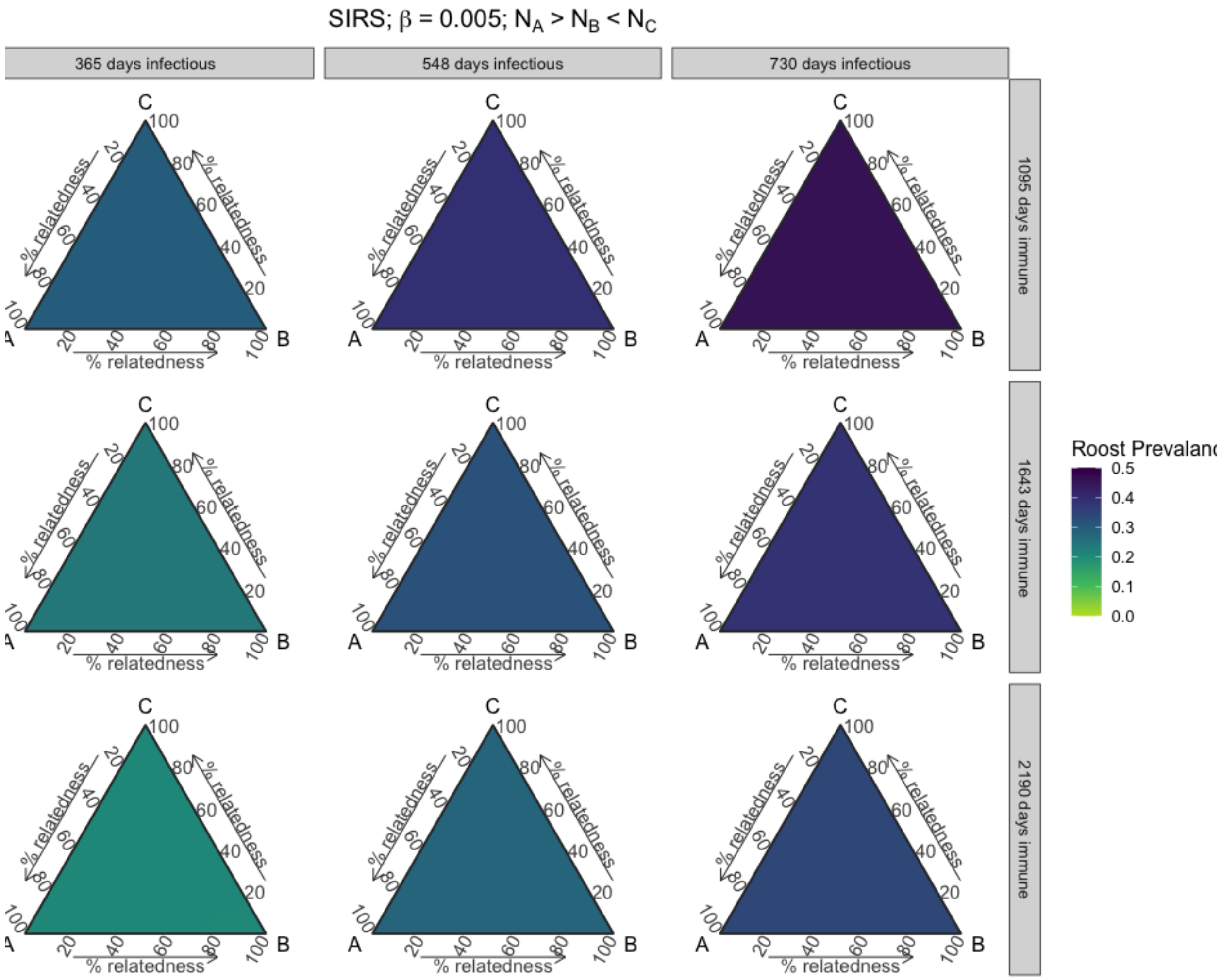


**Figure S35.** Roost prevalence dynamics vary with phylogenetic relatedness between three host species at equilibrium. Results from three-species SIRS models across parameter space for varying long infectious and waning immunity periods, when starting populations of species A is greater than both species B and C and species C is greater than B, and when intraspecific transmission (β) is 0.005.

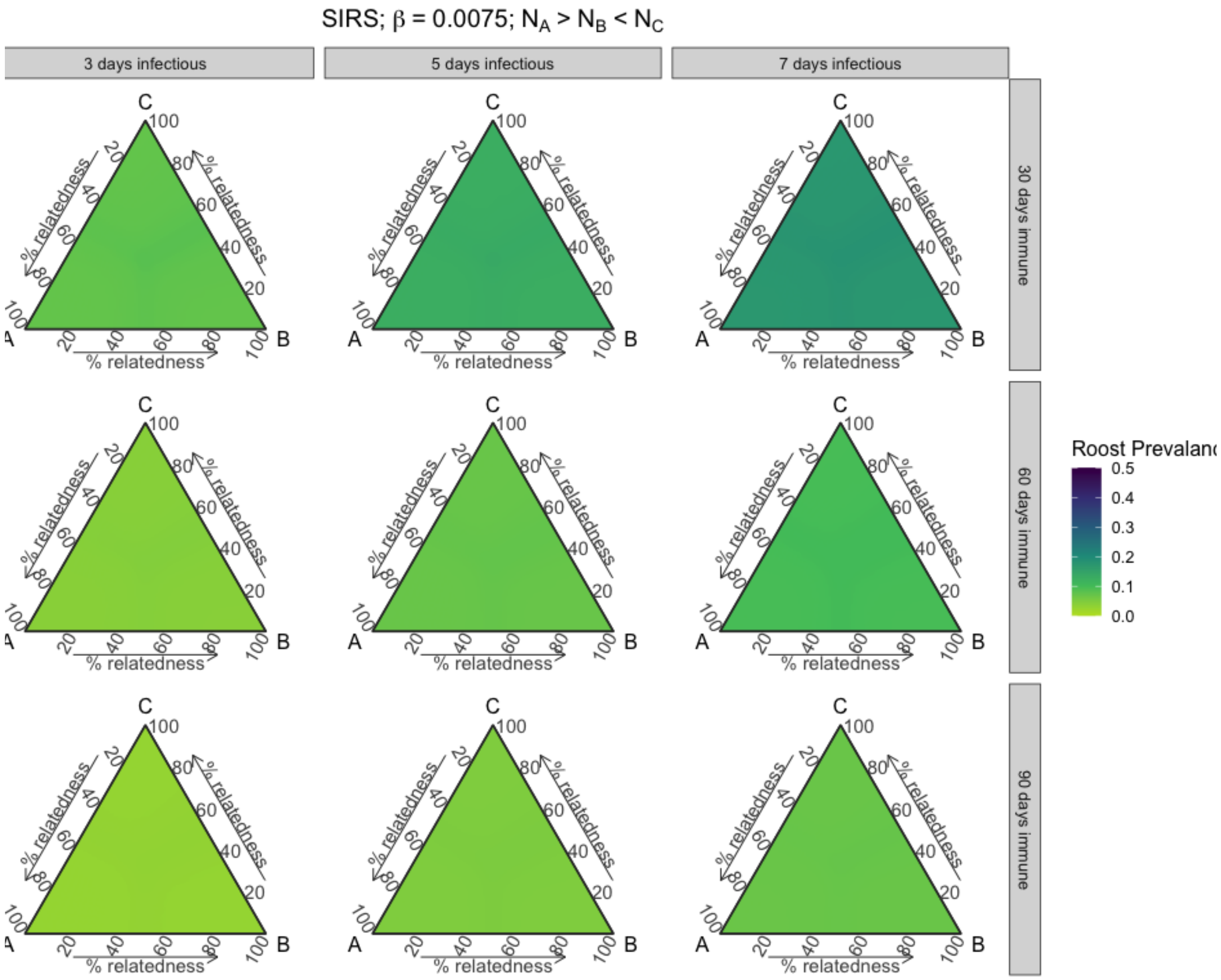


**Figure S36.** Roost prevalence dynamics vary with phylogenetic relatedness between three host species at equilibrium. Results from three-species SIRS models across parameter space for varying short infectious and waning immunity periods, when starting populations of species A is greater than both species B and C and species C is greater than B, and when intraspecific transmission (β) is 0.0075.

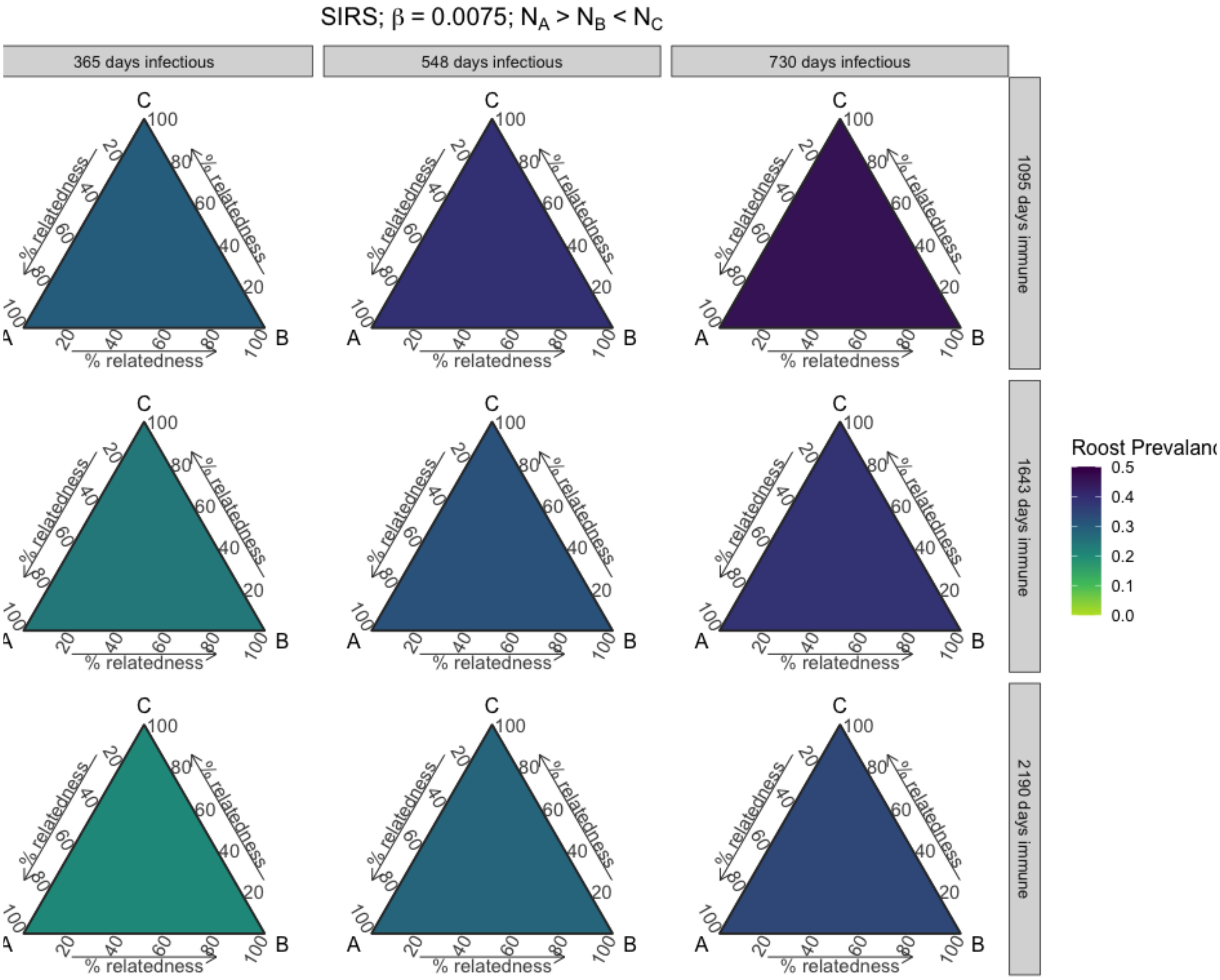


**Figure S37.** Roost prevalence dynamics vary with phylogenetic relatedness between three host species at equilibrium. Results from three-species SIRS models across parameter space for varying long infectious and waning immunity periods, when starting populations of species A is greater than both species B and C and species C is greater than B, and when intraspecific transmission (β) is 0.0075.

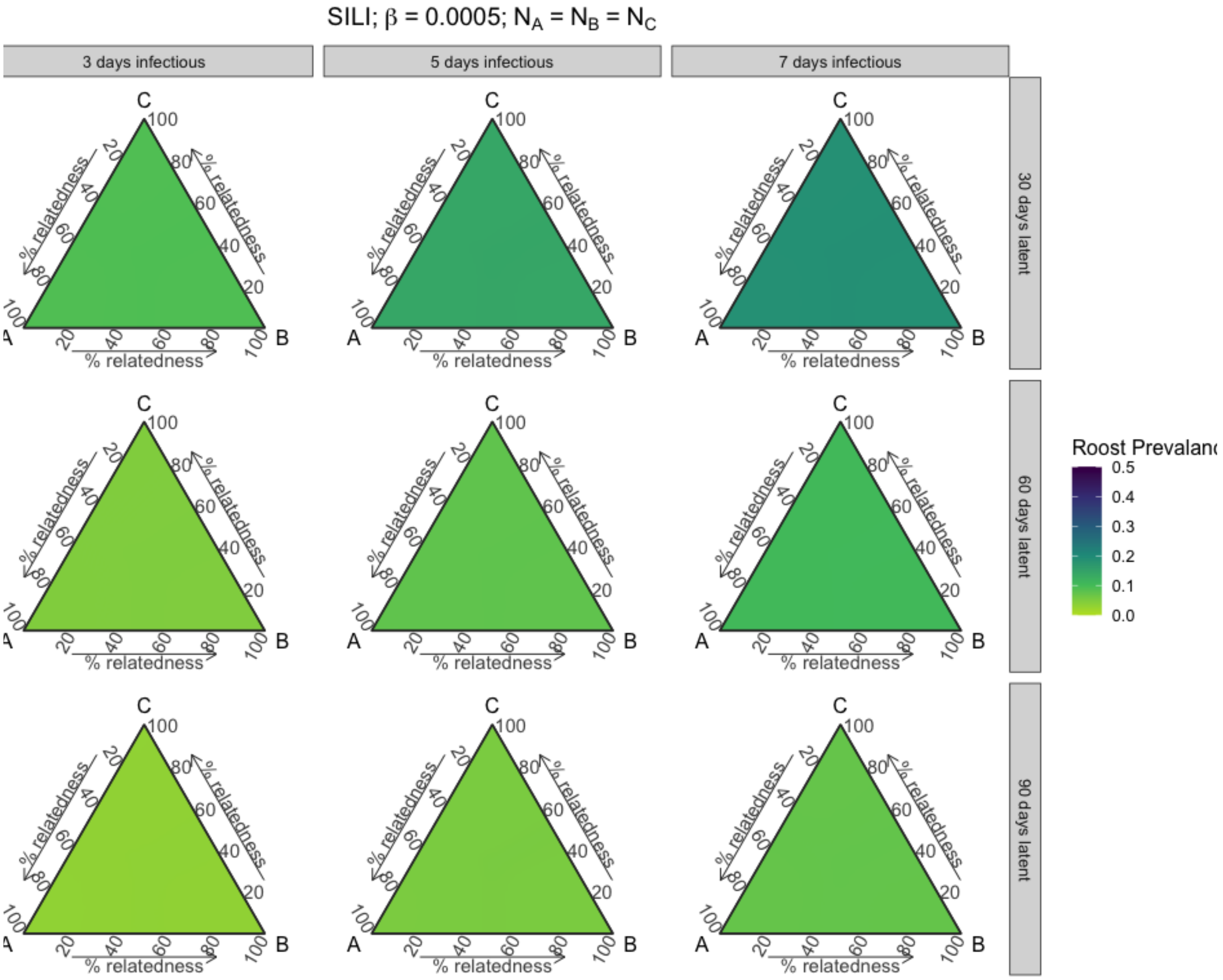


**Figure S38.** Roost prevalence dynamics vary with phylogenetic relatedness between three host species at equilibrium. Results from three-species SILI models across parameter space for varying short infectious and waning immunity periods, when species starting populations are equal, and when intraspecific transmission (β) is 0.0005.

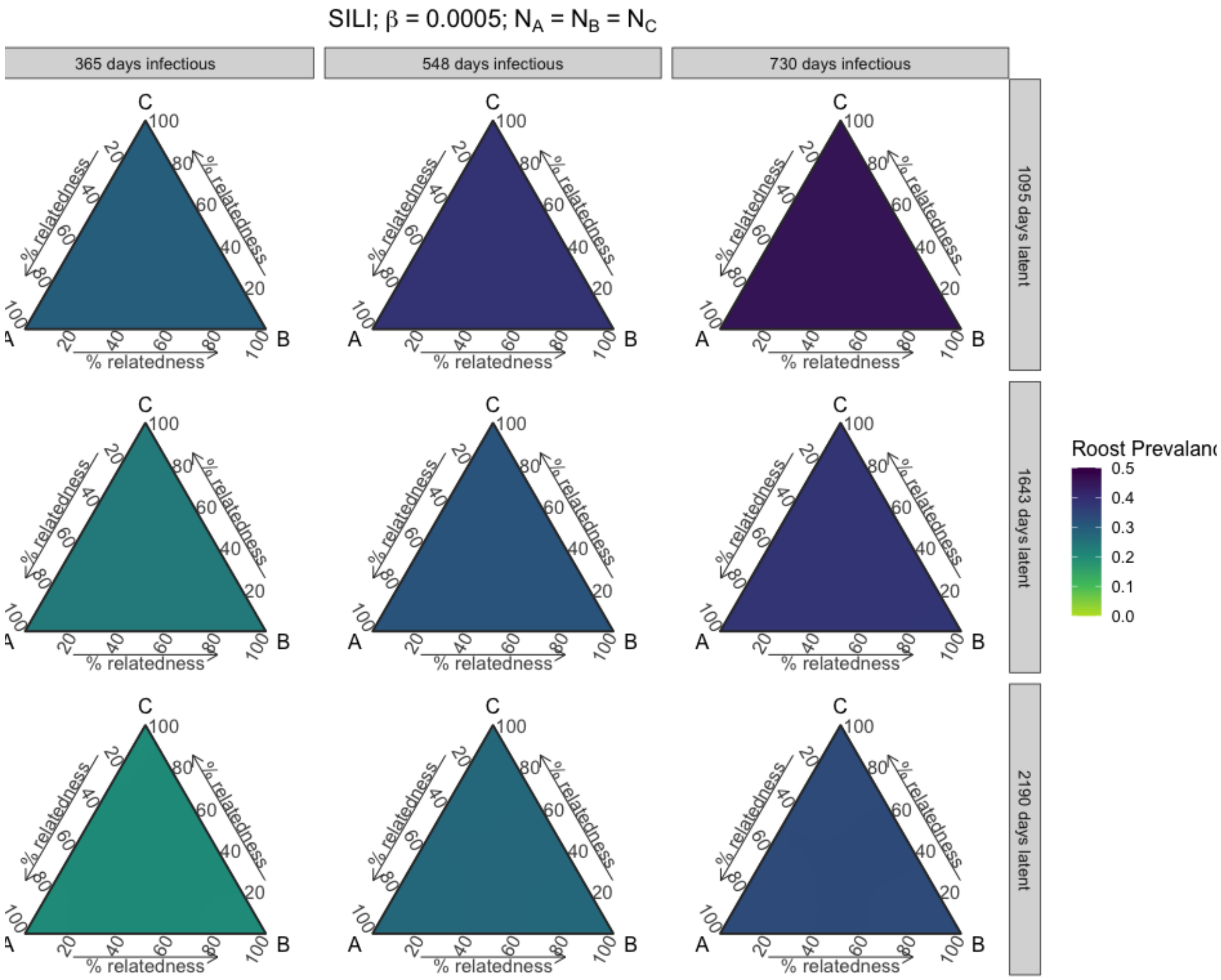


**Figure S39.** Roost prevalence dynamics vary with phylogenetic relatedness between three host species at equilibrium. Results from three-species SILI models across parameter space for varying long infectious and waning immunity periods, when species starting populations are equal, and when intraspecific transmission (β) is 0.0005.

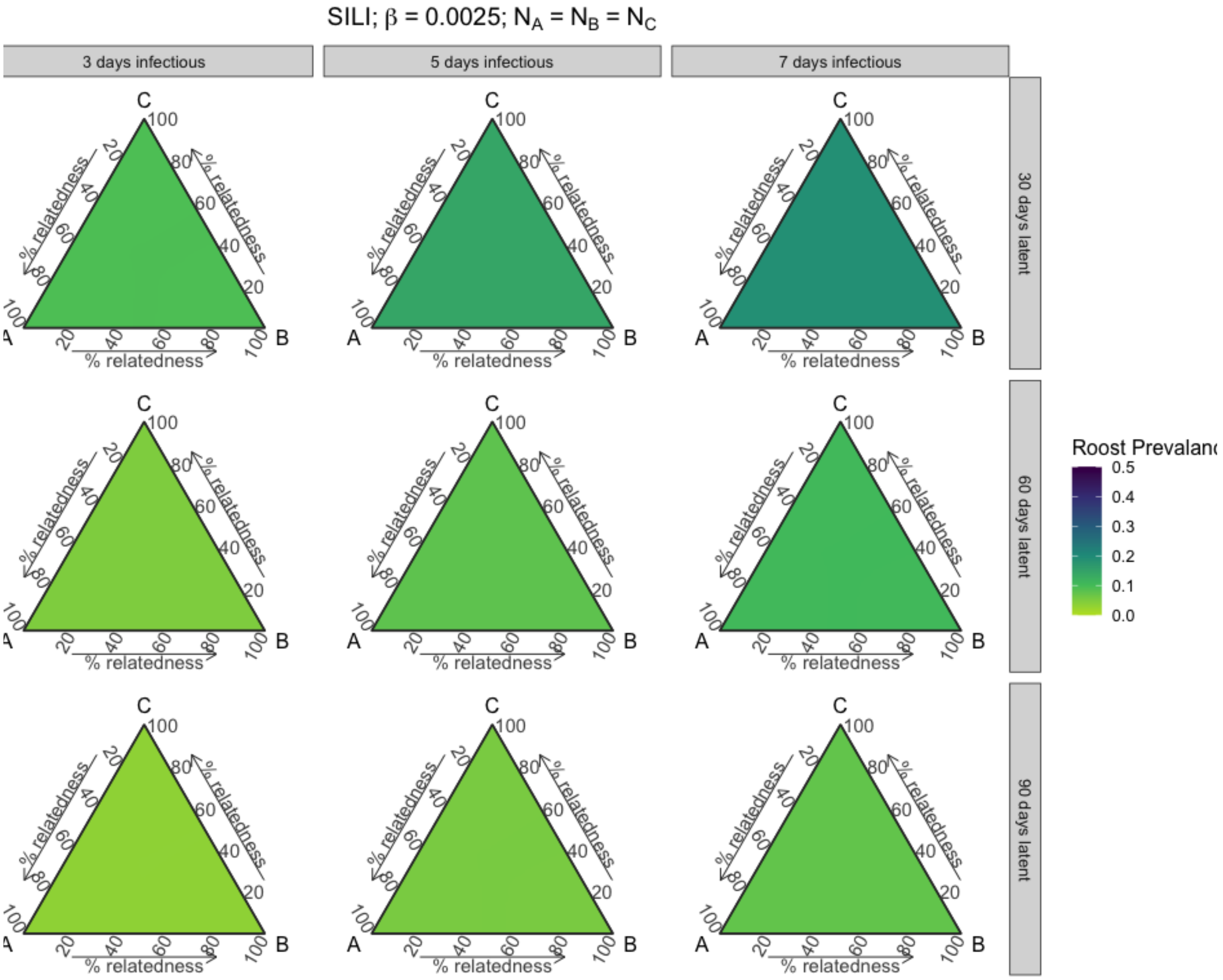


**Figure S40.** Roost prevalence dynamics vary with phylogenetic relatedness between three host species at equilibrium. Results from three-species SILI models across parameter space for varying short infectious and waning immunity periods, when species starting populations are equal, and when intraspecific transmission (β) is 0.0025.

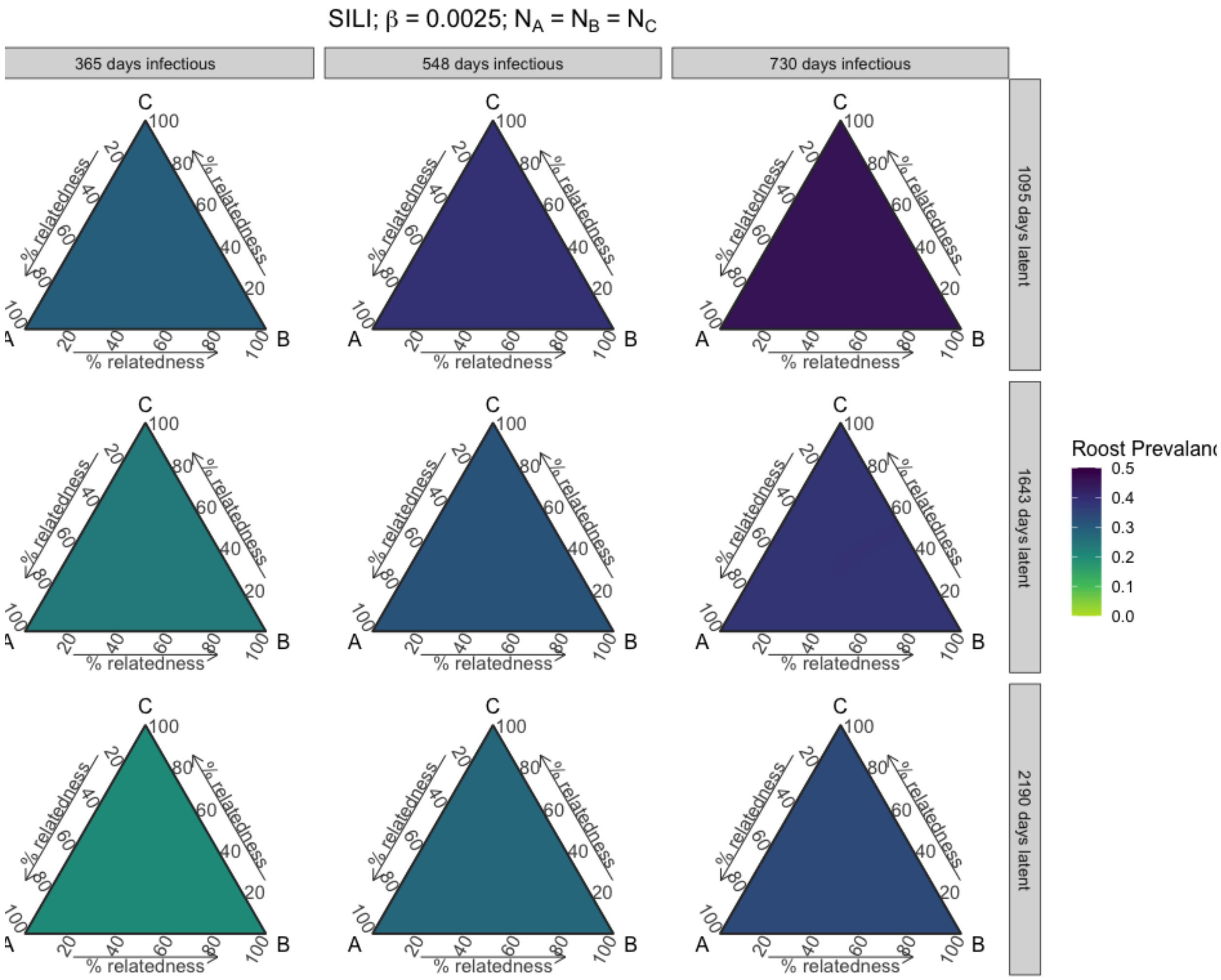


**Figure S41.** Roost prevalence dynamics vary with phylogenetic relatedness between three host species at equilibrium. Results from three-species SILI models across parameter space for varying long infectious and waning immunity periods, when species starting populations are equal, and when intraspecific transmission (β) is 0.0025.

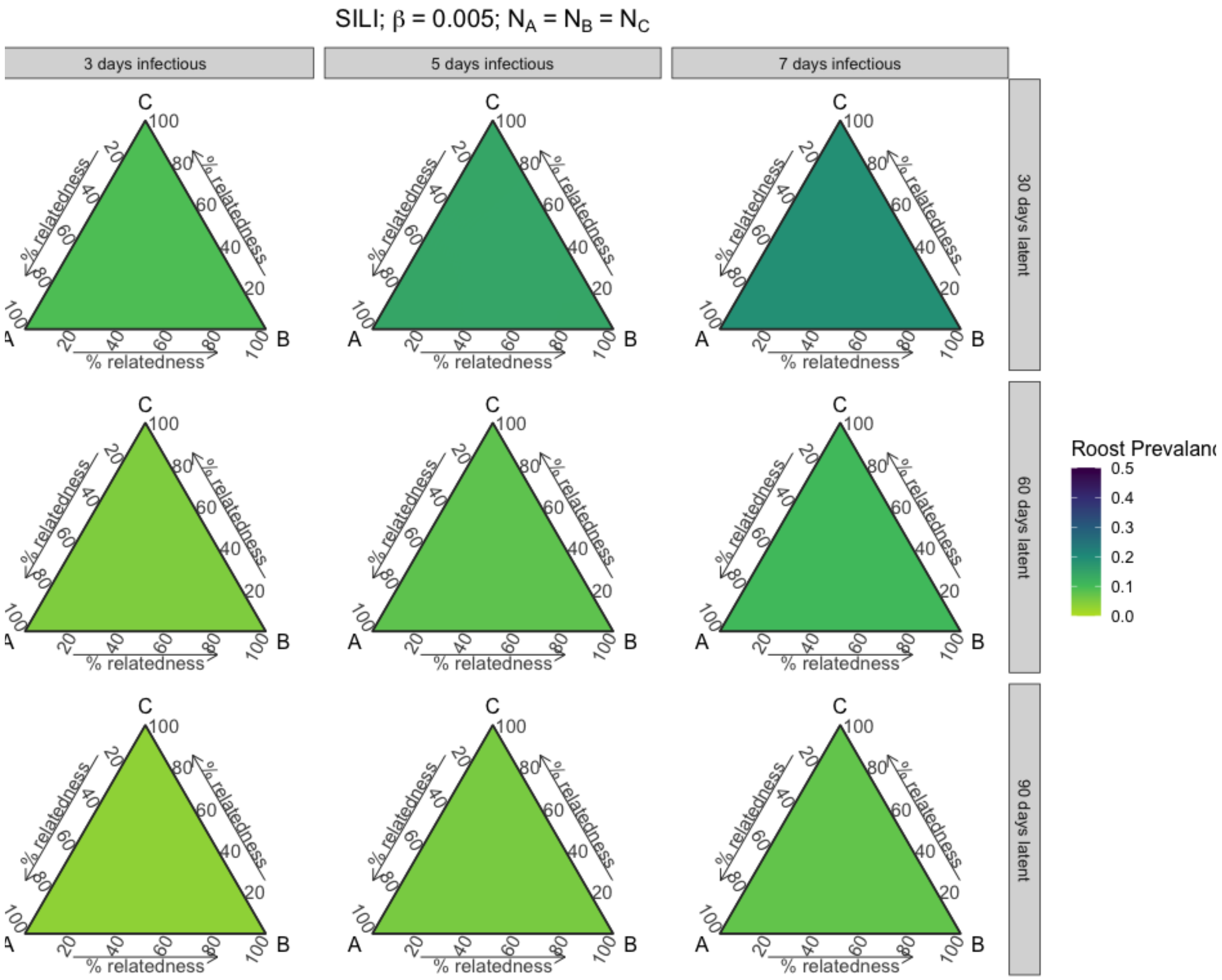


**Figure S42.** Roost prevalence dynamics vary with phylogenetic relatedness between three host species at equilibrium. Results from three-species SILI models across parameter space for varying short infectious and waning immunity periods, when species starting populations are equal, and when intraspecific transmission (β) is 0.005.

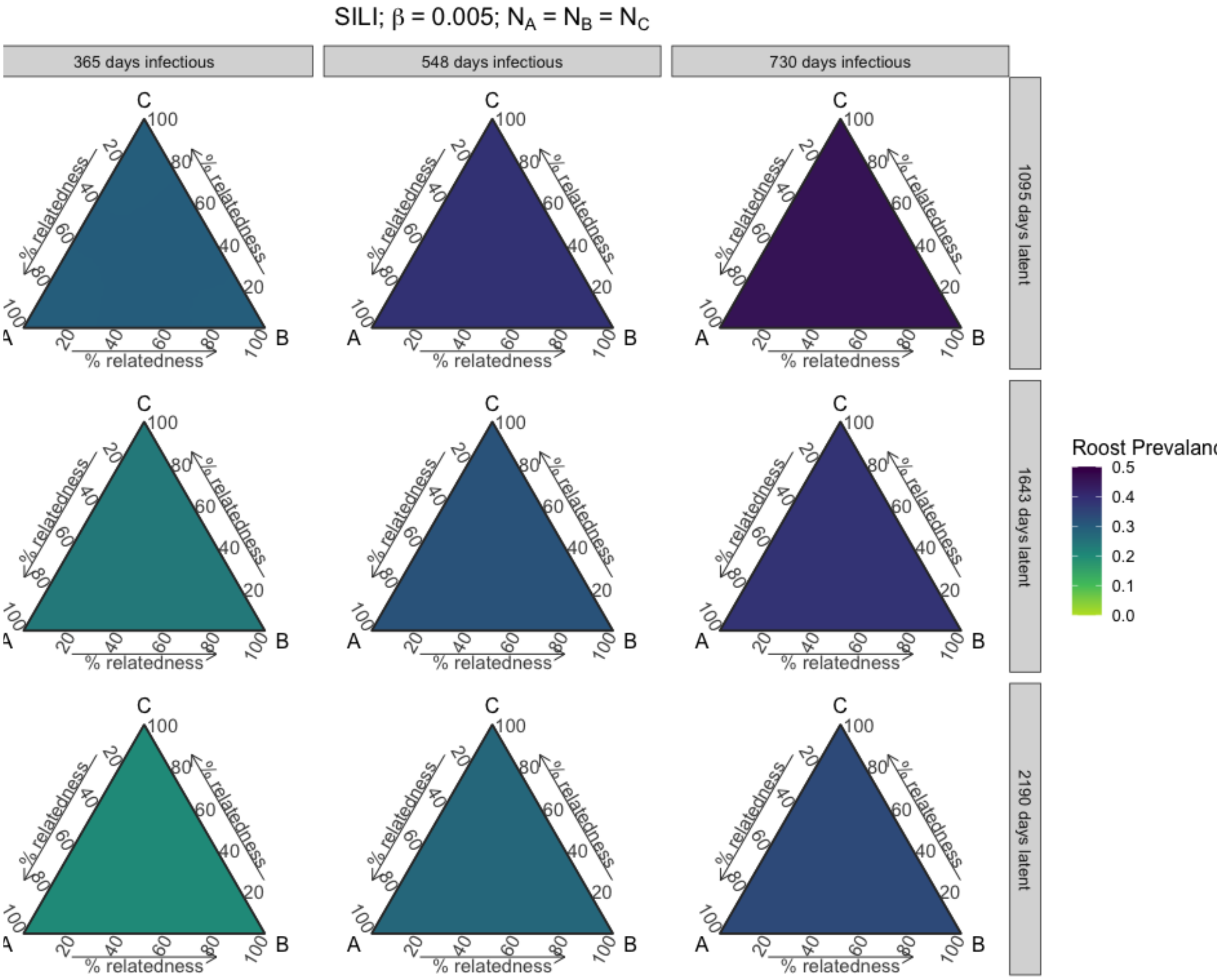


**Figure S43.** Roost prevalence dynamics vary with phylogenetic relatedness between three host species at equilibrium. Results from three-species SILI models across parameter space for varying long infectious and waning immunity periods, when species starting populations are equal, and when intraspecific transmission (β) is 0.005.

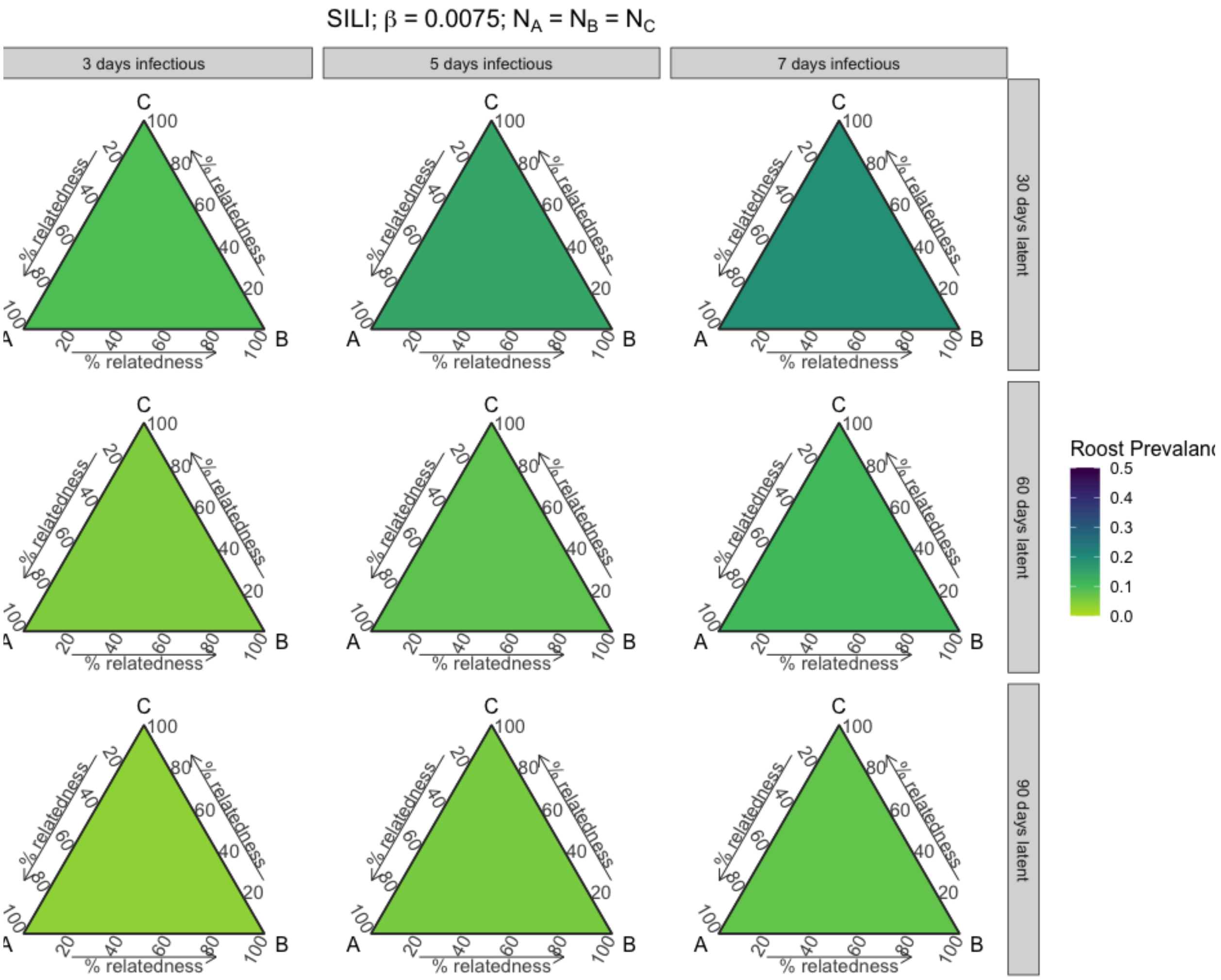


**Figure S44.** Roost prevalence dynamics vary with phylogenetic relatedness between three host species at equilibrium. Results from three-species SILI models across parameter space for varying short infectious and waning immunity periods, when species starting populations are equal, and when intraspecific transmission (β) is 0.0075.

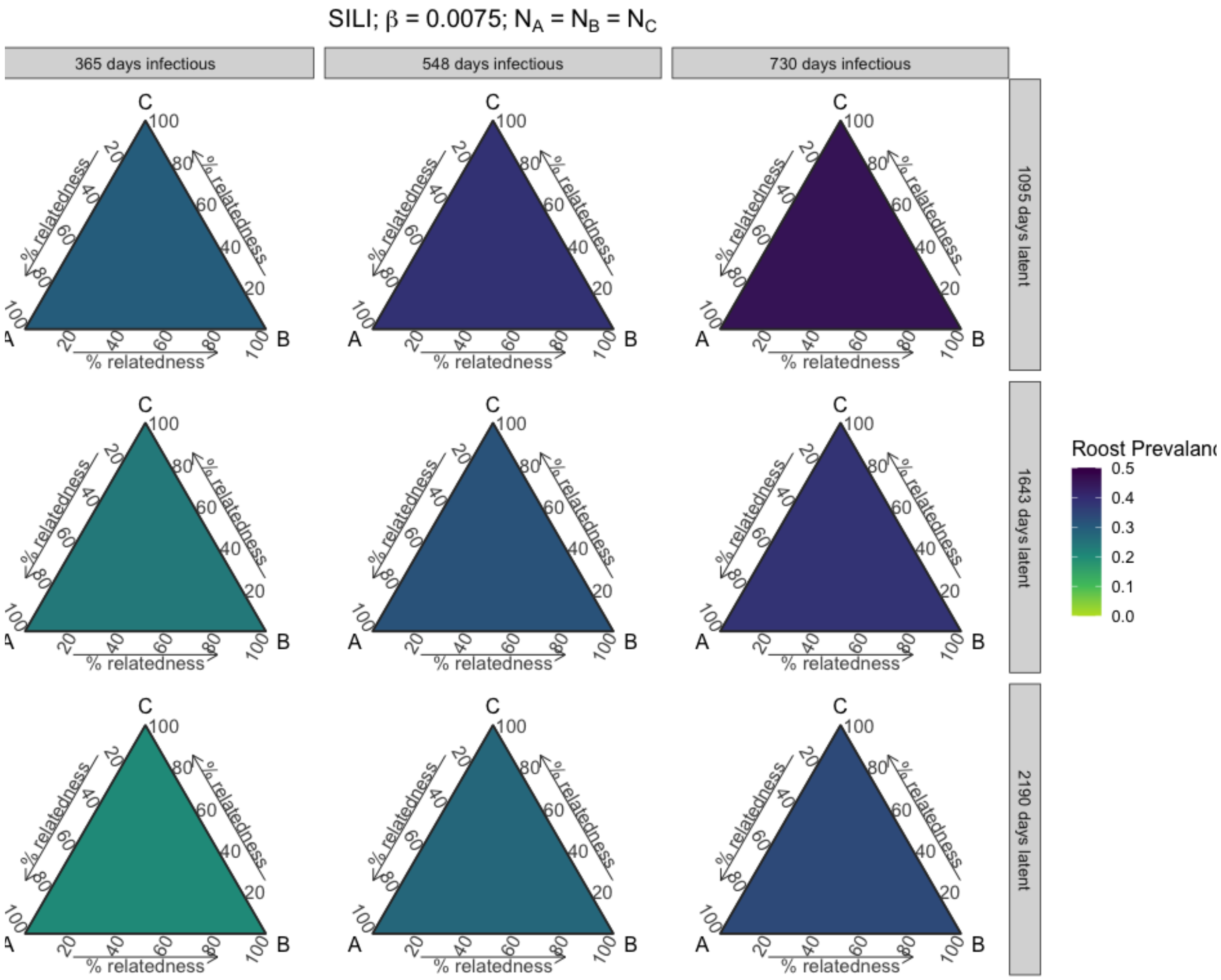


**Figure S45.** Roost prevalence dynamics vary with phylogenetic relatedness between three host species at equilibrium. Results from three-species SILI models across parameter space for varying long infectious and waning immunity periods, when species starting populations are equal, and when intraspecific transmission (β) is 0.0075.

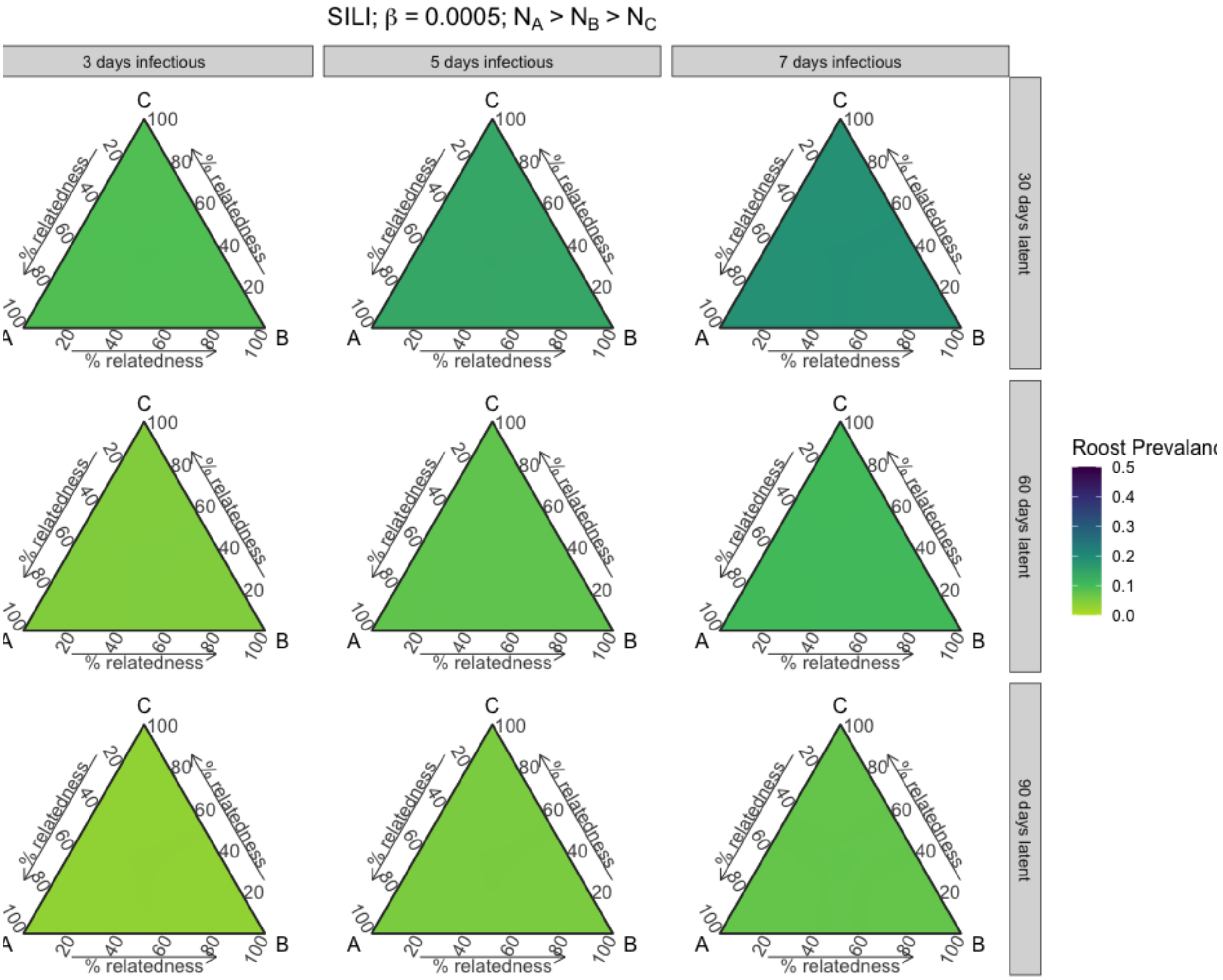


**Figure S46.** Roost prevalence dynamics vary with phylogenetic relatedness between three host species at equilibrium. Results from three-species SILI models across parameter space for varying short infectious and waning immunity periods, when species starting populations sequentially decrease, and when intraspecific transmission (β) is 0.0005.

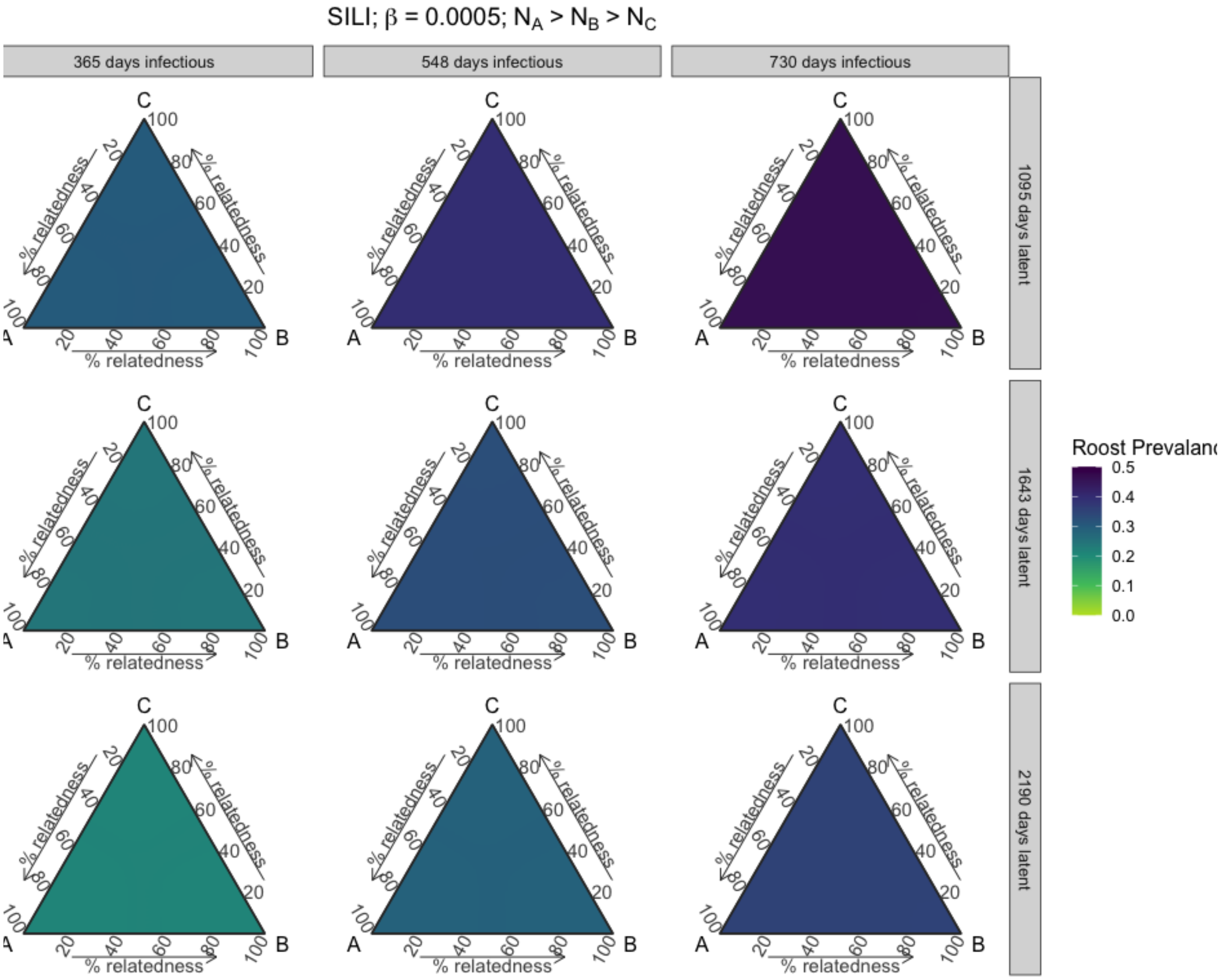


**Figure S47.** Roost prevalence dynamics vary with phylogenetic relatedness between three host species at equilibrium. Results from three-species SILI models across parameter space for varying long infectious and waning immunity periods, when species starting populations sequentially decrease, and when intraspecific transmission (β) is 0.0005.

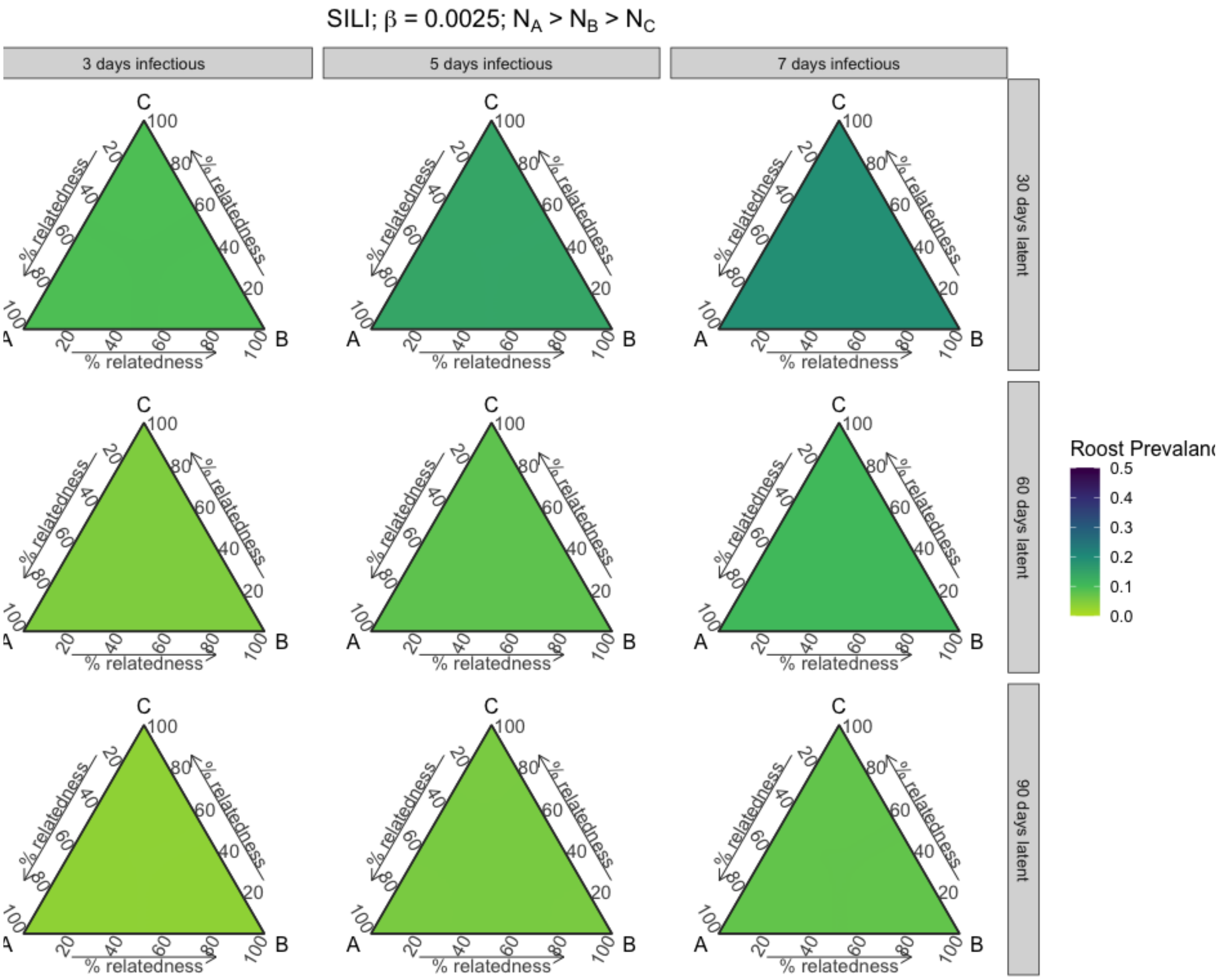


**Figure S48.** Roost prevalence dynamics vary with phylogenetic relatedness between three host species at equilibrium. Results from three-species SILI models across parameter space for varying short infectious and waning immunity periods, when species starting populations sequentially decrease, and when intraspecific transmission (β) is 0.0025.

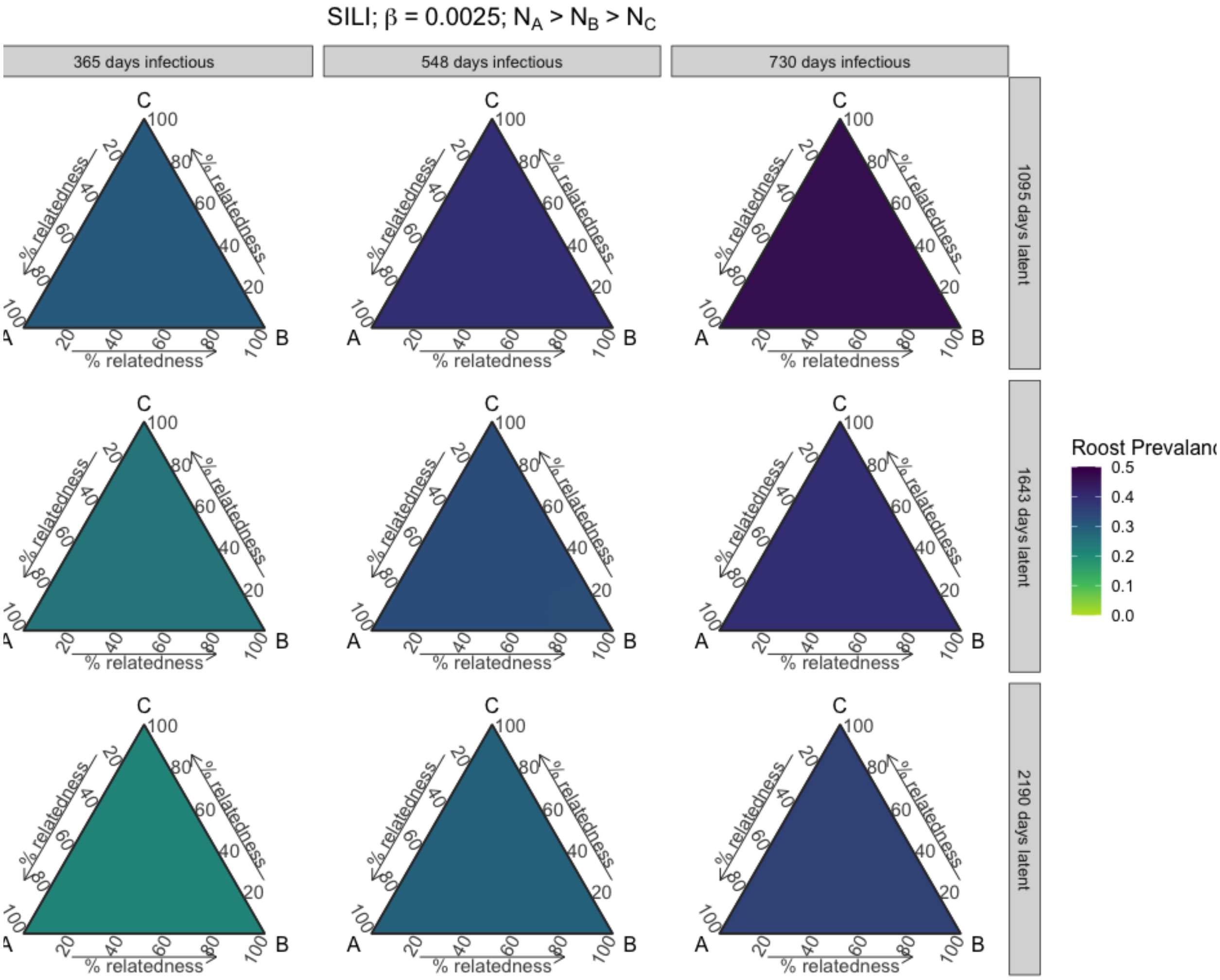


**Figure S49.** Roost prevalence dynamics vary with phylogenetic relatedness between three host species at equilibrium. Results from three-species SILI models across parameter space for varying long infectious and waning immunity periods, when species starting populations sequentially decrease, and when intraspecific transmission (β) is 0.0025.

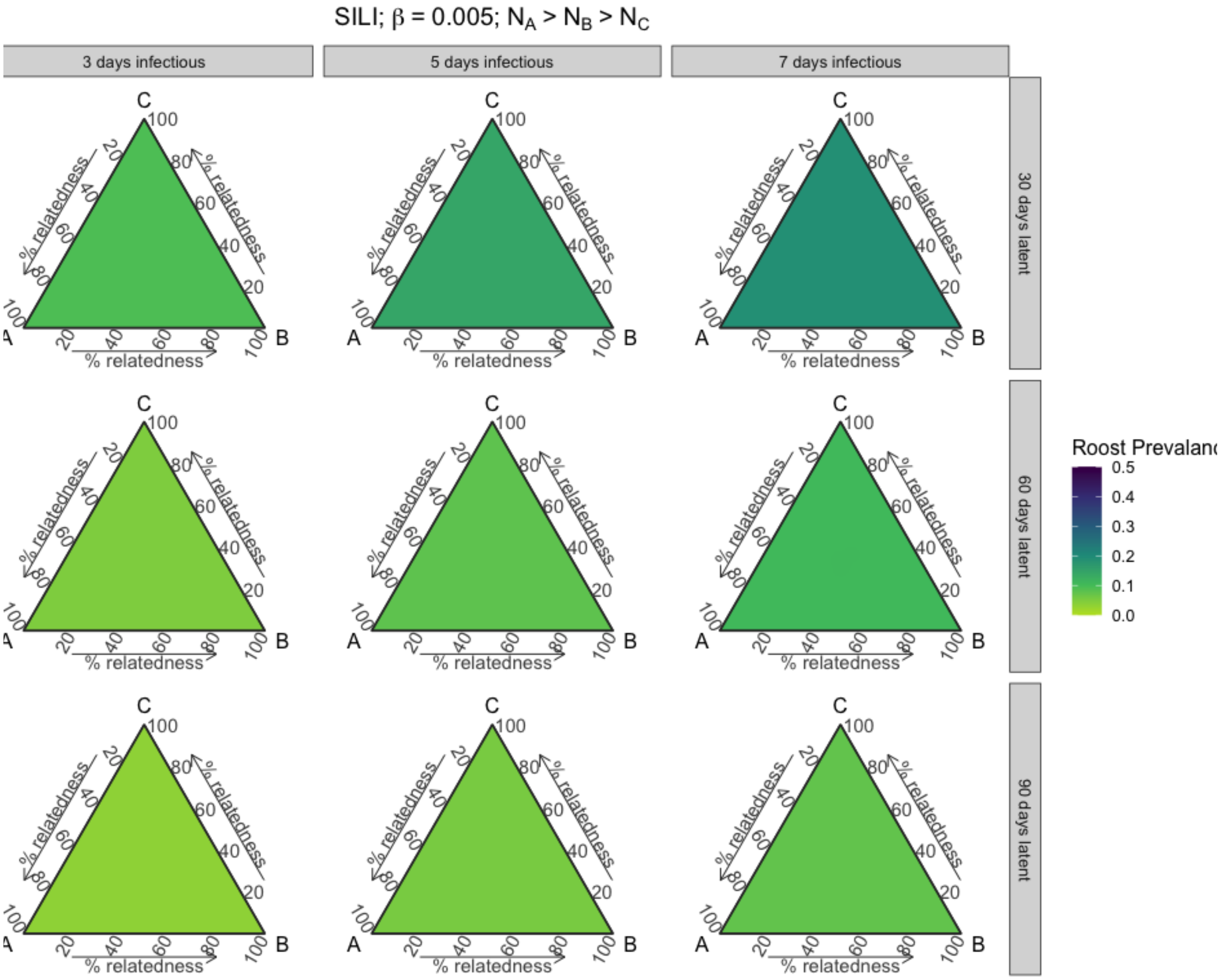


**Figure S50.** Roost prevalence dynamics vary with phylogenetic relatedness between three host species at equilibrium. Results from three-species SILI models across parameter space for varying short infectious and waning immunity periods, when species starting populations sequentially decrease, and when intraspecific transmission (β) is 0.005.

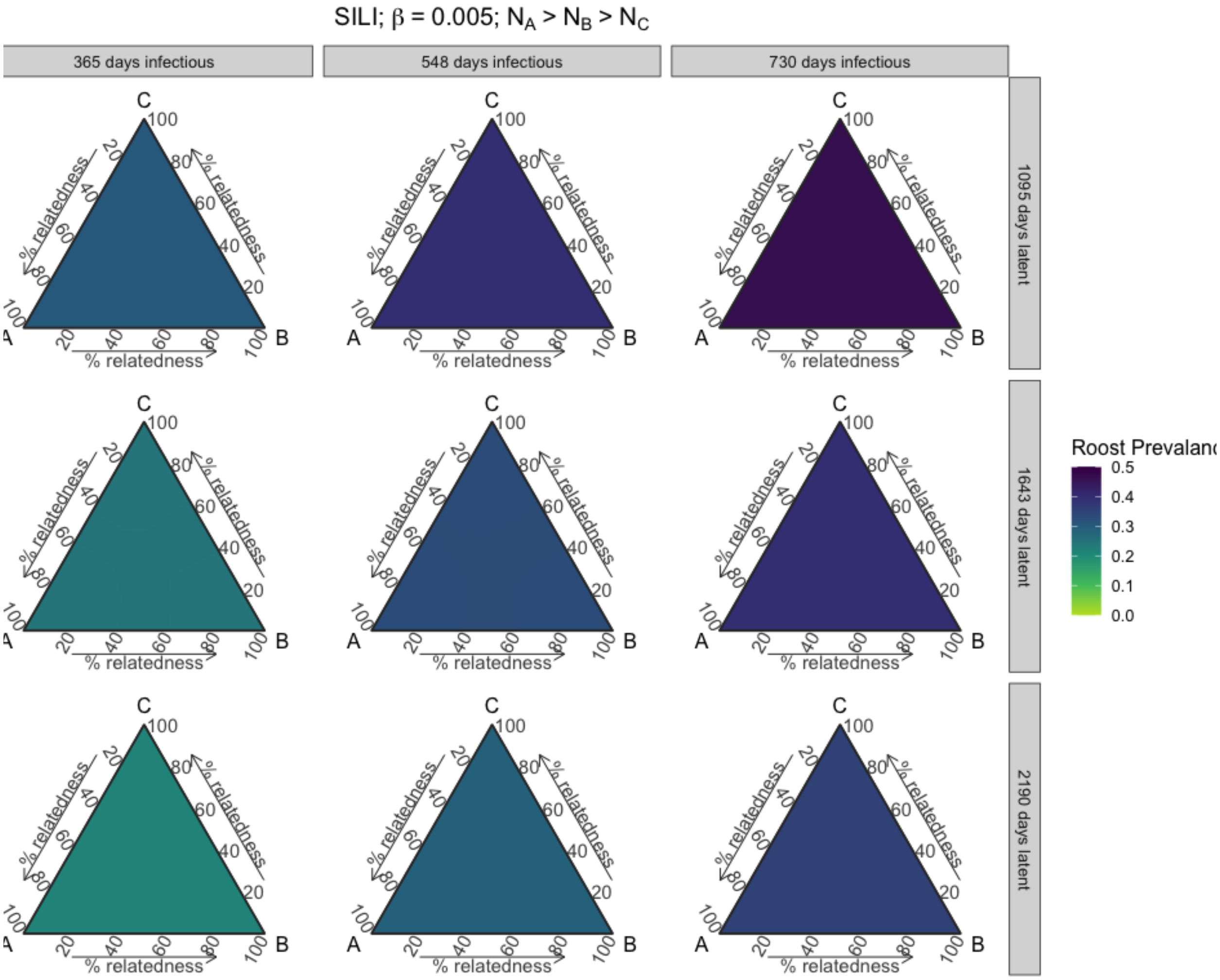


**Figure S51.** Roost prevalence dynamics vary with phylogenetic relatedness between three host species at equilibrium. Results from three-species SILI models across parameter space for varying long infectious and waning immunity periods, when species starting populations sequentially decrease, and when intraspecific transmission (β) is 0.005.

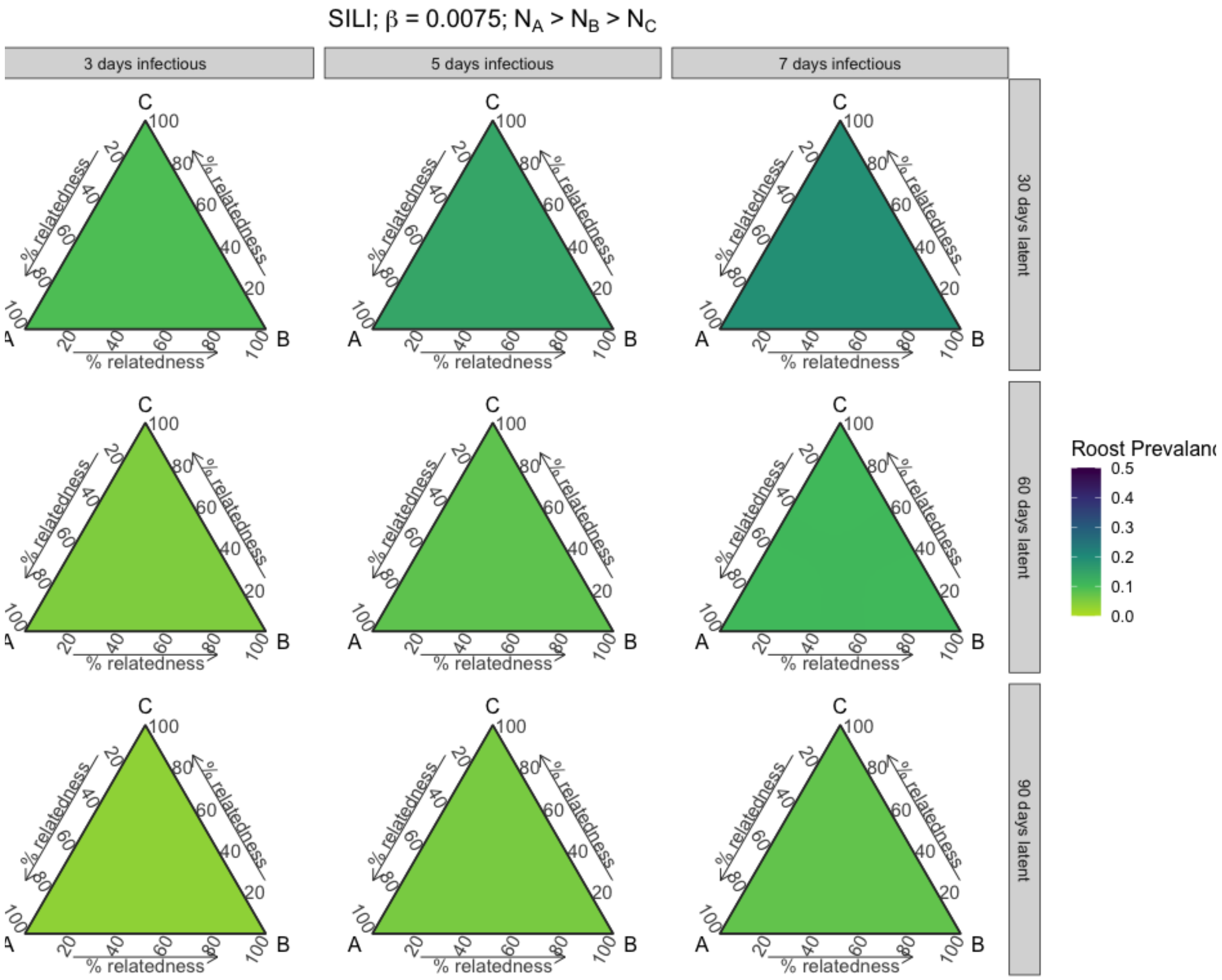


**Figure S52.** Roost prevalence dynamics vary with phylogenetic relatedness between three host species at equilibrium. Results from three-species SILI models across parameter space for varying short infectious and waning immunity periods, when species starting populations sequentially decrease, and when intraspecific transmission (β) is 0.0075.

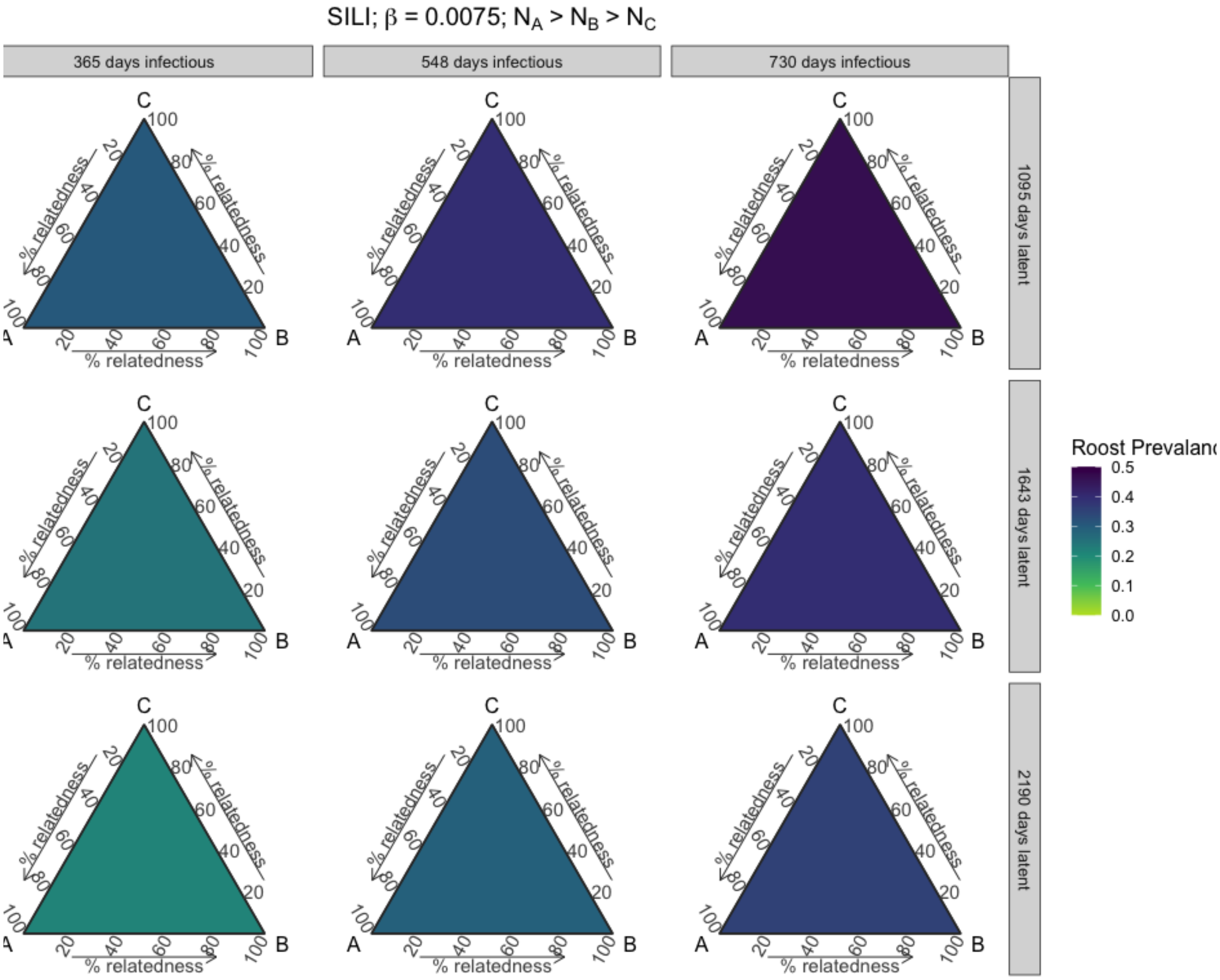


**Figure S53.** Roost prevalence dynamics vary with phylogenetic relatedness between three host species at equilibrium. Results from three-species SILI models across parameter space for varying long infectious and waning immunity periods, when species starting populations sequentially decrease, and when intraspecific transmission (β) is 0.0075.

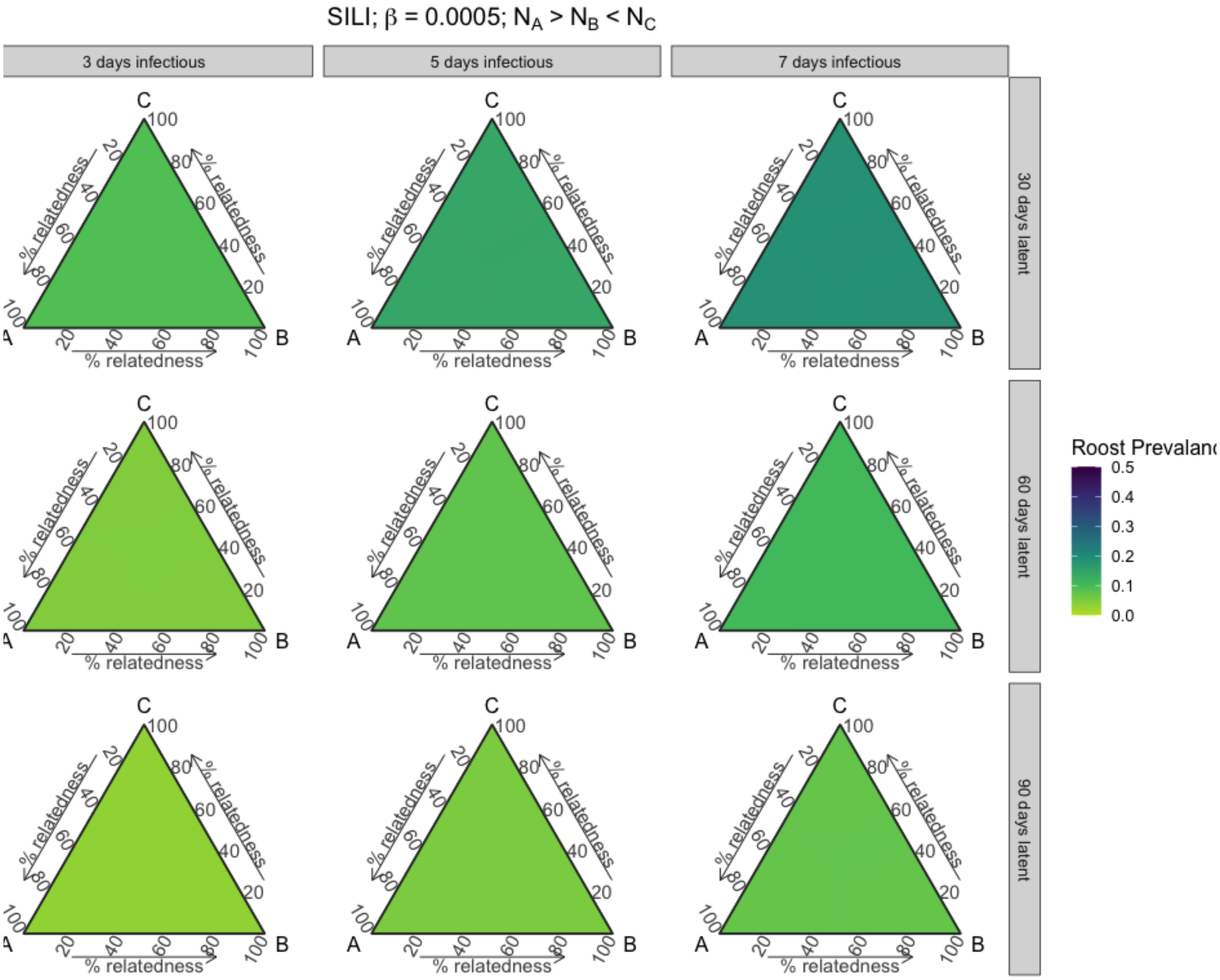


**Figure S54.** Roost prevalence dynamics vary with phylogenetic relatedness between three host species at equilibrium. Results from three-species SILI models across parameter space for varying short infectious and waning immunity periods, when starting populations of species A is greater than both species B and C and species C is greater than B, and when intraspecific transmission (β) is 0.0005.

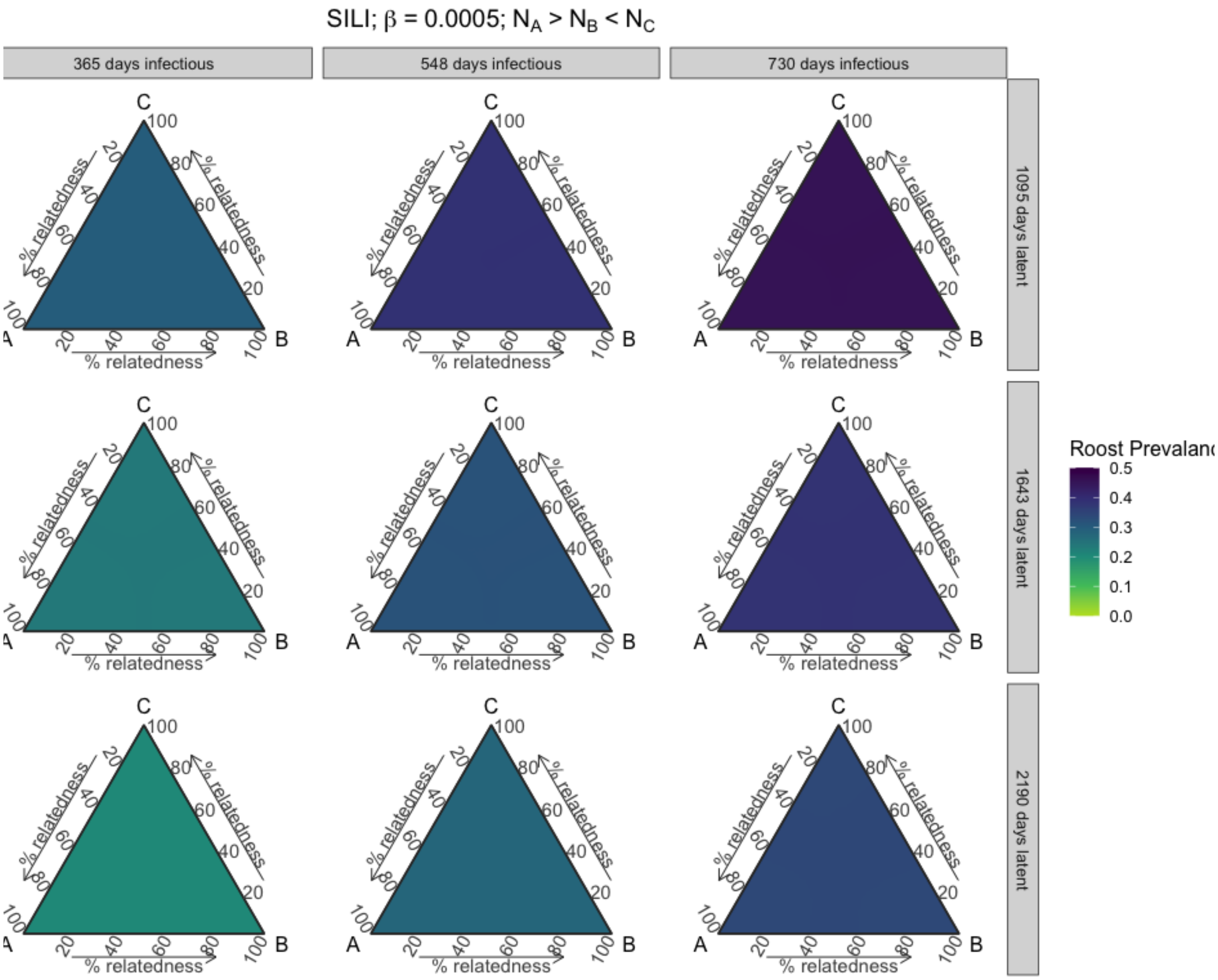


**Figure S55.** Roost prevalence dynamics vary with phylogenetic relatedness between three host species at equilibrium. Results from three-species SILI models across parameter space for varying long infectious and waning immunity periods, when starting populations of species A is greater than both species B and C and species C is greater than B, and when intraspecific transmission (β) is 0.0005.

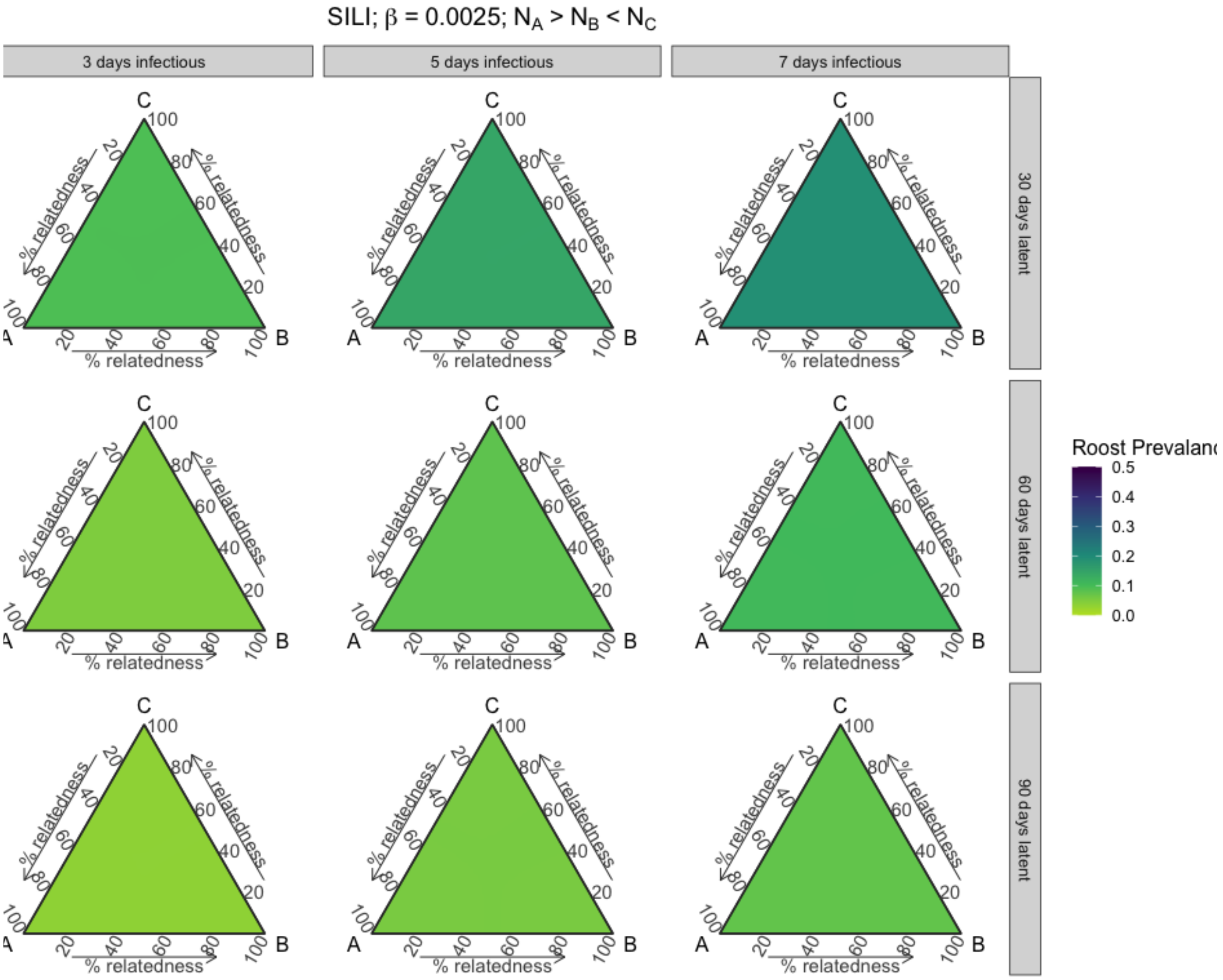


**Figure S56.** Roost prevalence dynamics vary with phylogenetic relatedness between three host species at equilibrium. Results from three-species SILI models across parameter space for varying short infectious and waning immunity periods, when starting populations of species A is greater than both species B and C and species C is greater than B, and when intraspecific transmission (β) is 0.0025.

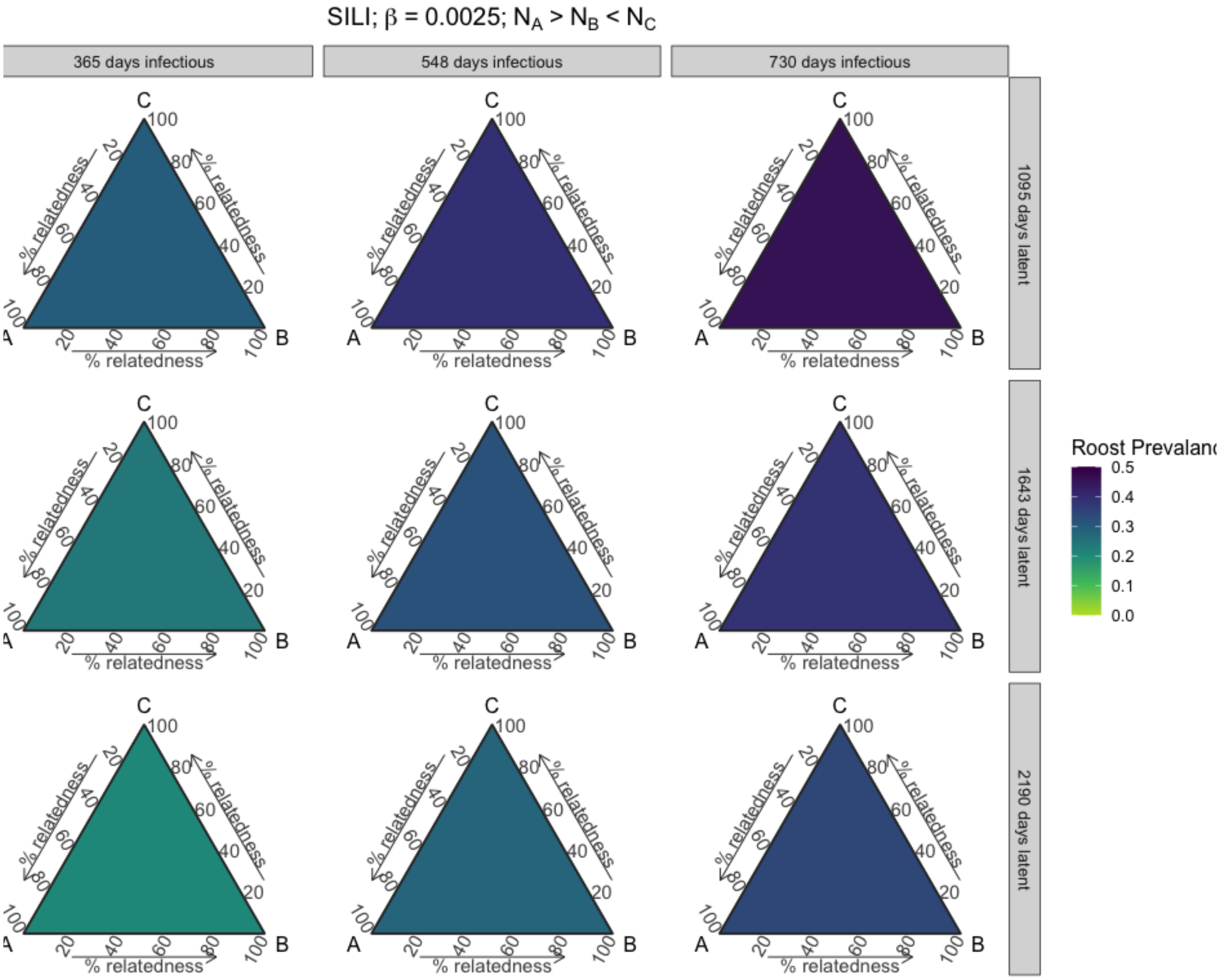


**Figure S57.** Roost prevalence dynamics vary with phylogenetic relatedness between three host species at equilibrium. Results from three-species SILI models across parameter space for varying long infectious and waning immunity periods, when starting populations of species A is greater than both species B and C and species C is greater than B, and when intraspecific transmission (β) is 0.0025.

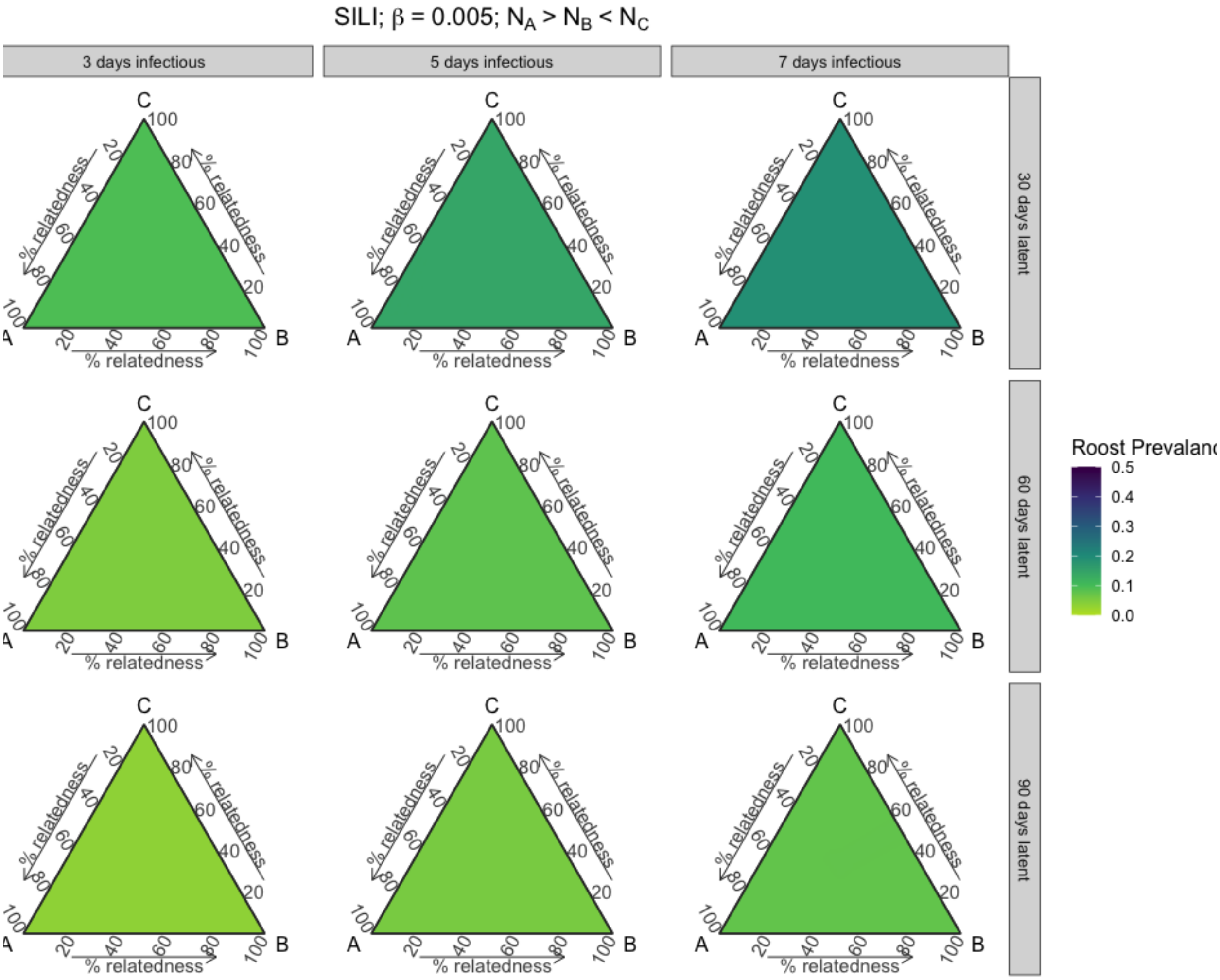


**Figure S58.** Roost prevalence dynamics vary with phylogenetic relatedness between three host species at equilibrium. Results from three-species SILI models across parameter space for varying short infectious and waning immunity periods, when starting populations of species A is greater than both species B and C and species C is greater than B, and when intraspecific transmission (β) is 0.005.

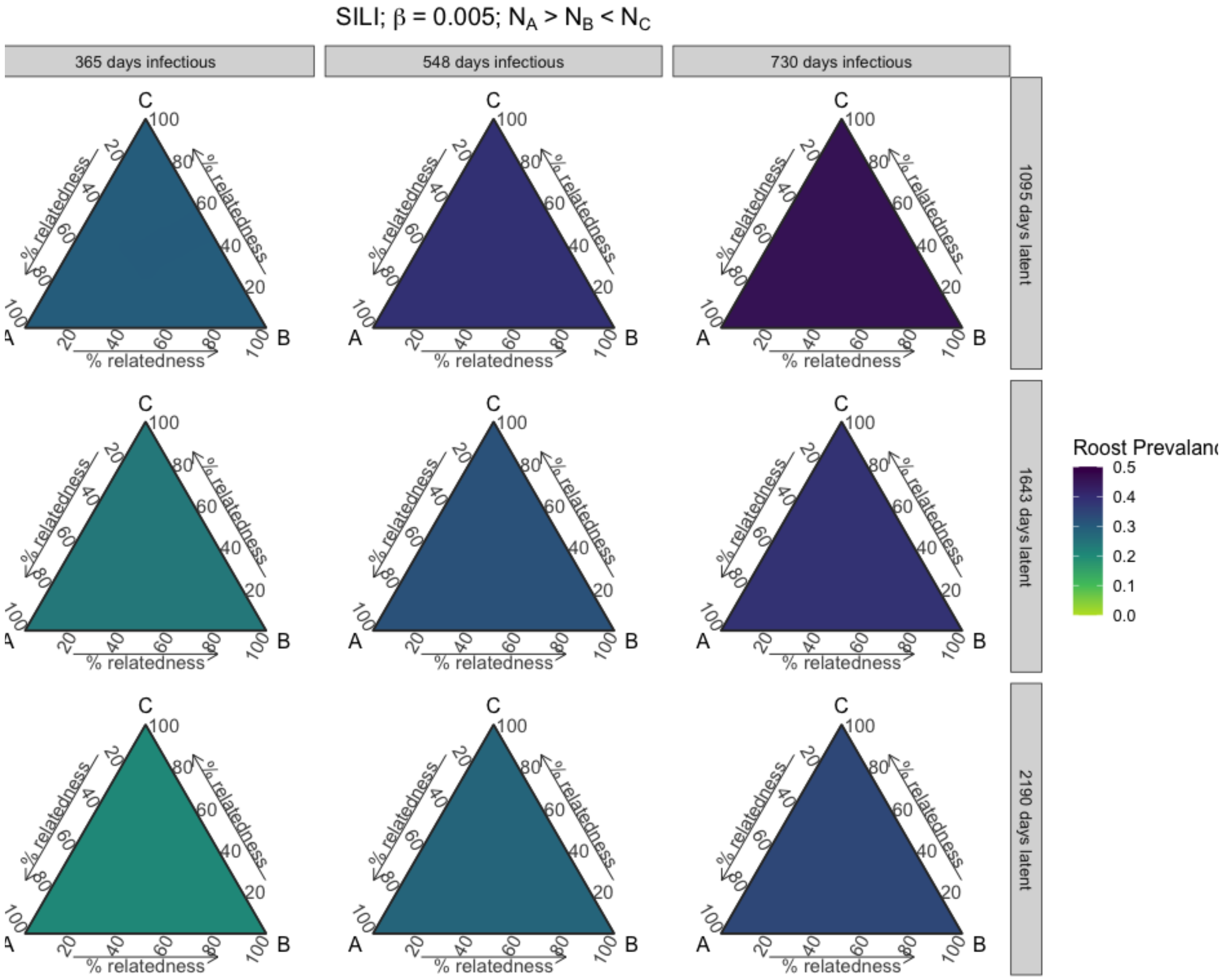


**Figure S59.** Roost prevalence dynamics vary with phylogenetic relatedness between three host species at equilibrium. Results from three-species SILI models across parameter space for varying long infectious and waning immunity periods, when starting populations of species A is greater than both species B and C and species C is greater than B, and when intraspecific transmission (β) is 0.005.

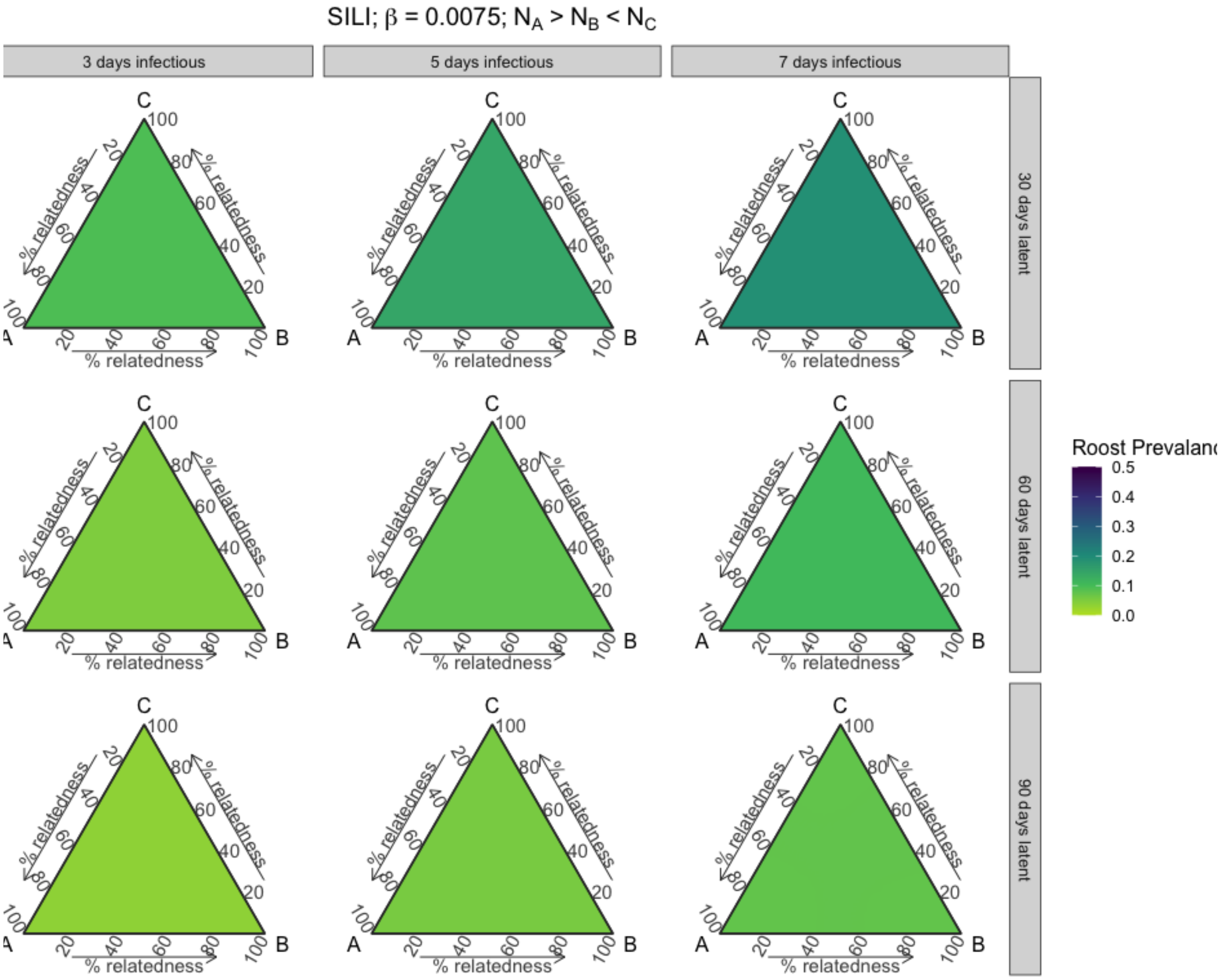


**Figure S60.** Roost prevalence dynamics vary with phylogenetic relatedness between three host species at equilibrium. Results from three-species SILI models across parameter space for varying short infectious and waning immunity periods, when starting populations of species A is greater than both species B and C and species C is greater than B, and when intraspecific transmission (β) is 0.0075.

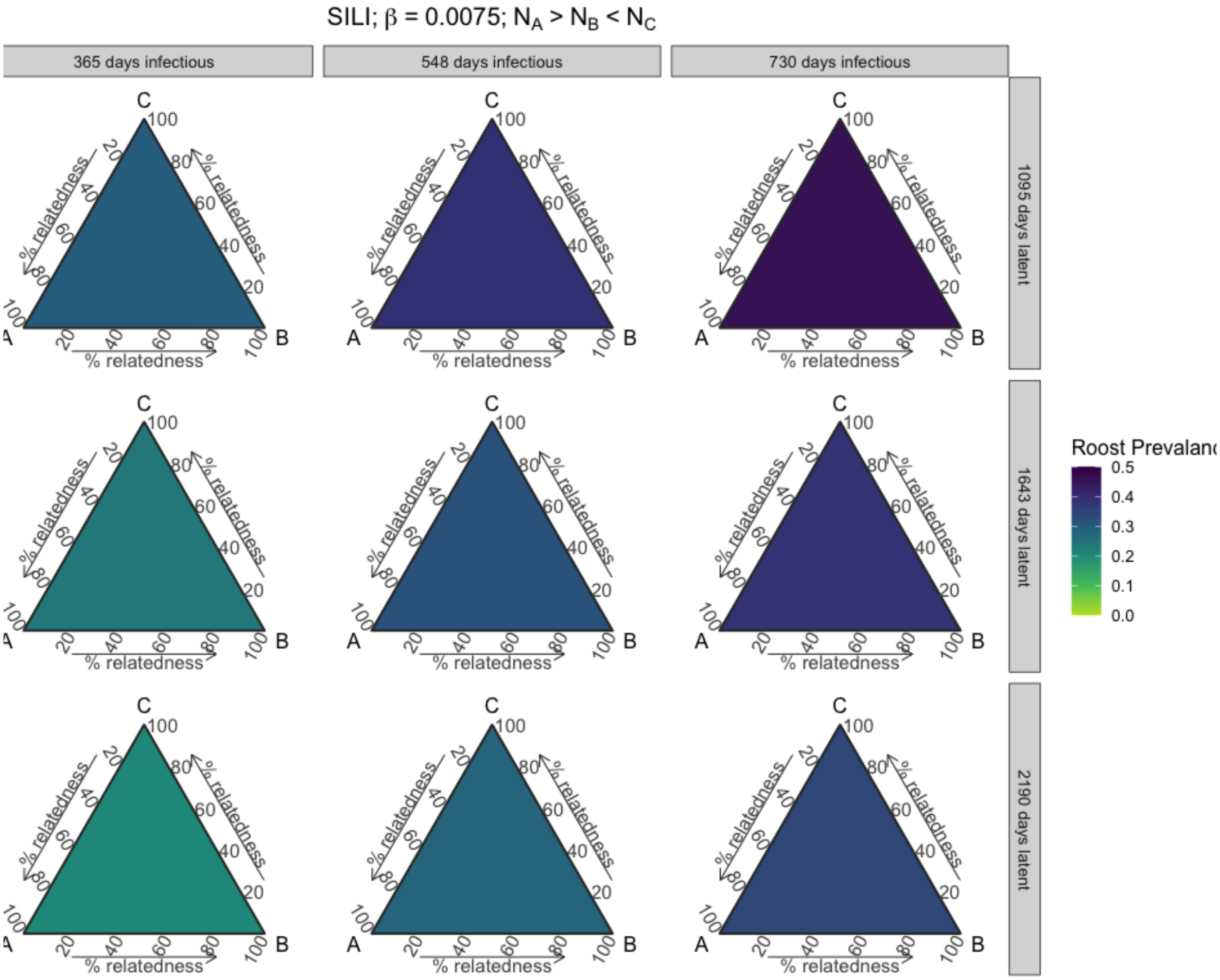


**Figure S61.** Roost prevalence dynamics vary with phylogenetic relatedness between three host species at equilibrium. Results from three-species SILI models across parameter space for varying long infectious and waning immunity periods, when starting populations of species A is greater than both species B and C and species C is greater than B, and when intraspecific transmission (β) is 0.0075.
